# Using GC content to compare recombination patterns on the sex chromosomes and autosomes of the guppy, *Poecilia reticulata*, and its close outgroup species

**DOI:** 10.1101/2020.05.22.110635

**Authors:** Deborah Charlesworth, Yexin Zhang, Roberta Bergero, Chay Graham, Jim Gardner, Lengxob Yong

**Affiliations:** University of Cambridge, Department of Biochemistry, Sanger Building, 80 Tennis Ct Rd, Cambridge CB2 1GA, UK; Centre for Ecology and Conservation, University of Exeter, Penryn, Falmouth, Cornwall, TR10 9FE, UK; Institute of Evolutionary Biology, School of Biological Sciences, University of Edinburgh, West Mains Road, EH9 3LF, U.K.

## Abstract

Genetic and physical mapping of the guppy (*P. reticulata*) have shown that recombination patterns differ greatly between males and females. Crossover events occur evenly across the chromosomes in females, but in male meiosis they are restricted to the tip furthest from the centromere of each chromosome, creating very high recombination rates per megabase, similar to the high rates in mammalian sex chromosomes’ pseudo-autosomal regions (PARs). We here used the intronic GC content to indirectly infer the recombination patterns on guppy chromosomes. This is based on evidence that recombination is associated with GC-biased gene conversion, so that genome regions with high recombination rates should be detectable by high GC content. Using intron sequences, which are likely to be under weak selection, we show that almost all guppy chromosomes, including the sex chromosome (LG12) have very high GC values near their assembly ends, suggesting high recombination rates due to strong crossover localisation in male meiosis. Our test does not suggest that the guppy XY pair has stronger crossover localisation than the autosomes, or than the homologous chromosome in a closely related fish, the platyfish (*Xiphophorus maculatus*). We therefore conclude that the guppy XY pair has not recently undergone an evolutionary change to a different recombination pattern, or reduced its crossover rate, but that the guppy evolved Y-linkage due to acquiring a male-determining factor that also conferred the male crossover pattern. The results also identify the centromere ends of guppy chromosomes, which were not determined in the guppy genome assembly.

## Introduction

The guppy, *Poecilia reticulata*, is an important organism for testing when and how recombination suppression between the sex chromosome pair evolved. One hypothesis is that the main cause of the evolution of recombination suppression between members of sex chromosome pairs is sexually antagonistic (SA) selection acting at a gene linked gene to the sex-determining gene, and maintaining a polymorphic state; the resulting two-locus polymorphism creates a selective force for closer linkage between the sex-determining locus and the gene with the SA polymorphism (Charlesworth and Charlesworth 1980; Bull 1983; Rice 1987). The guppy seems ideal for investigating this hypothesis, because sexually antagonistic male coloration polymorphisms are present in natural populations of the species; these are inferred to benefit males during mating while harming both sexes by increasing predation (Haskins, et al. 1961; Lindholm and Breden 2002). Moreover, the guppy sex chromosomes probably evolved recently, since the homologous chromosome is autosomal in outgroup species (see below). Suppressed recombination could therefore be evolving currently (Wright, et al. 2017; Bergero, et al. 2019; Gordon, et al. 2017)).

An alternative to the SA polymorphism hypothesis is that strong crossover localisation in guppy male meiosis is an ancestral state, and the appearance of a male-determining factor on a chromosome (the guppy LG12) led to instant isolation of most of this chromosome from its X chromosomal counterpart. This alternative can explain findings in European tree frogs (Berset-Brändli, et al. 2008). In the guppy, it is consistent with genetic mapping results that revealed sex differences in recombination patterns in three of the 22 autosomes, as well as the XY pair in guppies, with strong localisation of crossovers to terminal regions in males; LG9 and LG18 show a sex difference in crossover patterns similar to that seen for the XY pair, and results from LG1 were also consistent (Bergero, et al. 2019). Cytological data using MLH1 foci in testis cells showed that crossovers were highly localised to the chromosome termini in male meiosis, and is not specific to the sex chromosome pair (Lisachov, et al. 2015), though such experiments do not provide information about female meiosis. It is plausible that a sexually dimorphic crossover pattern could represent an ancestral state, as crossovers tend to be localised at the terminal regions of chromosomes in male meiosis in several organisms. With the development of molecular markers and genome sequences, it is becoming increasingly possible to estimate genetic maps and compare them with physical maps, and describe species’ recombination patterns and any sexual dimorphism that may exist in these patterns (Hinch, et al. 2014). Sexually dimorphic crossover patterns (or “heterochiasmy”), with crossover events on many chromosomes being more localised in male than female meiosis, have recently been described in fish, including the threespine stickleback (Sardell and Kirkpatrick 2019) and fugu (Kai, et al. 2011).

To test between these hypotheses about crossover localisation in the guppy, two approaches are possible. First, one can test whether crossover localisation in male meiosis is stronger on the sex chromosome pair, compared with the autosomes, and, second, one can test whether localisation is stronger in the guppy, compared with the pattern on homologous chromosomes of suitable outgroup species. Genome-wide heterochiasmy does not exclude the possibility that suppressed XY recombination evolved to prevent crossing over between the male-determining region and SA male coloration factors, as it is possible that the changes that achieved the present XY crossover pattern affected all the chromosomes. The second type of comparison, with closely related species, can test whether this is likely.

Here, we used high-throughput genetic mapping and analyses of genome sequence data to

1. confirm that crossovers are highly localised in physically small terminal regions of all the guppy chromosomes,
2. test whether localisation is stronger in the guppy pseudo-autosomal region (PAR) of partial sex-linkage, compared with other chromosome 12 regions, and whether all guppy autosomes show this pattern, and
3. compare localisation patterns with those in two outgroup species whose genome sequences are available, *Poecilia picta* and *Xiphophorus maculatus* (the platyfish), to test whether the localisation is stronger in the guppy. If not, this would suggest that the pattern of crossovers in the guppy is an ancestral state, and has not evolved in response to this species having evolved an XY sex chromosome pair between which recombination is very rare.

The guppy, *Poecilia reticulata*, has 23 chromosomes, making it difficult estimate male and female crossover patterns on all chromosomes. Closely related species that are suitable as outgroups have similar numbers. Rather than attempting to estimate genetic maps for so many chromosomes from multiple species, we studied a genomic signal that can detect genome regions in which the sequences are subject to unusually high rates of recombination, the GC content. Recombination is accompanied by gene conversion. Studies of sequence changes in crossover and non-crossover meiotic products in mammals (Arbeithuber, et al. 2015) show that recombination-related gene conversion in heterozygotes is biased towards G and C alleles (this process is known as GC-biased gene conversion, abbreviated to gBGC). Any such bias will cause the GC content at weakly selected or neutral sites in genomes to be positively correlated with local recombination rates, and such correlations have been detected for intron sequences and 3^rd^ codon positions in coding sequences in numerous species, including humans and other mammals, birds, rhabditid worms and insects, and plants (Galtier, et al. 2001; Marais and Galtier 2003; Galtier 2004; Galtier, et al. 2009; Lesecque, et al. 2013; Liu, et al. 2018). In yeast, *Saccharomyces cerevisiae* (Birdsell 2002), the correlation is complicated by mutational bias acting to cause sequence changes in the direction opposite to those due to and gBGC, and the rarity of heterozygotes in populations, and its effects may be slight at the genome-wide level (Harrison and Charlesworth 2011; Liu, et al. 2018).

Regions with extremely high crossover rates in one sex, have large GC differences from the rest of the genome, based on yeast recombination hotspots (Mieczkowski, et al. 2006) and physically small pseudo-autosomal regions (PARs) of sex chromosomes (Rouyer, et al. 1986; Hinch, et al. 2014). Most strikingly, in the mouse, a gene has been found that spans the PAR boundary (i.e. part of it is in the distal chromosome region of homology between X and Y chromosomes, while the rest of the gene is fully sex-linked). The exons and introns from the PAR part are more GC-rich than the fully sex-linked part of the gene (Marais and Galtier 2003). Rather than being an evolutionary consequence of local crossover rates, such correlations might instead reflect a process by which GC content controls crossover rates (Mieczkowski, et al. 2006; Liu, et al. 2018), or a tendency for GC-rich sequences, including CpG islands to evolve in highly recombinogenic genome regions (Han, et al. 2008). However, a region of moderate GC content that was recently transposed into the mouse PAR has been found to experience very frequent recombination, indicating that the recombination rate is not, at least in this case, a consequence of high GC content (Morgan, et al. 2019). If the recombination rate is very high in a region, even in just one of the two sexes, it can therefore lead to high GC content, and this evolutionary consequence will be detectable in sequence data from both sexes. To gain an understanding of recombination rate patterns, and differences in these patterns between different guppy chromosomes and sex chromosome regions, and between these and the patterns in related fish species, we therefore analysed GC content in the guppy, and homologous sequences of related species. To avoid, as far as possible, other factors that can influence GC content, we focused on the GC content of introns of guppy genes.

Recombination is also mutagenic (Lercher and Hurst 2002; Arbeithuber, et al. 2015; Halldorsson, et al. 2019; Kessler, et al. 2020). This effect leads to the further consequence of higher numbers of substitutions per site in high-recombination regions, compared with the rest of the genome, and the GC-biased region of the mouse gene that spans the PAR boundary gene indeed also evolves much faster than the fully sex-linked part (Galtier 2004). Such a substitution rate difference is also detected in pseudo-autosomal genes of the human and ape sex chromosomes (Filatov and Gerrard 2003), and in birds (Rousselle, et al. 2018). We therefore also analysed substitutions between guppy genes, and their orthologues in related species, and compared the rates between regions with local very high recombination rates and other genome regions.

## Results

### Chromosome homologies in the species studied

The analysis described in the Methods section identifies the platyfish homologues of most guppy chromosomes, which mostly consist almost exclusively of sequences from single platyfish chromosomes, with small contributions from other chromosomes (see the folder “GUPPY-platy chromosome homologies” under Dryad **doi number to be added**), and Supplementary Table S1 summarizes the homologies for the 20 guppy chromosomes where they were clear-cut. Seven guppy chromosome assemblies appear to include substantial portions from more than a single platyfish chromosome. Excluding the guppy chromosome 2, which is known to be a fusion between two chromosomes that are separate in *X. maculatus* (Künstner, et al. 2017), the other such chromosomes are the guppy LGs 6, 8, 9, 14 and 19, and LG16 has a small anomaly (see the Dryad folder “GUPPY-platy chromosome homologies”). For example, LG9 largely corresponds to the platyfish chromosome 12, but a region of about 14% of the 34 Mb chromosome (around 5 Mb, near the zero end of the assembly) is assembled on the platyfish chromosome 11 (Supplementary Figure S1). These instances may be due to assembly errors, or could be genuine genomic rearrangements (some errors are detected in our genetic mapping, see Supplementary Figures S7 and S9, also unpublished results of Charlesworth et al. 2020 and Fraser et al. 2020). Assembly error could account for the result for the guppy chromosome 7: the large region at the right possibly assembled in an inverted order (see the Dryad folder “PNG+TRACE_G (dot plots)”) may explain why, in the family where the largest number of SNPs can be mapped (LAH), a set of SNPs in the middle of the assembly co-segregate, and the SNPs on both sides of them are assigned to recombining regions. The homology is, however, clear for the guppy XY pair, LG12, which is homologous to chromosome 8 of the platyfish and to chromosome 12 of *P. picta*.

### GC content of first introns versus other introns

Before testing for a relationship between GC content and physical position, we first established that the values for first introns do not differ from those of other introns (Supplementary Table S1). First introns of genes are often under stronger selective constraint than other introns. As reviewed by Park (2014), the most 5’ introns have distinctive properties in several organisms, and a class of “intron-mediated enhancers” regulating gene expression level are known, are located unexpectedly often within mouse first introns; first introns also tend to be longer than other introns and have the greatest sequence conservation in several taxa, including *Drosophila* (Marais, et al. 2005) and humans (Park, et al. 2014). Strong purifying selection might therefore act, and might maintain low GC values, even in genome regions with high rates of recombination.

There was no significant for any guppy chromosome (Wilcoxon signed-rank test, p > 0.1) apart from a small difference for LG10 (median of first introns = 0.372, versus 0.368 for other introns, Wilcoxon signed-rank test, p < 0.0005). For the platyfish, 23 chromosomes had non-significant results, while a small difference was detected for chromosome 23 (the medians for first and other introns were 0.364 and 0.362, respectively; this is not significant after False Discovery Rate correction). The results described below are therefore based on the GC values for all introns of each gene.

### Change-point analysis of intron GC content and determination of centromere positions in the guppy and the platyfish

We used change-point analysis (see Methods) to test which chromosomes show significant changes in GC content, and to estimate the positions where the changes occur, and their magnitudes. All but four guppy chromosomes, and all but three in the platyfish, had a prominent spike in GC content at one end of the chromosome (Supplementary Figure S2; Supplementary Figure S3 shows the relationships of the species studied).

The centromere regions were not identified in the published genome assemblies of either the guppy (Kunstner *et al.*, 2016) or the platyfish, but can be identified based on the GC analysis Assuming that the spikes in intronic GC values indicate terminal recombining regions of these acrocentric chromosomes, the other ends should represent the centromeres (LG2 showed a spike in the middle of the chromosome, perhaps indicating a signal retained from the pre-fusion period of its evolution). Among the 22 other guppy chromosomes, 19 have clear terminal spikes, 5 at the start of the assembly, and 15 at the end (Table 1); only a few chromosomes have no large change. In the 21 platyfish chromosomes with clear spikes, 15 had elevated CG at the start, versus 6 at the end of the assembly (Supplementary Table S2). For 17 chromosomes, the same end was identified as the centromere end in both species, taking account of the fact that, for some chromosomes, the genes assigned the lowest assembly positions in the platyfish are homologous to genes with the highest positions in the guppy assembly (see Supplementary Table S2).

**Table 1.**
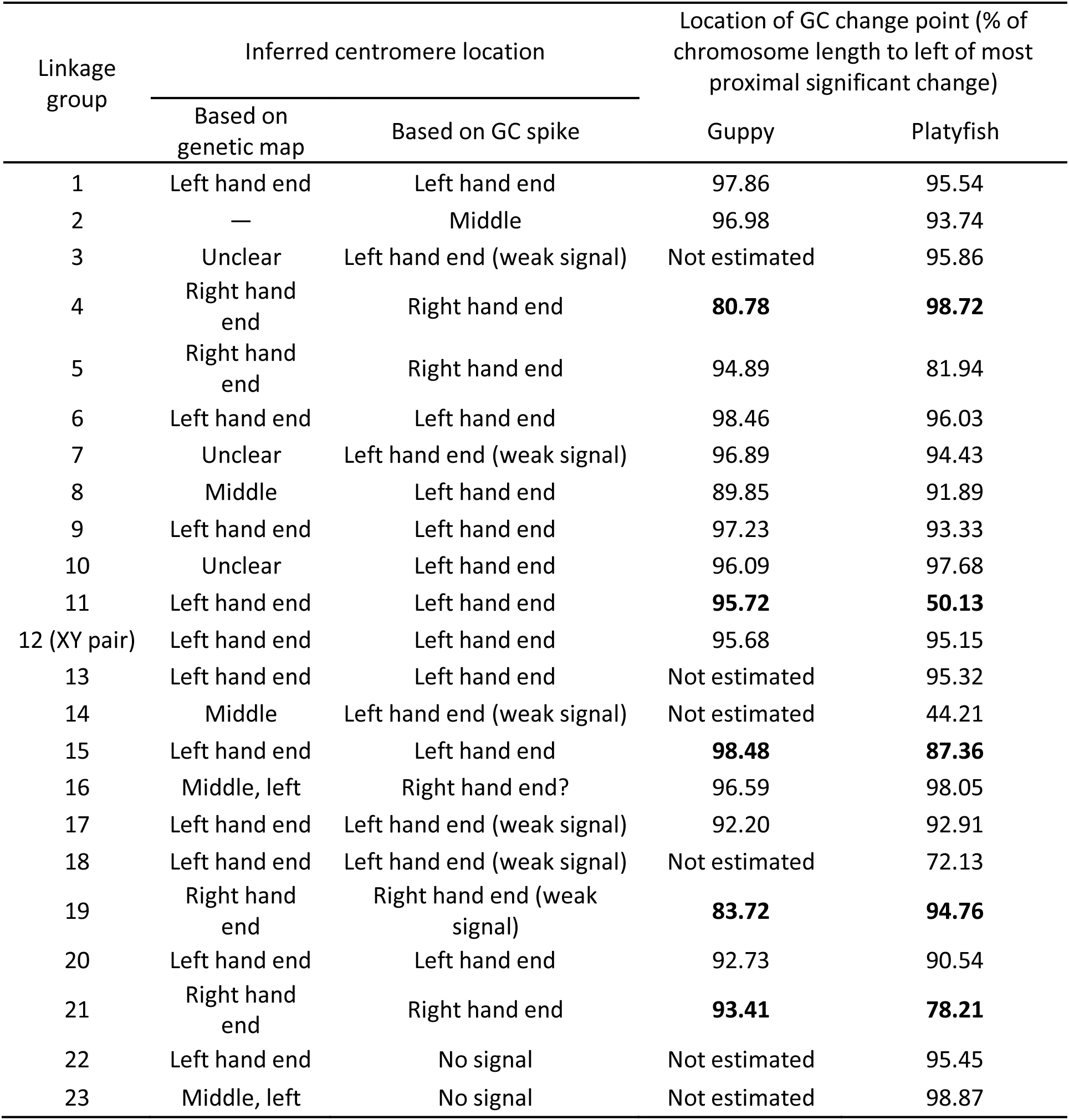
Inferred centromere ends of guppy chromosomes, based on GC_intron_ spikes at the opposite chromosome ends, implying that those ends had high levels of recombination, and therefore cannot be the centromere ends. The table also shows the location of statistically significant GC change points in the guppy and platyfish; to enable the comparison, these are shown as percentages of the chromosome length, because the estimated chromosome sizes differ slightly in the two species. Chromosomes showing large differences in change-point locations between the species are shown in bold font in the right-hand two columns.

| Linkage group | Inferred centromere location |  | Location of GC change point (% of chromosome length to left of most proximal significant change) |  |
| --- | --- | --- | --- | --- |
|  | Based on genetic map | Based on GC spike | Guppy | Platyfish |
| 1 | Left hand end | Left hand end | 97.86 | 95.54 |
| 2 | — | Middle | 96.98 | 93.74 |
| 3 | Unclear | Left hand end (weak signal) | Not estimated | 95.86 |
| 4 | Right hand end | Right hand end | **80.78** | **98.72** |
| 5 | Right hand end | Right hand end | 94.89 | 81.94 |
| 6 | Left hand end | Left hand end | 98.46 | 96.03 |
| 7 | Unclear | Left hand end (weak signal) | 96.89 | 94.43 |
| 8 | Middle | Left hand end | 89.85 | 91.89 |
| 9 | Left hand end | Left hand end | 97.23 | 93.33 |
| 10 | Unclear | Left hand end | 96.09 | 97.68 |
| 11 | Left hand end | Left hand end | **95.72** | **50.13** |
| 12 (XY pair) | Left hand end | Left hand end | 95.68 | 95.15 |
| 13 | Left hand end | Left hand end | Not estimated | 95.32 |
| 14 | Middle | Left hand end (weak signal) | Not estimated | 44.21 |
| 15 | Left hand end | Left hand end | **98.48** | **87.36** |
| 16 | Middle, left | Right hand end? | 96.59 | 98.05 |
| 17 | Left hand end | Left hand end (weak signal) | 92.20 | 92.91 |
| 18 | Left hand end | Left hand end (weak signal) | Not estimated | 72.13 |
| 19 | Right hand end | Right hand end (weak signal) | **83.72** | **94.76** |
| 20 | Left hand end | Left hand end | 92.73 | 90.54 |
| 21 | Right hand end | Right hand end | **93.41** | **78.21** |
| 22 | Left hand end | No signal | Not estimated | 95.45 |
| 23 | Middle, left | No signal | Not estimated | 98.87 |

In the guppy, significant changes in GC_intron_ values were detected for 17 chromosomes (Table 1). For all except three of these (LGs 4, 8 and 19), the region of elevated GC occupied less than 8% of the chromosome’s total length. Chromosomes 4 and 8, had high GC in 19 and 10% of the chromosome, respectively, and the changepoint is uncertain for LG19, where the signal was weak. Overall, therefore, most guppy chromosomes show signals consistent with strongly localized recombination in one or both sexes. There is no clear difference between the guppy sex chromosome, LG12, and the autosomes. The high GC identified by our change-point analysis for LG12 occupies the terminal 4.3% of this chromosome (Figure 1), similar to the results for several other chromosomes (see Figure 4 below); overall, 7 guppy autosomes have smaller regions of high GC_intron_, and 10 have larger regions (Table 1). The total assembled sequence length of LG12 is 26.4 Mb (Kunstner *et al.*, 2016), and the location of its major change in GC content indicates that, consistent with our previous genetic mapping results (Bergero, et al. 2019), only about 1.15 Mb of the terminal end of the chromosome undergoes crossovers.

**Figure 1.**
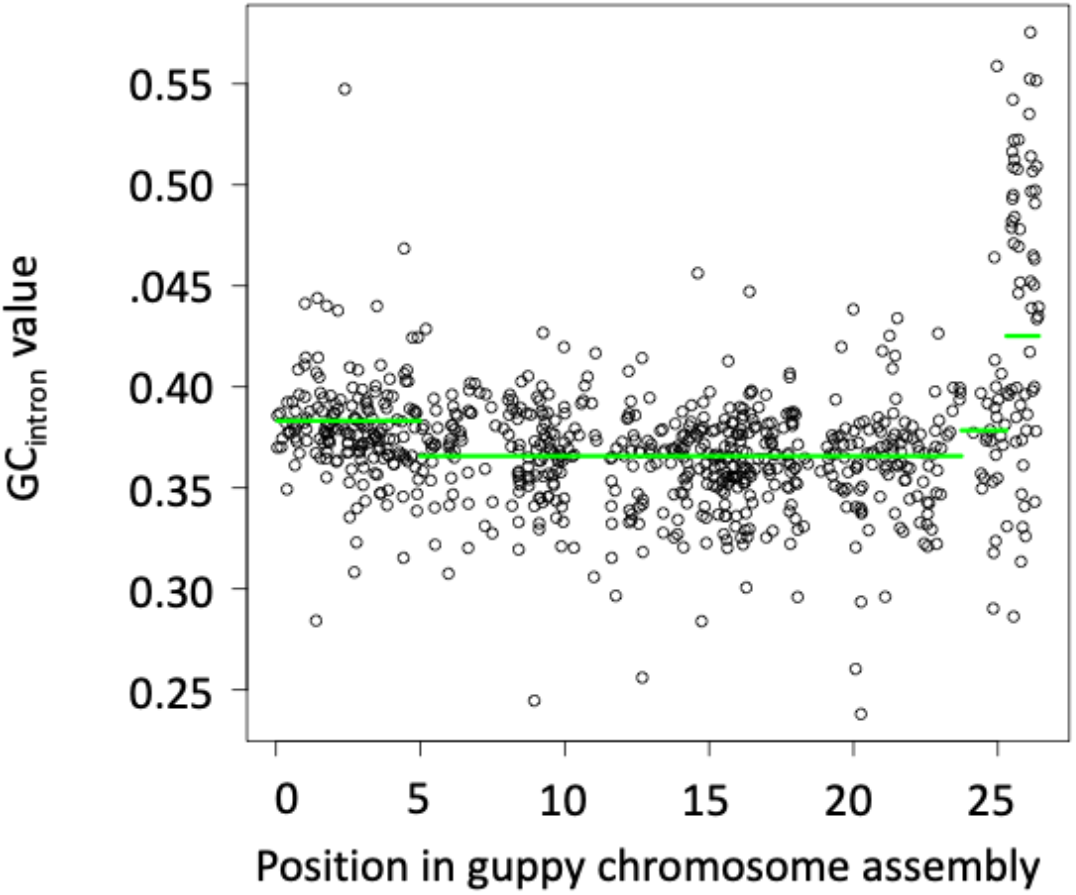
GC content of the guppy X chromosome. The circle symbols show the GC values for the introns of individual genes, and green horizontal lines show the change-points detected by our analysis (see text), with a large change detected near the right-hand end of the chromosome, and two small changes in other locations.

### Gene density

We tested the possibility that GC differences might be related to local gene densities, since coding sequences have higher GC than introns. In the human genome, the GC content is positively correlated with gene density (Payseur and Nachman 2002). However, we found no such effect in the guppy introns (Supplementary Table S3); the table shows that only 6 guppy chromosomes show significant relationships, and these all indicate negative, not positive correlations. The high GC content at the chromosome termini in the guppy therefore does not appear to be caused by an enrichment of coding sequences.

### Analyses of P. picta introns

We next examined intron sequences in *P. picta*, the closest relative of the guppy (Supplementary Figure S3), using low coverage sequencing data. Our analysis used the female with the highest coverage (about 13-fold, see Methods section). Although the intron GC content patterns are much less clear than in the guppy and platyfish, elevated values are clearly visible at one end of many the chromosomes (Supplementary Figure S4), and agree very well with those in the guppy, including identifying the same four or five chromosomes that showed no terminal GC spikes, guppy LGs 3, 14, 17, 22 and 23 (Supplementary Figure S2, and Table S2; the full results for *P. picta* are shown in a folder named “Picta INTRONIC_GC” in Dryad **doi number to be added**). The recovery of a signal when using the guppy as a reference for assembling/analysing reads from this species suggests that the signal is unlikely to be specific to the guppy lineage. We did not attempt change-point analyses, but tested whether this species also has statistically significant changes by Wilcoxon signed rank tests for each chromosome, comparing the median intronic GC proportion between the tips where GC spikes were detected in the guppy, and the remainder of the chromosome. Significant one-tailed test results were obtained when the tips of each chromosome were defined by various different percentage cut-off values; for the terminal 10%, P = 0.00013, for 5% or 2% it was 0.0011, and results for the means were similar.

### GC in coding sequences

A less clear pattern is expected in coding regions than for intronic GC, given that selection on codon usage may affect such sites (Labella, et al. 2019), and also because GC content is generally higher in coding regions than introns, as reviewed by Beauclair (2019), making increases less likely to be detected. Nevertheless, GC in third codon positions (GC3 values) also show significant increases at the ends of many guppy chromosomes. Change-points were detected only at the right-hand ends of seven LGs (1, 6, 8, 10, 11, 13 and 15), and the left-hand ends of the three LGs 16, 19 and 21; these all agree with the chromosome end with the GC_intron_ spike. Change-point analysis detects a slightly, but significantly, higher value at the tip of LG12 (Figure 2); figures for the other chromosomes, many of which show clearer signals of elevated GC3, are in files “GC3 CPA guppy” and “GC3 CPA platyfish” in Dryad).

**Figure 2.**
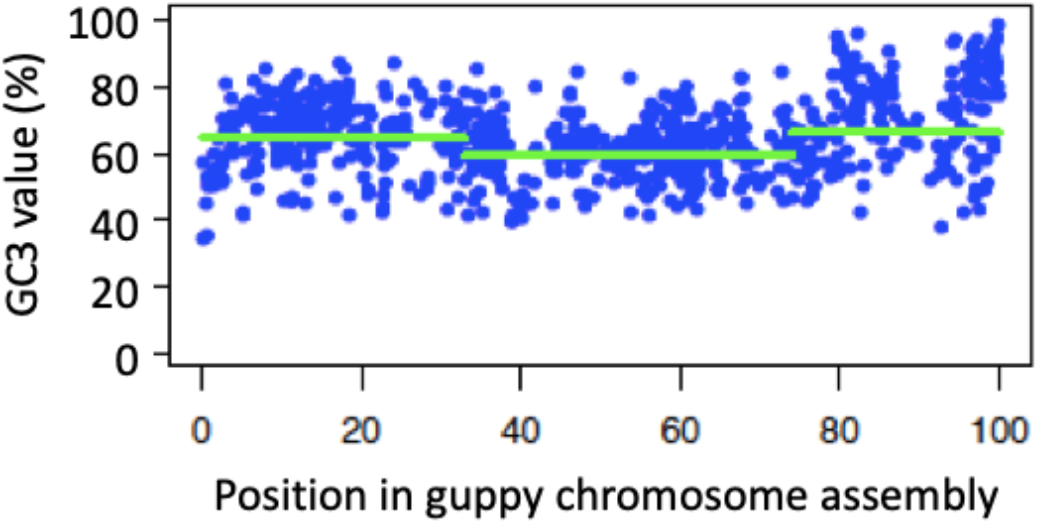
GC3 content of guppy X-linked coding sequences. The figure shows the GC3 value for each gene plotted against its position along the chromosome for guppy sex chromosomes (LG12).

### Comparisons with change-points in the platyfish

In the platyfish, all 24 chromosomes have detectable change points, unlike the guppy result (Supplementary Table S4A). Excluding chromosomes such as LG14, which also has too weak a signal in the guppy to estimate a change-point, see Table 1), we can compare the change-point locations in 18 chromosome homologues of these two species (Figure 3). When a large increase in GC_intron_ values is detected in either species, it is mainly near the end of a chromosome (Figure 4). The chromosome ends also show an excess of decreases in GC (three guppy chromosomes have such decreases exceeding 5%, and 5 platyfish chromosomes, see Supplementary Table S4B). Although it is likely that these regions include more assembly errors than other parts of the chromosomes, this seems unlikely to account for this effect.

**Figure 3.**
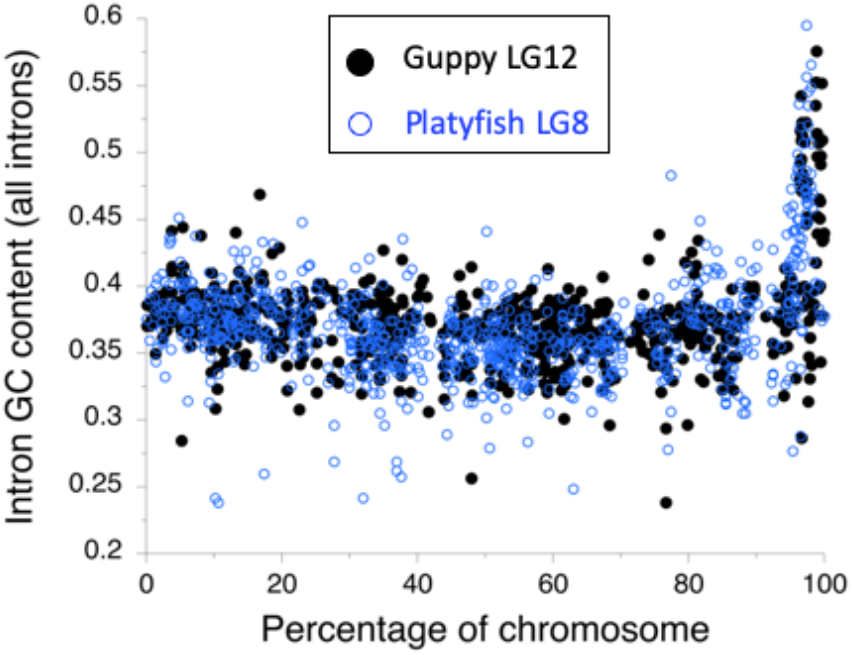
Comparison between GC content of introns in genes of the guppy LG12 (black dots) and the homologous platyfish chromosome (chromosome 8, blue circles).

**Figure 4.**
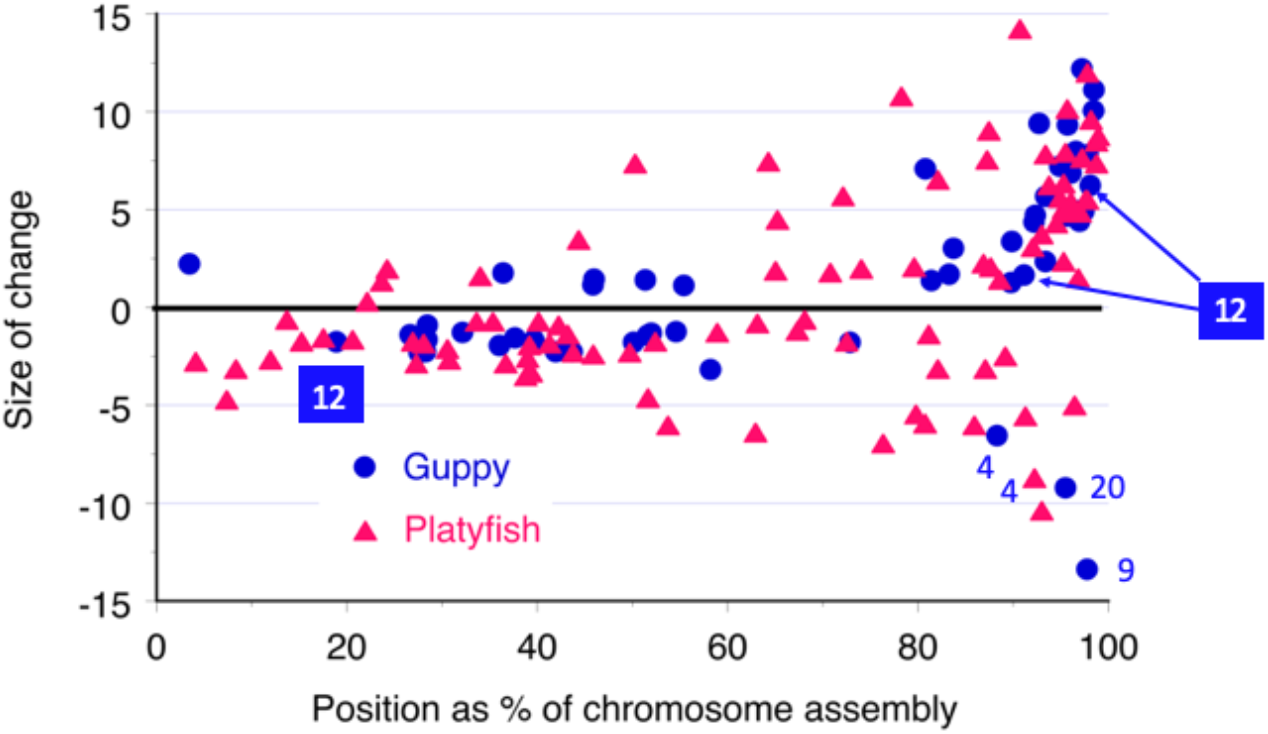
Comparison of intronic GC content change-point sizes for all guppy chromosomes and platyfish (blue circles and red triangles, respectively). The y axis shows the sizes of all changes detected, in either direction (the three changes detected for the sex chromosome pair are indicated). Because the chromosomes are of varied sizes, the x axis shows the locations of all changes detected by our analyses, as percentages of the relevant lengths of the chromosomes, and the chromosomes are oriented with the inferred centromeric end at the left, when this could be determined (see text).

Four guppy chromosomes (LGs 5, 11, 15 and 21) have substantially more proximal change-point locations in the platyfish homologues, while the homolog of the guppy LG4 has a substantially more distal location (though the assemblies of this chromosome in the two species differ too much to be reliably compared, see the folder of dot plots in Dryad). A different change-point analysis, “ruptures” (see Methods) did not change the inferred centromere ends or the chromosomes showing no clear GC spikes (in both species). The estimated change-point locations were also similar, except that LG15 showed a smaller difference between the two species, and eight LGs had change-points substantially more proximal in the guppy than the platyfish. We therefore conclude that the guppy does not show any consistent tendency to have more terminal GC spikes. Proximal changes from low to high GC content are unlikely to indicate regions with very high recombination rates, because counts of MLH1 foci show that guppy chromosomes rarely have more than one crossover per bivalent (Lisachov, et al. 2015), and our genetic maps never greatly exceed 50 cM for any linkage groups in meiosis of males or females (Bergero *et al.*, 2019). They could reflect local recombination hotspots not in the terminal regions of platyfish chromosomes, particularly in the platyfish, as map lengths estimates in *Xiphophorus* are larger than in the guppy (Walter, et al. 2004), suggesting that many chromosomes may often have multiple crossovers, not just one at the terminus (see discussion below).

The difference in change-point locations between the two species was tested by a paired Wilcoxon Rank Sign test of the median change point locations in terms of the percentage of the chromosome occupied by the high GC region. For five chromosomes, two consecutive change points were detected (Supplementary Table S4B). Using the most distal change-point for all chromosomes, the median and mean values for the guppy are 3.3% and 4.8%, versus 4.8% and 8.8% for the platyfish, which is not a significant difference (P = 0.172); using the more proximal of the two consecutive change points for the five chromosomes with two successive change-points, the differences between the species is again non-significant (P = 0.122). Overall, the data from both species strongly suggest terminal localization of recombination in both species, and do not support any major genome-wide increased localization in the guppy. The slightly more extreme crossover localisation in the guppy compared with platyfish is not a significant difference.

### Synonymous divergence between species

To test whether the inferred high recombination regions of guppy chromosomes also show the expected signal of higher mutation rate than other regions of the same chromosomes (see Introduction), we estimated synonymous site divergence (*K*_s_) values between guppy and platyfish coding sequences; intron sequences are less reliably alignable, and were not analysed. Figure 5 shows estimates for LG12, using two different models (see Methods section). The two models that use simple corrections for the saturation expected under high divergence, NG and LPB, gave very similar results, and neither showed clear spikes in divergence at the chromosome ends (Figure 5). With the GY model, however, significantly elevated values are detected at the ends of chromosomes, as expected for a model designed for high divergence (Figure 5; results for all guppy chromosomes are shown in Supplementary Figure S5). One-tailed Wilcoxon tests indicate a highly significant overall tendency for terminal regions of the chromosomes to have high *K*_s_ whether we compare the 10% of the assembly most distant from the inferred centromere, or when comparing smaller terminal regions with the centromere-proximal 95%, 97% or 99% regions; in all cases P < 0.0006 for the medians, and 0.0002 for the mean values. Thus the chromosome ends display the high divergence expected if recombination rates are high and recombination is mutagenic.

**Figure 5.**
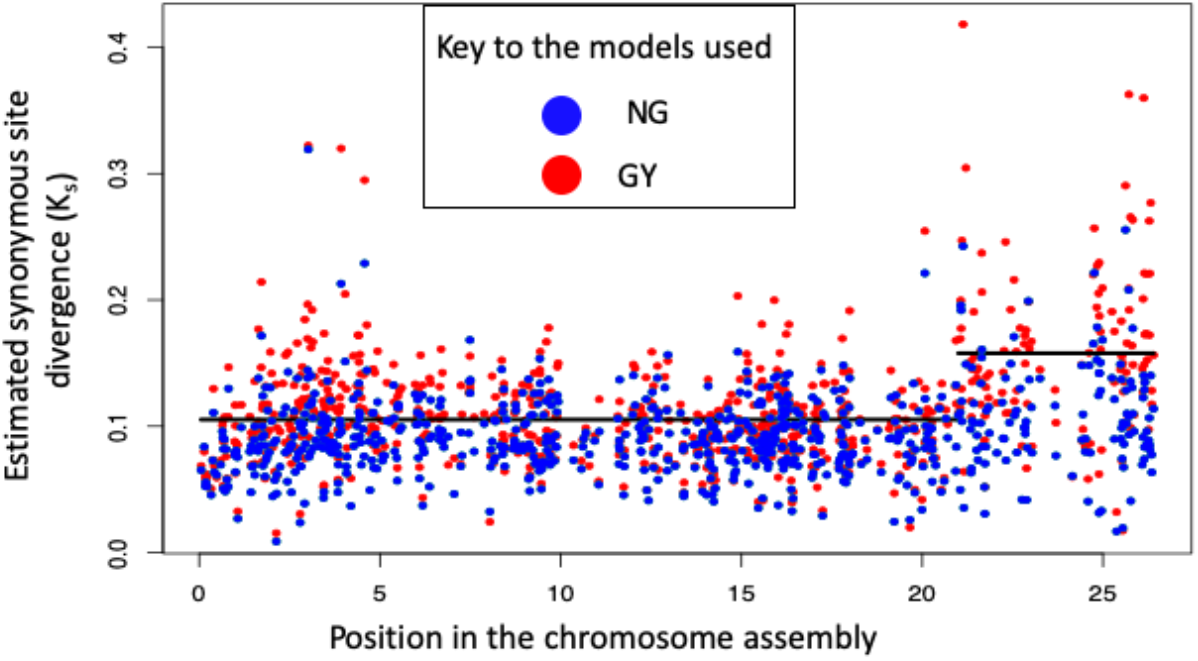
Synonymous site divergence (*K*_s_ values) between the guppy and platyfish for genes on the guppy sex chromosome pair, LG12. The different colours show *K*_s_ values obtained using three different models (see the Methods section; blue represents the LPB and NG models, which are indistinguishable, and red shows the GY estimates. The horizontal lines denote the values inferred by change-point analysis.

### Comparison with high-throughput genetic mapping results for guppy autosomes

To test whether the GC spikes detected on the guppy chromosomes correspond with regions of frequent crossing over, we compared their genetic maps with the GC patterns, using high-throughput SNP genotyping data in guppy families (described in detail in Charlesworth et al. 2020), and examining the results for genic SNPs that are informative in meiosis of the male parent. When the data for a chromosome included such SNPs near both ends of the assembly (Supplementary Figure S6), we recorded the end that showed the crossovers. We used results from female meiosis to check that the regions mapped include genes that belong to the chromosome under study, rather than occasional mis-assembly of a gene that belongs on another chromosome, which would produce spurious signals on crossing over. Other types of assembly errors, such as inversion of a physically large region, are unlikely to create disagreements between the GC spike positions and the regions where crossovers are detected. For example, a highly recombining tip wrongly assembled in the middle of the chromosome should also yield a GC spike in a middle region, which was not seen (Supplementary Figure S2).

Overall, five guppy chromosomes have GC spikes that are weak (LGs 3 and 17) or undetectable (LGs 14, 22 and 23), and we thus have no clear prediction about which end of the guppy chromosome is the centromere, and which end is expected to show recombination events. Such cases could be due to terminal regions being missing from the assemblies of some or all of these chromosomes; however, for most chromosomes few sequences present in the platyfish assembly appear to be missing from the guppy. The exceptions are both termini of LG2, the right-hand ends of guppy LGs 9 and 14, the latter with no GC signal, LG16, and the left-hand end of LG21 (see file “PNG+TRACE_G (dot plots)” in the Dryad folder). Of the 19 chromosomes where GC spikes are detected (even if not significant in change-point tests), 13, including LG12 which is not shown here, show a recombining region at the predicted end, while 6 chromosomes do not (Supplementary Table S2). However, in at least one of the 6 sibships, 6 LGs show apparent recombination in the middle of the assembly, as well as at the expected end (see Supplementary Table S5, Figure S7, and the file “Autosomal segregation (6 fams HT results)” in the Dryad folder). For some of these chromosomes (LGs 5, 7, 8, 10, 16 and 23), crossing over patterns in male meiosis may be similar to those in females.

## Discussion and Conclusions

### Crossover patterns in the guppy and related fish

Our results concerning crossing over are consistent with cytogenetic observations on guppy in testis cells. Crossovers on both autosomes and sex chromosomes show terminal localization (Lisachov, et al. 2015). Consistent with this, genetic mapping results in families from multiple guppy natural populations (Bergero *et al.*, 2019), found that crossing over in male meiosis is very frequent in the terminal 1-2 Mb of the XY chromosome (LG12) most distant from the centromere, but very rare elsewhere, and that several autosomes show similar crossover localization in male meiosis.

Our inference that crossover patterns are similar on most other guppy chromosomes, and in related species, is indirect, using analysis of GC in introns, similar to the analysis of GC in third positions of codons used to infer recombination differences in bird sex chromosomes (Xu, et al. 2019). The basis of our inference is that high rates in meiosis of one (or both) sexes are expected to result in high GC values in the genome sequences of both sexes, allowing us to use data from the published guppy genome assembly from a sequenced female. If regions of high GC content in these fish are consequences of evolution under high recombination rates in male meiosis, like those detected in the LG12 PAR by genetic mapping, the causative high crossover rates should be restricted to males. Such analyses cannot tell us whether the sexes differ or not. These terminal regions could have much lower rates in female meiosis, as our genetic mapping suggests (Bergero, et al. 2019).

Finding high GC in the chromosome regions where high male crossover rates are detected in our genetic maps for the species cannot exclude the possibility that other factors also contribute to elevated GC levels, as reviewed in the Introduction section. However, we found that elevated GC levels at guppy chromosome tips are accompanied by elevated synonymous site substitution rates, as expected if recombination is mutagenic (as also reviewed in the Introduction). This observation suggests that high recombination rates are indeed likely to be involved, rather than these guppy genome regions having special properties that have selected for high GC levels, or unusually high gene densities (which, as explained above, our tests do not find). However, the high GC levels might not be due to GC-biased gene conversion, as it is also possible that high GC directly causes higher recombination rates than occur in genome regions with lower GC levels, as has been demonstrated in experiments in yeast (Kiktev, et al. 2019).

Overall, our results are consistent with our previous findings suggesting that crossovers in guppy male meiosis are often restricted to physically small regions of many chromosomes, and that LG12 is not exceptional. Therefore, if the sexual dimorphism in crossing over evolved to restrict recombination of the Y chromosome with the X, perhaps due to the Y carrying sexually antagonistic factors such as male coloration factors, any such evolutionary change was probably not specific to the XY pair.

The localization of crossovers in male meiosis in all guppy populations tested suggests that this pattern probably evolved before the establishment of any male coloration polymorphisms that evolved within individual populations (Bergero, et al. 2019). If so, it might be similar in closely related fish species. Our intron GC content analysis indeed suggests similar GC spikes in *X. maculatus*, the platyfish, and in the even closer relative, *P. picta*, and GC changes do not occur nearer the centromere in the platyfish than in the guppy (Figure 4). The two chromosomes where the platyfish does not show strongly terminal GC spikes are not conclusive evidence for less terminal crossover localization, or for the unlikely possibility that crossovers are strongly localized to physically small internal chromosome regions, because, as mentioned above, assembly errors are possible. In the absence of a male genetic map in the platyfish, we cannot distinguish between these and true differences in arrangements in these species. The GC content differences affect a very large number of sites, and suggest that a very high recombination rate in these terminal chromosome regions has persisted for a long evolutionary time. The high rate probably affects the terminal region uniformly (rather than there being a hotspot near the PAR boundary), as the most terminal scaffold mapped has a uniformly high GC content (see Supplementary Figures S8A and S9). However, we cannot exclude the possibility that there could have been subtle changes in either the platyfish or the guppy lineage that the intron GC values cannot detect (Supplementary Figure S8B).

Although no genetic map has been estimated from crosses within the platyfish, mapping of microsatellite markers in a large progeny of an cross between Xiphophorus species did not suggest a consistent sex difference (Walter, et al. 2004). Many LGs had female maps exceed 50 cM, suggesting that multiple crossovers might occur on at least some chromosomes, unlike the single crossovers in the guppy (see above). The lengths of the maps suggest that terminal regions are often included, including any regions with high recombination rates in males, but the genome had not yet been sequenced and so genetic and physical lengths could not be compared to determine crossover localization patterns. A more recent genetic map also used an inter-species cross, and mapped RAD markers, whose positions in the genome assembly are known (Amores, et al. 2014); however, the map was estimated from an F1 female, and distorted segregation ratios also affected the map. The two genetic maps from females do not yield correlated map lengths for the chromosomes (the r^2^ value for a linear regression is only 0.009). It is possible that the interstitial crossovers could result from displacement to these regions due to rearrangements elsewhere on the chromosomes in these hybrids, as such effects are produced in heterozygotes for rearrangements (Mary, et al. 2018), which can also change the relative proportions of crossover and non-crossover outcomes (Crown, et al. 2018). Overall, given these uncertainties, the *Xiphophorus* maps do not contradict our results suggesting terminal localization of crossovers on platyfish chromosomes, but suggest a need for more mapping using crosses among *X. maculatus* individuals; it remains possible that the crossover pattern has changed in the lineage leading to either *Xiphophorus* (adding interstitial crossover events), or to the guppy (with loss of even occasional interstitial events). We therefore conclude that extreme crossover localization probably did not evolve recently in the guppy lineage, but may represent an old-established situation with very extensive pericentromeric regions showing low crossover rates in male meiosis.

### When did the sex difference in recombination evolve?

Heterochiasmy, however, is probably weaker in the more distantly related fish medaka, *Oryzias latipes*, as genetic mapping in this species suggests that only the sex-determining region itself fails to recombine in male meiosis, while most regions of the sex chromosome may are pseudoautosomal (Kondo, et al. 2001). The same applies in several cichlid fish, in which analyses of F_ST_ between the sexes yielded clearly localized regions that show signals of complete sex-linkage, and extensive pseudo-autosomal regions e.g. (Gammerdinger and Kocher 2018), implying that recombination occurs in male meiosis in the latter. The state in the guppy and its closest relatives therefore probably evolved after the split with the cichilds and *Oryzias* species, though It remains uncertain when the change occurred.

Our evidence that crossing over in male meiosis is highly localized to the tips of many guppy chromosomes, and similarly localized in the platyfish (very likely also in males only), supports our previous proposal that the guppy Y could have arisen by an event that brought a male-determining factor onto a chromosome in an ancestral species with such sexually dimorphic crossing over, and that this event simultaneously produced the male-specific crossover pattern, instantly isolating this chromosome from its homolog and creating an “XY” pair (Bergero, et al. 2019), without any evolutionary change in the sexual dimorphism in crossing over in the guppy.

### Evolution of recombination patterns versus sex-limited expression

Most guppy male coloration traits are Y-linked, but those that are partially sex-linked show male-limited expression (Lindholm and Breden 2002). The concentration of male coloration genetic factors on the guppy LG12 is consistent with theoretical models showing that complete or close partial linkage to the sex determining locus is favorable for the spread of sexually antagonistic factors in populations, other things being equal (Jordan and Charlesworth 2012). If, as just suggested, recombination was already rare in males, the coloration factors that became established in populations may have had time to evolve male-limited expression. Testosterone treatment experiments reveal that almost all females in natural guppy populations from low-predation localities in four Trinidad rivers carry coloration factors that they do not normally express (Haskins, et al. 1961; Gordon, et al. 2012). In females from high-predation sites, the proportions were consistently lower, but still high (between < 10% up to 80%); these polymorphic factors may be rarer in males from these populations, so the difference need not suggest any difference in the control of expression of the traits studied. As the coloration factors have not yet been identified, their frequencies are currently difficult to estimate.

Sexually antagonistic selection may be involved in the evolution of sex chromosomes through selection for reduced recombination, but resolving conflicts by evolution of sex differences in gene expression may also be important when alleles that are beneficial in one sex spread, and the other sex experiences deleterious effects. It has been suggested that this might have prevented the evolution of suppressed recombination with the sex-determining locus, accounting for the maintenance of large partially sex-linked regions in Paleognathous birds (Vicoso, et al. 2013), whereas in Neognathous birds ZW recombination has become suppressed, perhaps due to selection generated by partially sex-linked SA polymorphisms. This idea appeared to be supported by estimates suggesting that many emu PAR genes show sex biased expression, but this is probably not the case (Xu, et al. 2019). Nevertheless, expression evolution of partially sex-linked SA genes could indeed lessen or eliminate the selection pressure for closer linkage. Given our evidence that the guppy crossover pattern is similar to that in the platyfish, it may be interesting in the future to study expression of genes on the guppy sex chromosome.

## Methods

### Fish samples, DNA extraction and genetic mapping

Genetic mapping data were obtained by high-throughput genotyping (SeqSNP) experiments, using SNPs ascertained from our own resequencing study of Trinidadian guppies (10 males and 6 females) sampled from a natural population (Bergero, et al. 2019). We selected an excess of SNPs at both ends of each chromosomal assembly for genotyping, in order to maximize the chance of detecting crossover events in male meiosis, assuming that these events might be localized in the chromosomal termini. The guppy families used for the high-throughput SNP genotyping are described in Charlesworth et al. (2020).

Genomic DNA for genotyping was extracted using the Echolution Tissue DNA Kit (BioEcho, Germany). For SNP ascertainment for SeqSNP genotyping, we identified genic sequences found in all 16 *P. reticulata* individuals sampled from a captive population recently collected from a natural site with a high predation rate in the Aripo river, Trinidad (Bergero, et al. 2019). The SNPs to be targetted were selected from within coding sequences, with the criterion that about 50 bp of sequence flanking each such SNP should also be coding sequence, in order to maximise the chance that the sequence would amplify in diverse populations, and to minimise the representation of SNPs in repetitive sequences. To further avoid repetitive sequences, the SNPs were chosen to avoid ones whose frequencies in the ascertainment sample were 0.5 in both sexes. The SNPs and their locations in the guppy genome assembly are listed in the file “Autosomal segregation (6 fams HT results)” **in Dryad (doi to be added**). The experiments were carried out by LGC Genomics (LGC Genomics GmbH, Ostendstraße 25, 12459 Berlin, Germany, www.lgcgroup.com/genomics). As expected, the primers work well for most targeted sequences.

The SNP genotype data were analysed for the autosomes as well as the guppy XY pair, chromosome 12. For each chromosome, we examined the progeny genotypes and recorded the locations of crossover events, in order to show which regions co-segregated in male meiosis, and which recombined. When the data for a chromosome included SNPs informative in male meiosis near both ends of the assembly (Supplementary Figure S6), we recorded which end of the assembly showed the crossovers, for comparison with the positions of elevated GC content (estimated as described below). Maps based on the lAH family, with the most informative markers, were estimated using LepMap3 (Rastas 2017) and are shown in Supplementary Figure S7.

### Analyses of genome sequences

#### Preparation of datasets

The female guppy genome assembly and annotation files were obtained from the NCBI Annotation Release 101 (GenBank under accession number GCF_000633615.1). There are 22 pairs of autosomes and one sex chromosome pair. The southern platyfish (*Xiphophorus maculatus*) genome assembly, with 24 chromosome pairs, was downloaded from NCBI Annotation Release 102 (under accession number GCF_002775205.1 at the GenBank). The GFF3 files for both species were downloaded from Ensembl (release 97), to provide gene names and sequences, and the exon and intron locations for each transcript in the longest transcript for each gene, as many guppy genes are annotated with multiple transcripts. Scripts for the analyses are deposited in Dryad, under accession number **XXXX [to be added]**.

#### Intron GC content analysis in *P. reticulata and X. maculatus*, including determining chromosome homologies between the two species

For each gene, the coordinates of the exon ends were converted to yield the intron positions, and the intron sequences of every gene from each chromosome were retrieved from the genomic assembly, and their GC content values were calculated using the code in the file ‘maincode.py’ deposited in Dryad. Genes with no introns were not included. The set of genes from LG12, the XY pair, includes 824 genes with at least one intron (56 of which had just a single intron), and 44 with no introns (the numbers for other chromosomes are given in Supplementary Table S1). For genes with more than a single intron, the GC values for all introns were pooled, as the values for first introns did not differ significantly from those of other introns in either the guppy of platyfish (see Results section).

We note that the assemblies of guppy and platyfish chromosome homologues show multiple breaks in synteny, which may be true rearrangements between these species, or could, in some or all cases, represent assembly errors (Supplementary Figure S1 and files in Dryad folder “PNG+TRACE_G (dot plots)”). As discussed in the Results section, assembly errors that incorrectly order genes on a chromosome can lead to elevated GC values in the incorrect chromosome region; in addition, genes assigned to the wrong chromosome will produce a false appearance of crossing-over within the chromosome. We assessed the latter possibility by comparing the gene contents of the guppy and platyfish chromosomes, following the approach described by (Schartl, et al. 2013). This revealed few indications of such problems (files in Dryad folder “GUPPY-platy chromosome homologies”). Supplementary Figure S1 shows the example with the largest such region we detected: most of LG6 corresponds to the platyfish chromosome 2, but a region of several Mb corresponds to part of the platyfish chromosome 3 (most of which corresponds to the guppy LG16).

##### P. picta

Sequences were made available from the lab or Cameron Ghalambor. Reads were first stringently quality controlled as follows. The reads were processed for adapters with Trimmomatic ILLUMINACLIP. ‘Standard’ settings of 2:30:10 were used, allowing at most two mismatches in the adapter seed, and quality cutoffs of 30 and 10 for paired or unpaired reads, respectively; only paired reads were retained. Trailing Ns at the starts and ends of reads were removed, as were trailing sections with quality < 3. A sliding window of size 4 was applied to the read, and clipped if mean quality, as assessed by fastqc analysis, dropped below 15. Reads of shorter than 108 bp were discarded. Next, the reads were aligned to the guppy reference genome sequence, using BWA mem (Li and Durbin 2010) with the default settings. The alignment was converted to BAM format with Samtools, then sorted and indexed. As a further quality control, only reads that mapped as a pair correctly were retained; optical or PCR duplicates, and reads with mapping quality ≤ 30, were discarded. Introns were called from the guppy GFF3 annotation from Ensembl using the ‘-addintrons’ option of GenomeTools (http://genometools.org/tools/gt_gff3.html), and the output was processed to produce a bed file with the intron regions of each guppy chromosome based on the GFF3 chromosomal annotation.

The intronic bed file was sorted, and bedtools map was used to concatenate all reads from a single individual *P. picta* female that mapped to a given intron in the chromosomal alignment, and the proportion of GC bases was computed for each intron.

The individual chosen for analysis was AWCSU02 / N705 / AK 403; CAR_H female; identifier number 7 (for comparison with other experiments). This female was selected due to having coverage roughly 3-fold higher than other individuals in the same population. The increased coverage was not due to over-representation of repeats or regions of poor quality sequence: the library reduction during quality control was similar to that for most other individuals; for example, the RPK of LG12 in this female dropped from ~56 to ~55 after Trimmomatic processing, similar to the decrease from ~18 to ~17 for the female with identifier number 6 who was typical of the population. The observed differences in GC content between chromosome regions might nevertheless be inaccurate, but there is no reason to suspect any sytematic difference that could cause the differences we observe. First, regions with extreme base composition, in either direction, generally have low coverage in PCR-based sequences such as those analyzed (Benjamini and Speed 2012), and the resulting lower representation of such regions will increase the variance in GC content. Second, our fastqc analysis allowed the inclusion of many AC and GT repeat sequence that just passed the 0.1 threshold. Finally, repetitive regions tend to be AT-rich, and if they were preferentially excluded from our analysis, the overall GC content might be over-estimated; however, this seems unlikely to affect differences in GC between introns of genes in different chromosome regions.

Our analysis retained the longest isoform of the coding sequence for each gene, and analysed only the introns based on this isoform. To do this, the table of GC proportions per intron for this female was filtered to exclude introns with more than one read, and redundancy removal was enacted to remove those with the same start site, and keep those with the longest isoform (latest end point), and those with the same end site, again keeping the longest isoform (earliest start site).

#### Change-point analysis

We visualized the GC values along the guppy chromosomes using the LOESS package (Jacoby 2000) in R (Ihaka and Gentleman 1996) to plot smoothed lines and 95% confidence intervals. To examine the significance of the changes, we used the cumSeg package (Muggeo and Adelfio 2010) in R to detect change-points in the GC content in the chromosome assemblies. This change-point analysis software is designed for analysis of genome sequences, and it estimates the number and location of significant change points using a non-parametric test, and plots the mean value of the quantity of interest (here, GC content) in each segment identified. For most chromosomes, we applied model 3, which yields the best estimates. For a few chromosomes in the guppy or platyfish where no clear change-point was identified, model 2 were also used. When two consecutive change points were detected, we tested for differences between the species either by choosing the most distal of the change-points to compare, or the more proximal one of the two. Supplementary Table S4 shows the five chromosomes where the results differ. Because different change-point analysis approached yield differing results, we also applied a different method, Pelt, implemented in the “ruptures” package (arXiv:1801.00826); we used the “Pelt” method, because this estimates the number of breakpoints, rather than the user specifying a fixed number, with the default parameter values (Truong, et al. 2020).

#### GC3 values in exon sequences

To examine whether coding sequences also show evidence for GC-biased gene conversion in regions with high recombination rates, we also computed the GC3 (G+C in the third positions of codons) for the exon sequences of the genes whose introns were analysed. The coding sequences were extracted using the GFF annotation together with the genome sequence of each chromosome using a Python script from R. Ness (University of Toronto). The Python module GC123 from the Biopython SeqUtils package (Cock, et al. 2009) was used to calculate the GC3 value for each gene.

#### Gene density analysis

The gene densities for each guppy chromosome were estimated from the ‘genomicDensity’ package in R using sliding windows of size of 500,000 bp and a 250,000 bp gap, and the intronic GC content of genes in each window.

#### Estimating between-species divergence for orthologous genes

To estimate divergence between guppy and platyfish sequences, and to understand the relationships between the species studied, we compared genes on homologous chromosomes, determined as described above. We used BLAT v38 to find orthologues by reciprocal best hits between sequences of guppy cDNAs with chromosomes assigned in the female assembly, versus cDNAs from the complete genome sequences of the platyfish or *Gambusia affinis*. After amino acid sequence alignment using MAFFT (Katoh and Standley 2013), the resulting DNA sequences were analyzed using PAML software, to estimate synonymous the site divergence (K_s_) values summarised in Supplementary Figure S2.

Divergence analysis was also done to test whether the mutation rate is higher in regions with high GC content than other chromosome regions, based on substitutions between the guppy and platyfish. For this, orthologous gene pairs were found using reciprocal BLAST of coding sequences from the two species available from the NCBI repository; to avoid paralogous genes, as far as possible, and to relate the divergence values to the genes’ chromosomal locations, the BLAST tests were conditioned on each being present on the homologous chromosomes of the other species, as determined using the NUCmer function of MUMmer 3.0 (Kurtz, et al. 2004). Reciprocal best hit pairs were selected based on the e-value, score and identity by parsing the output of both files from the BLAST. Finally, as the guppy and platyfish are diploid, many sites in the coding sequences have codes indicating heterozygosity, we assigned one of the two bases randomly at each heterozygous site. Since the synonymous site divergence between the two species is generally < 10% (Supplementary Figure S2A), our estimates of raw sequence divergence will be only slightly affected by neglecting intra-species differences. The resulting single coding sequences for each gene were then aligned in MAFFT version 7, using the default settings (Katoh and Standley 2013). Each alignment was checked manually, and a few obviously unreliable segments were discarded, including incomplete coding sequence in either species, pairs with stop codons detected in either sequence, and sequences whose total the length was not a multiple of three (overall 5.1% of sequences were excluded from the analysis, and the percentages from each chromosome were similar, see numbers in Supplementary Table S2).

The alignments after filtering were used to estimate divergence for each gene, using the KaKs_Calculator 2.0 software (Wang, et al. 2010) to estimate K_s_ values for genes with known positions in the guppy chromosomes. We applied the three following models: NG (Nei and Gojobori, 1986), LPB (Li, 1993; Pamilo and Bianchi, 1993), and the GY (Goldman and Yang 1994) codon-based model, which corrects for saturation. Although saturation is unlikely over the evolutionary times separating these species (see Results section and Supplementary Figure S2A), correction may be necessary for detecting local genome regions with unusually high divergence, which could characterize regions with very high recombination rates (see Introduction section).

## Acknowledgements

The project was supported by ERC grant number 695225 (GUPPYSEX). We thank A.J. Wilson and D.P. Croft (University of Exeter) for help with collecting samples of wild *P. reticulata* from Trinidad, and animal technicians at U. Exeter (Cornwall campus) for fish husbandry and maintenance. We also thank Dr. R.W. Ness (University of Toronto) for assistance with analysis of annotated genome sequences, and Dr. Cameron Ghalambor (Colorado State University) for sharing *P. picta* genome sequences.

## Supplementary Figures

**Supplementary Figure S1**. Homologies between sequences on the guppy LG9 and on platyfish chromosomes, showing the region discussed in the text where most genes in the LG9 assembly appear in the platyfish chromosome 12 assembly, but a more or less contiguous set appear on 11.

**Supplementary Figure S2**. Intron GC content values for all guppy chromosomes, with changepoints indicated. The y axis shows the GC content of each intron-containing gene, and the x axis shows the assembly positions of the genes whose introns were used.

**Supplementary Figure S3**. Relationships between the species used in GC intron analyses. A. Synonymous site divergence between the guppy versus the platyfish, *Xiphophorus maculatus*, and *P. picta*. B. Schematic diagram of the relationships based on relative synonymous site divergence values, to show that *P. picta* is a closer outgroup than the platyfish.

**Supplementary Figure S4.** GC content of introns in *P. picta* genes with homologs on the guppy LG12. Because the *P. picta* assembly is not contiguous, each dot represents an individual intron. The pattern is therefore less clear than for the guppy, where we pooled introns for each gene (see Figure S2). The red line shows smoothed GC content values (based on a smooth spline with 1/250 the maximum degrees of freedom for each chromosome).

**Supplementary Figure S5**. Ks values estimated by the Goldman-Yang method (see Methods section of main text). Values are shown for all guppy chromosomes.

**Supplementary Figure S6.** Regions of each guppy chromosome with high-throughput mapping data informative in male meiosis (left) and meiosis of the dam (right). In the diagram for the sire meiosis, the red dots indicate the five guppy chromosomes whose left-hand assembly end has the GC spike, when a spike was detected, and blue dots indicate chromosomes for which no spike was detected; all other chromosomes had a spike at the right-hand end of the assembly, except that a middle region spike was also detected for the fusion chromosome, LG2.

**Supplementary Figure S7.** High-throughput genetic maps of all guppy chromosomes with adequate numbers of SNP markers (see Figure S6).

**Supplementary Figure S8.** Possible patterns of crossovers, and the expected GC patterns. A. Crossovers could occur uniformly across the region where they occur, or be localised. B. If crossovers occur uniformly across a large recombining region, the GC content will be only slightly higher in the region. If, however, they occur exclusively in a physically small region, the recombination rate will be very high, and a large increase in GC is expected.

**Supplementary Figure S9.** High GC content in a scaffold that is unplaced in the guppy assembly, NW_007615031.1, but that we found to be located at the terminus of the guppy sex chromosome pair (unpublished result in Charlesworth et al. 2020).

## Supplementary Tables

**Supplementary Table S1**. Gene numbers analyzed for guppy and platyfish chromosomes, and comparisons between first introns and other introns, for genes with more than a single intron.

**Supplementary Table S2**. Information about centromere positions, where they could be determined, and numbers of genes analyzed for divergence between these two species (columns O to Q). The guppy chromosomes are shown at the left, and the chromosome homologies with the platyfish, together with the guppy chromosome sizes. The centromere positions are inferred to be at the chromosome ends opposite the ends with the GC spikes; the spike positions in the guppy, *P. picta* and the platyfish chromosomes are shown in columns I, J and K. Some platyfish chromosome assemblies start at the opposite end from the guppy assemblies, as indicated in column L. Column M indicates whether the analysis identifies the same end of the chromosome as the centromere in the two species, or not, taking such “reversals” into account.

**Supplementary Table S3**. Tests for correlations between gene densities on each guppy chromosome and GC intron content.

**Supplementary Table S4**. List of all change-points detected in the guppy and the platyfish.

**Supplementary Table S5**. Summary of crossover locations on guppy autosomes, and GC spikes.

## Notes

### Competing Interest Statement

The authors have declared no competing interest.

