## Supplementary material for "Using GC content to compare recombination patterns on the sex chromosomes and autosomes of the guppy, *Poecilia reticulata*, and its close outgroup species": Figure S2(CPA_Guppy_length).pdf

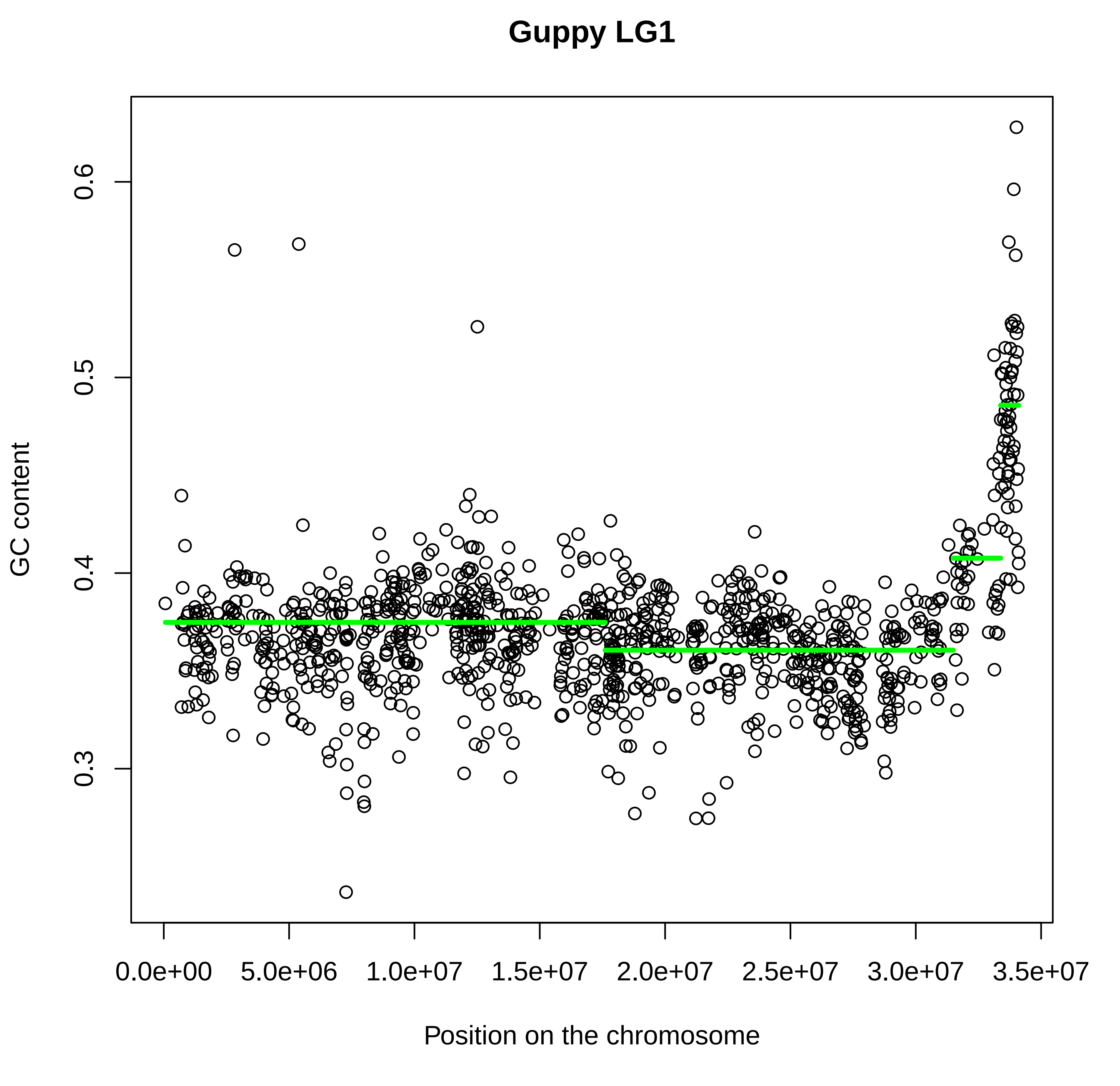
Guppy LG1
GC content
0.3
0.4
0.5
0.6
0.0e+00
5.0e+06
1.0e+07
1.5e+07
2.0e+07
2.5e+07
3.0e+07
3.5e+07
Position on the chromosome

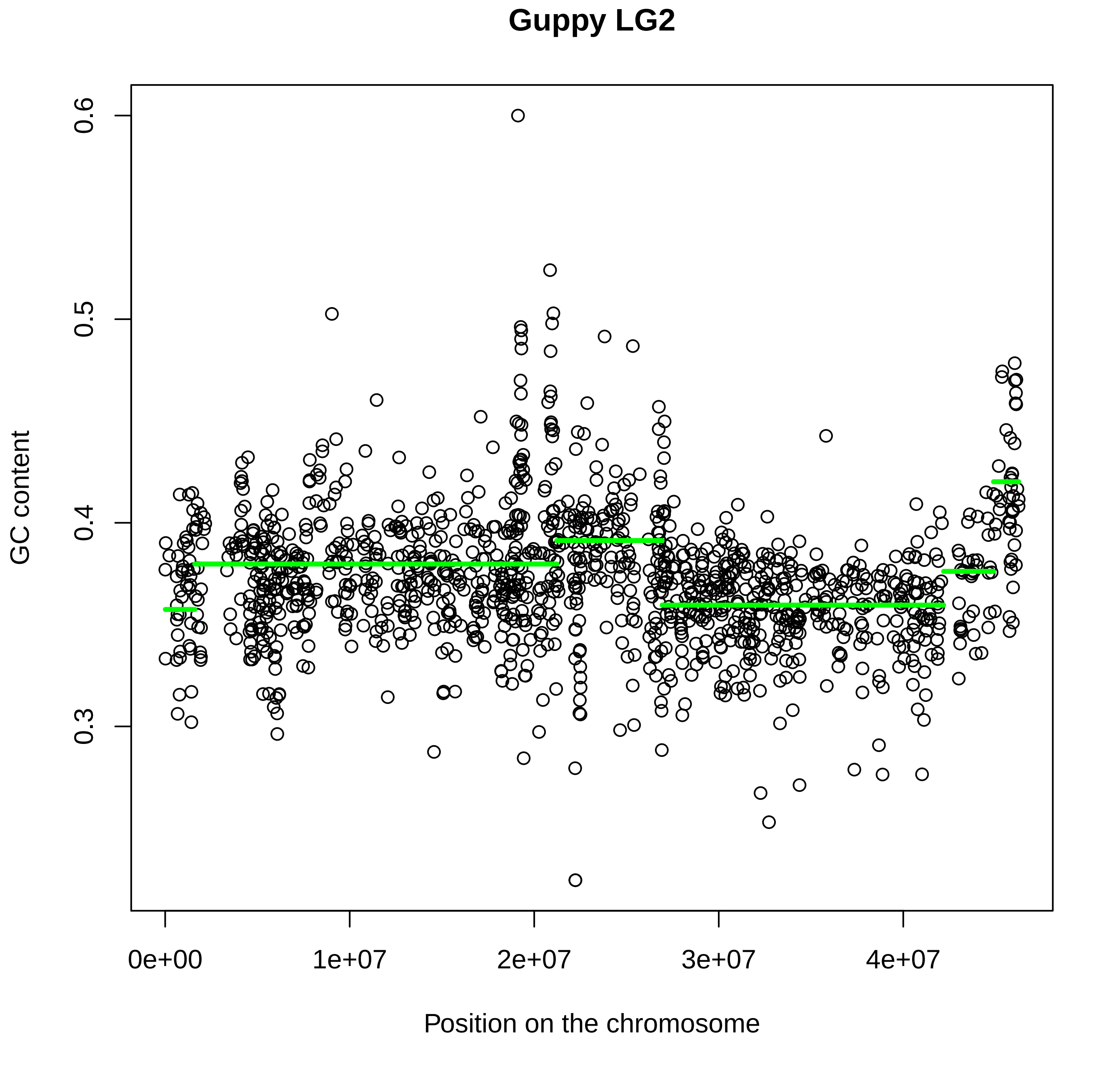
Guppy LG2
GC content
0.6
0.5
0.4
0.3
0e+00
1e+07
2e+07
3e+07
4e+07
Position on the chromosome

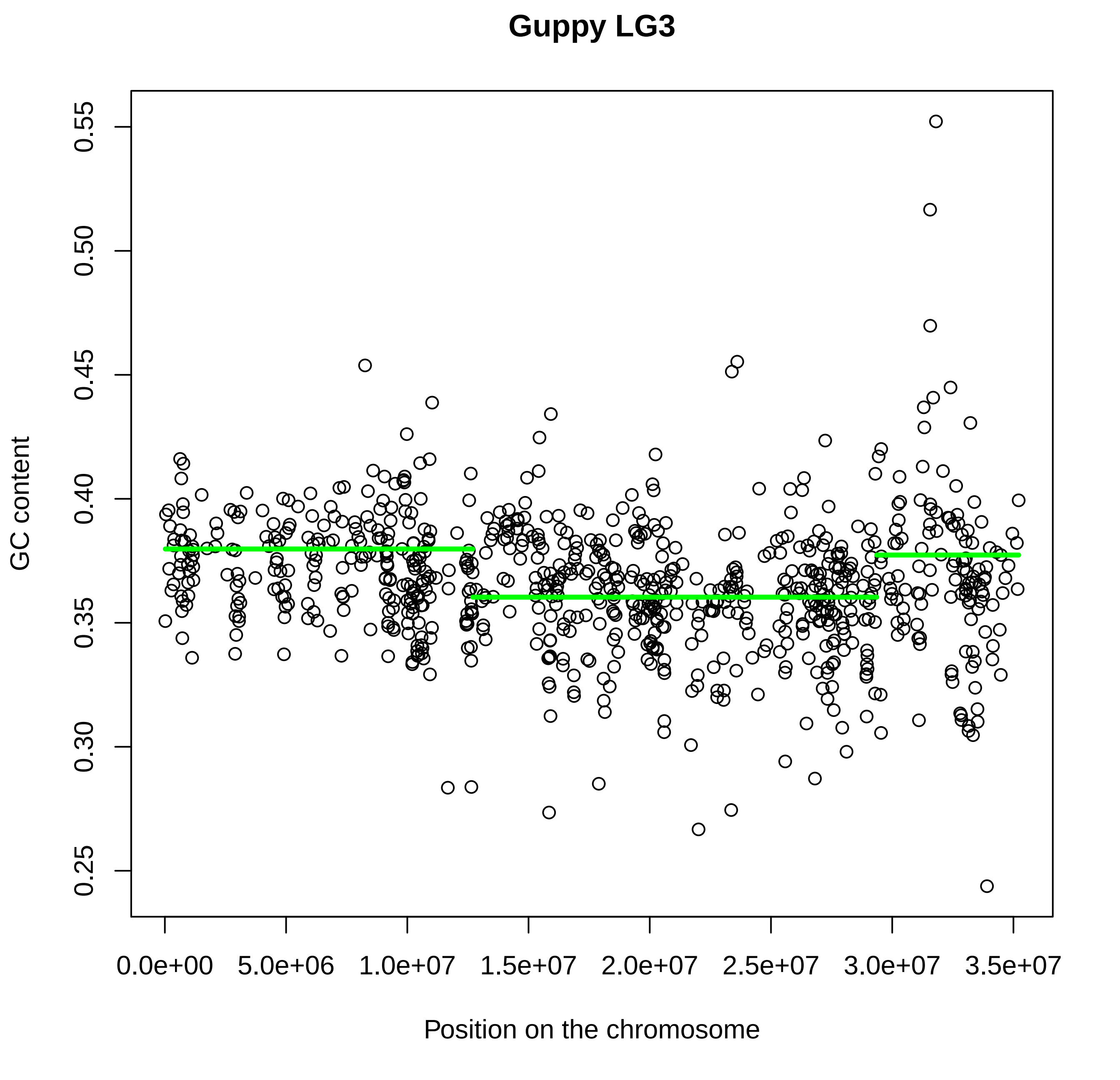
Guppy LG3
GC content
Position on the chromosome
0.25
0.30
0.35
0.40
0.45
0.50
0.55
0.0e+00
5.0e+06
1.0e+07
1.5e+07
2.0e+07
2.5e+07
3.0e+07
3.5e+07

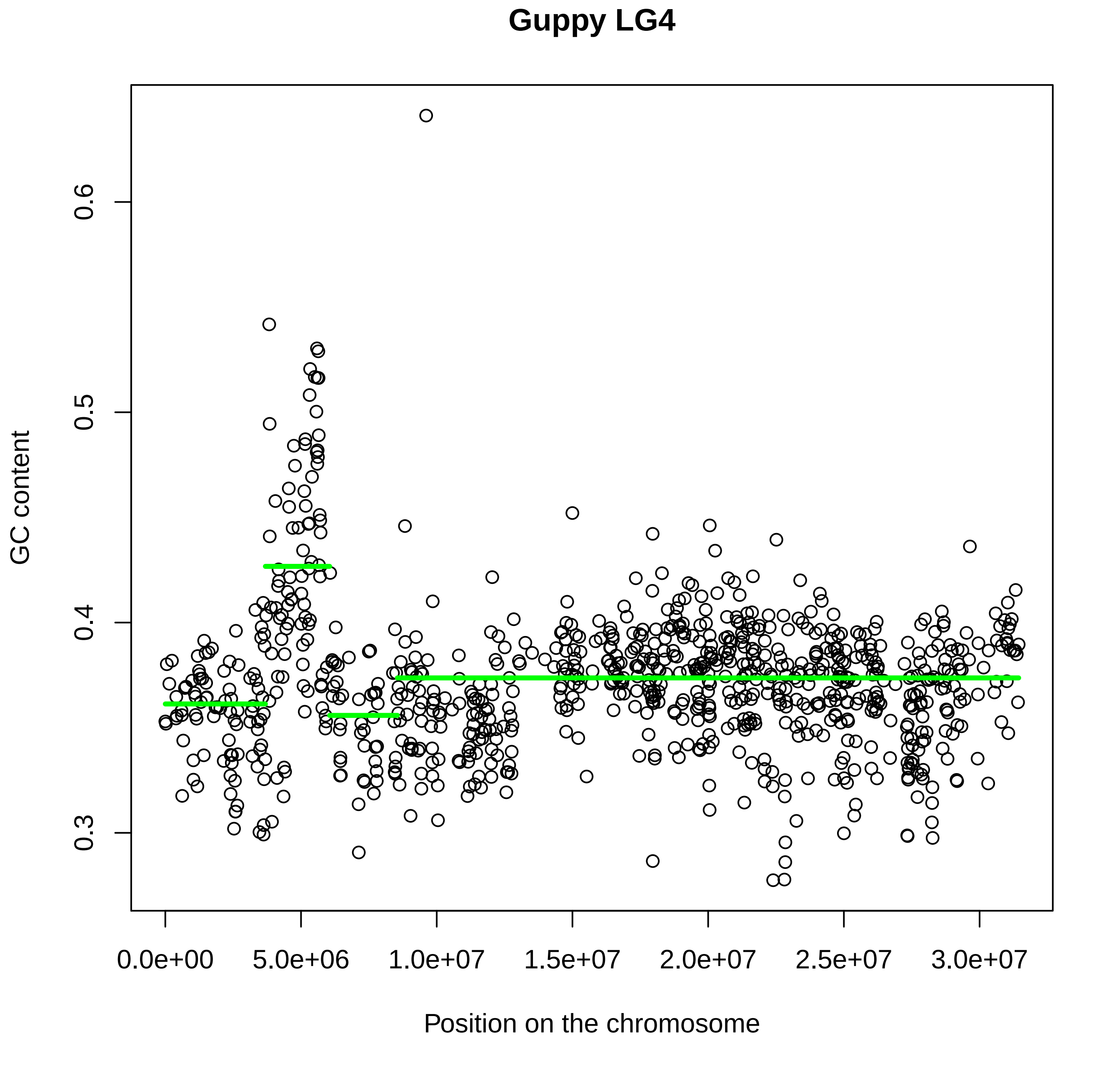

Guppy LG4
GC content
0.3
0.4
0.5
0.6
0.0e+00
5.0e+06
1.0e+07
1.5e+07
2.0e+07
2.5e+07
3.0e+07
Position on the chromosome

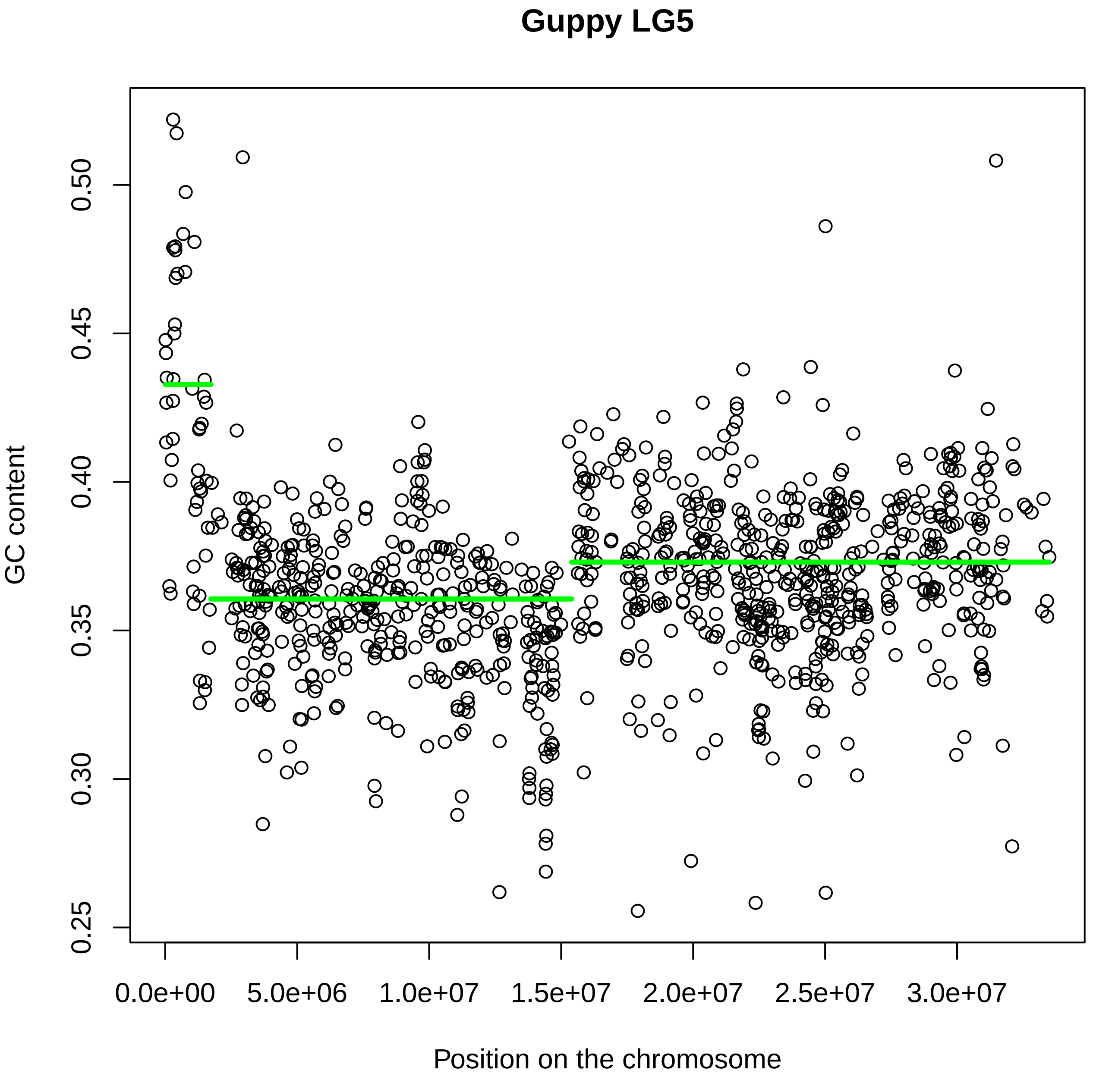
Guppy LG5
GC content
0.25
0.30
0.35
0.40
0.45
0.50
0.0e+00
5.0e+06
1.0e+07
1.5e+07
2.0e+07
2.5e+07
3.0e+07
Position on the chromosome

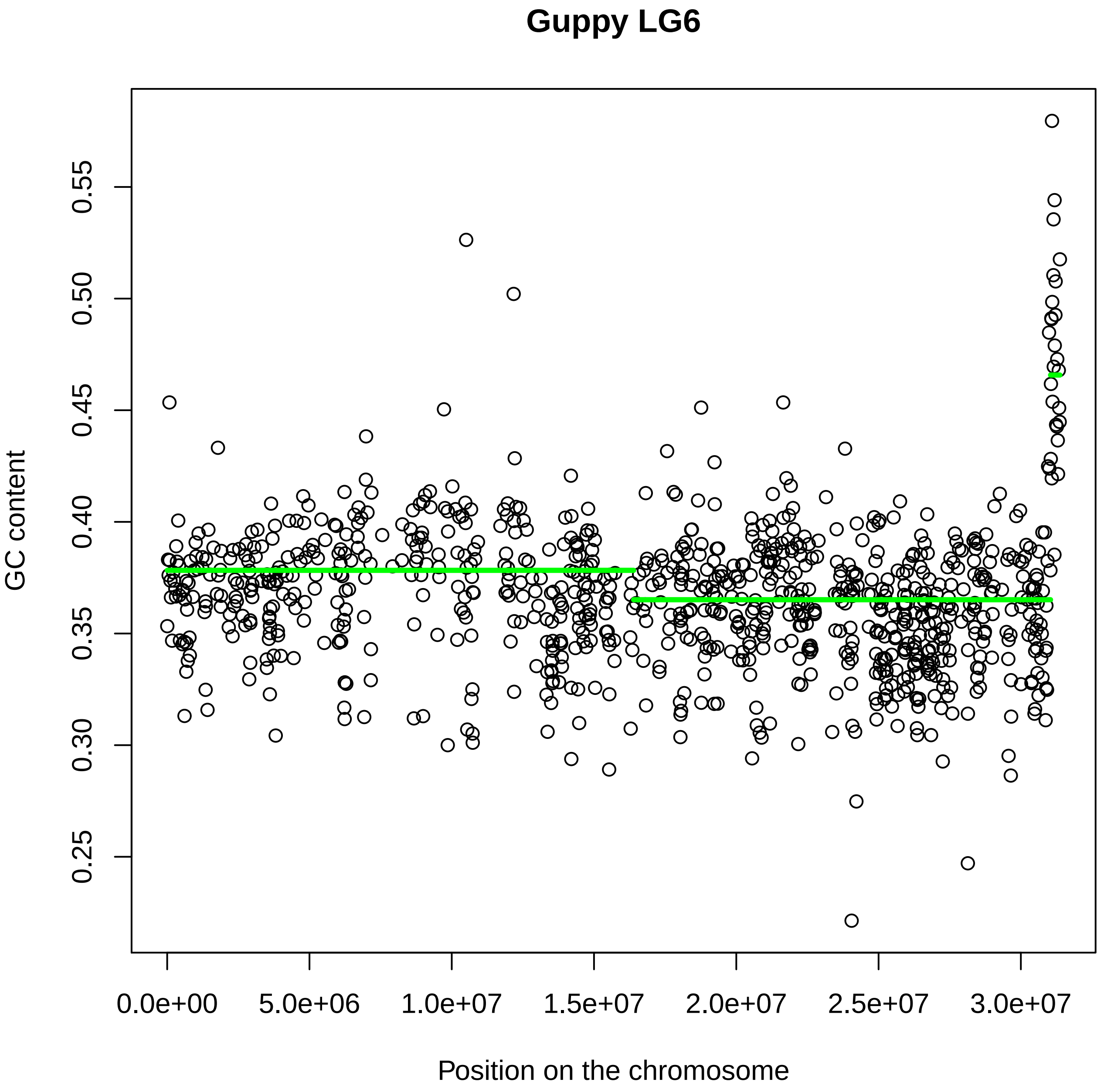

Guppy LG6
GC content
0.25
0.30
0.35
0.40
0.45
0.50
0.55
0.0e+00
5.0e+06
1.0e+07
1.5e+07
2.0e+07
2.5e+07
3.0e+07
Position on the chromosome

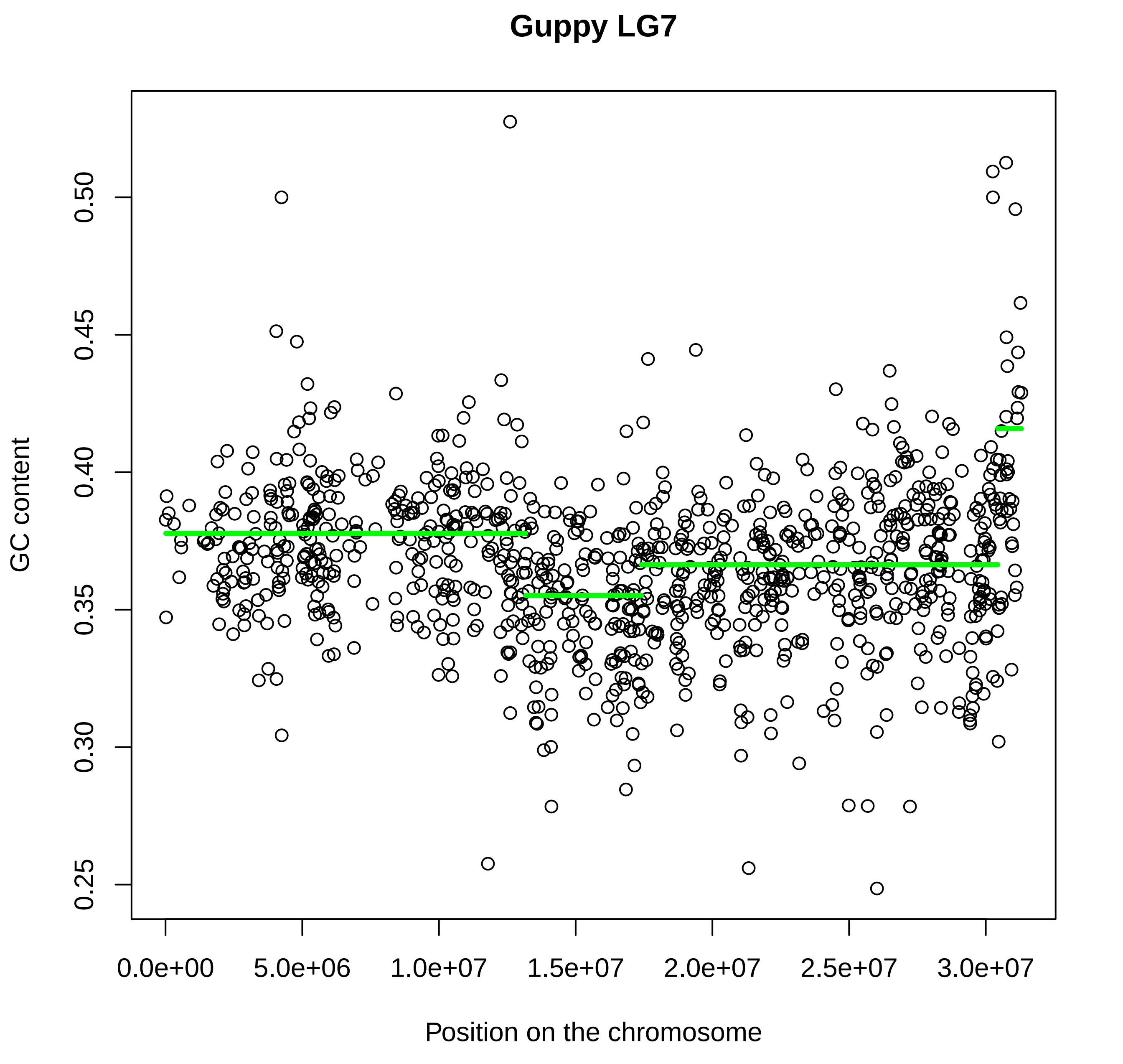
Guppy LG7
GC content
0.25
0.30
0.35
0.40
0.45
0.50
Position on the chromosome
0.0e+00
5.0e+06
1.0e+07
1.5e+07
2.0e+07
2.5e+07
3.0e+07

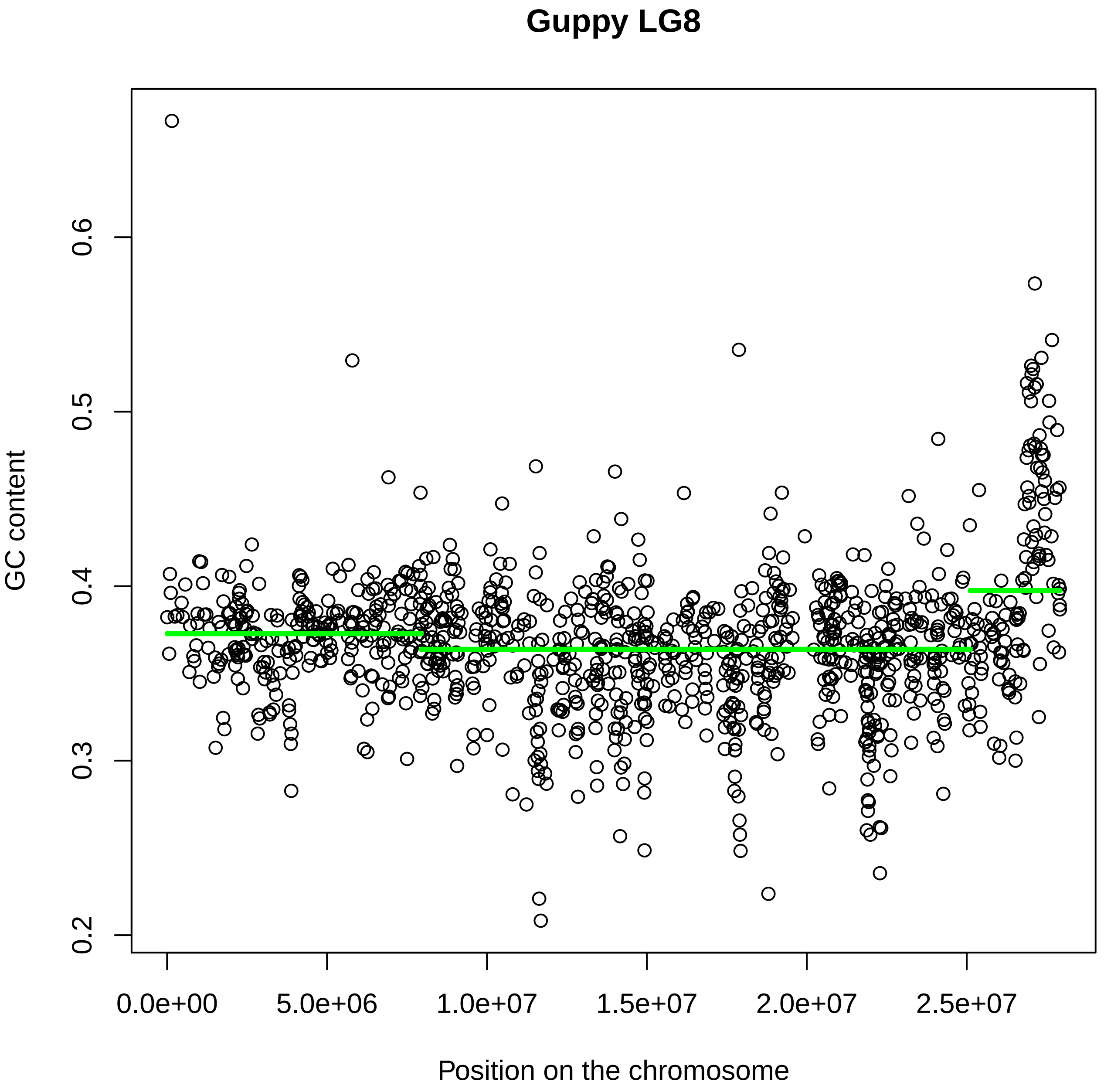

Guppy LG8
GC content
0.2
0.3
0.4
0.5
0.6
0.0e+00
5.0e+06
1.0e+07
1.5e+07
2.0e+07
2.5e+07
Position on the chromosome

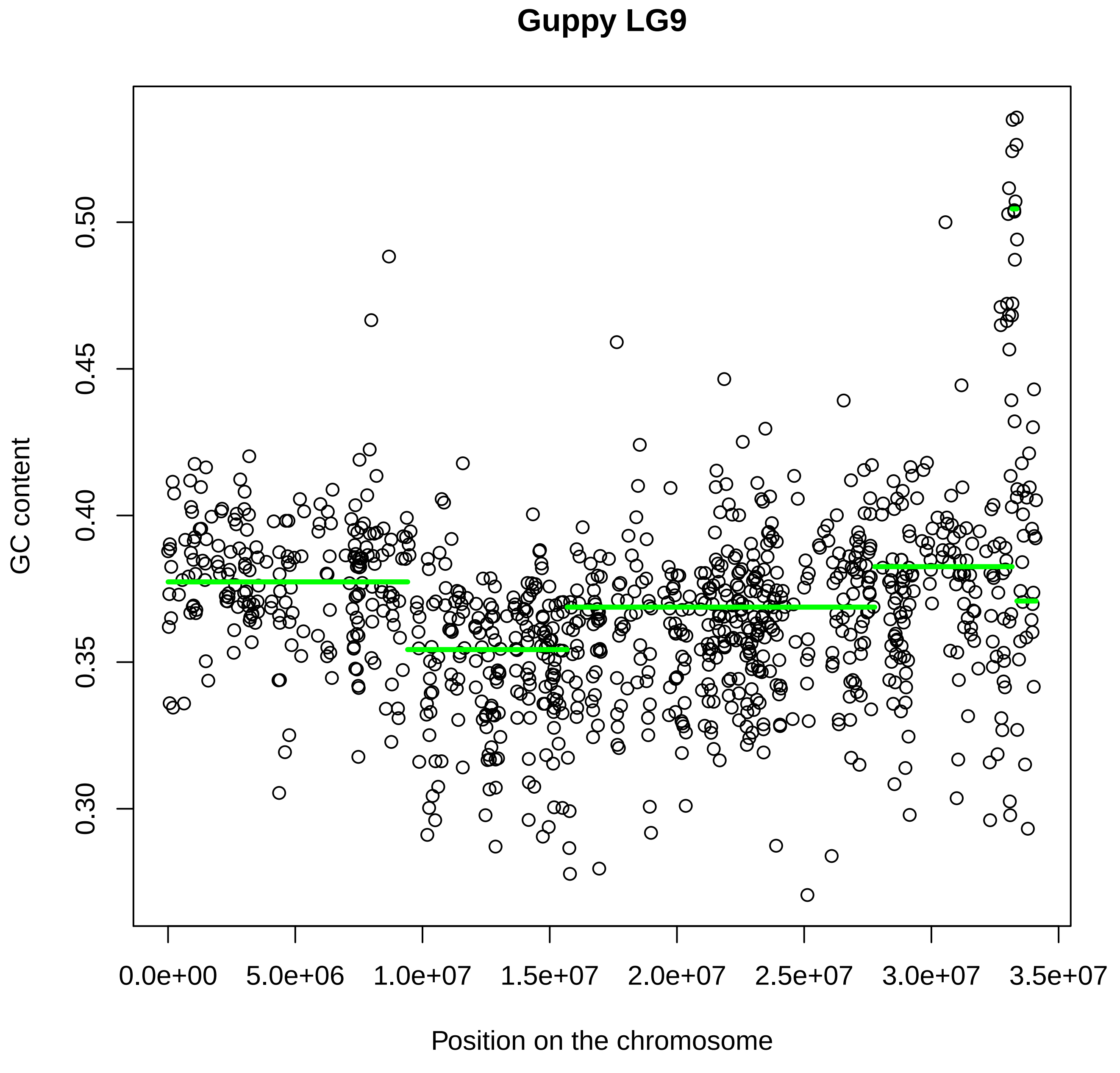
Guppy LG9
GC content
0.30
0.35
0.40
0.45
0.50
0.0e+00
5.0e+06
1.0e+07
1.5e+07
2.0e+07
2.5e+07
3.0e+07
3.5e+07
Position on the chromosome

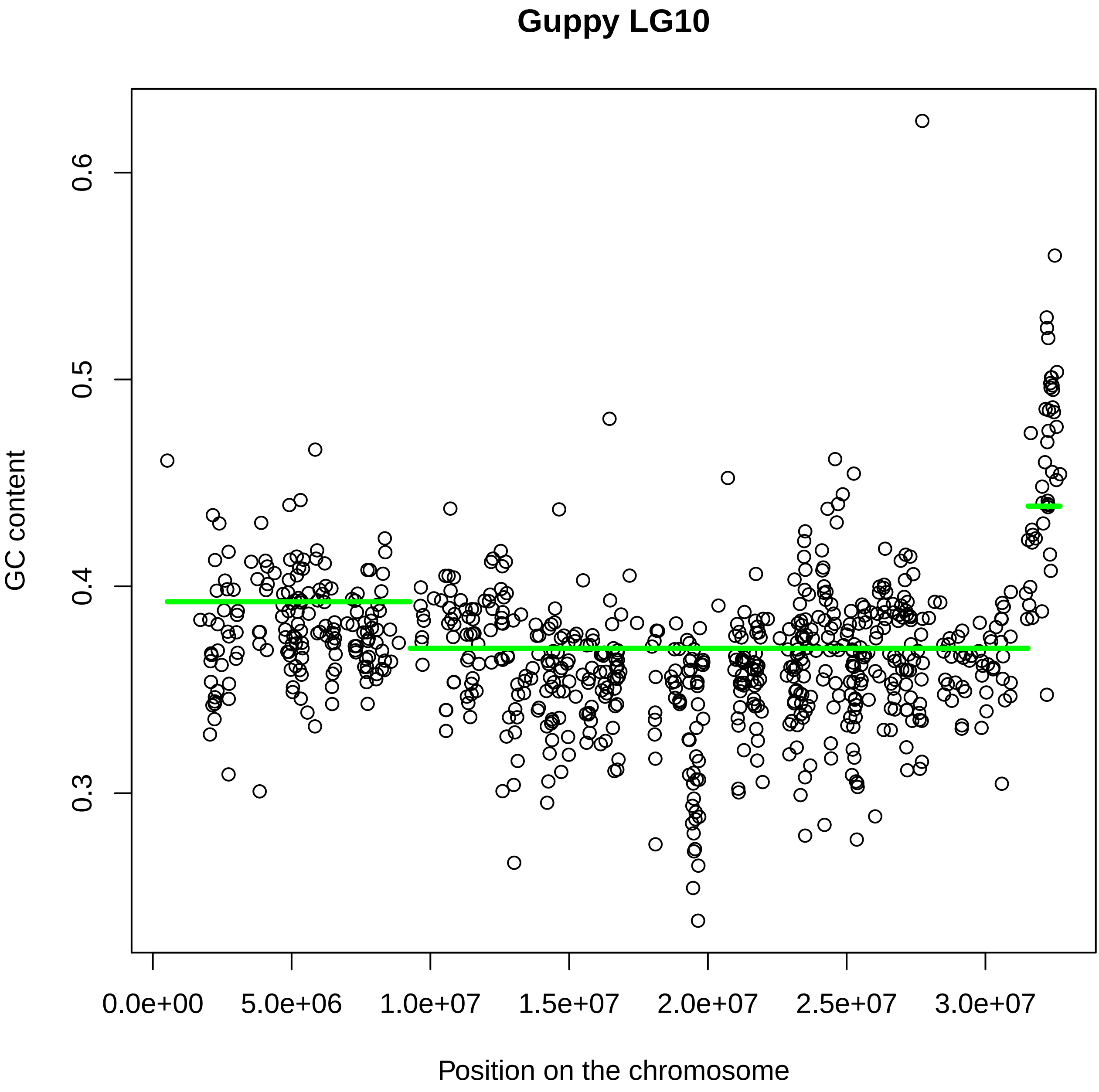

Guppy LG10
GC content
0.3
0.4
0.5
0.6
0.0e+00
5.0e+06
1.0e+07
1.5e+07
2.0e+07
2.5e+07
3.0e+07
Position on the chromosome

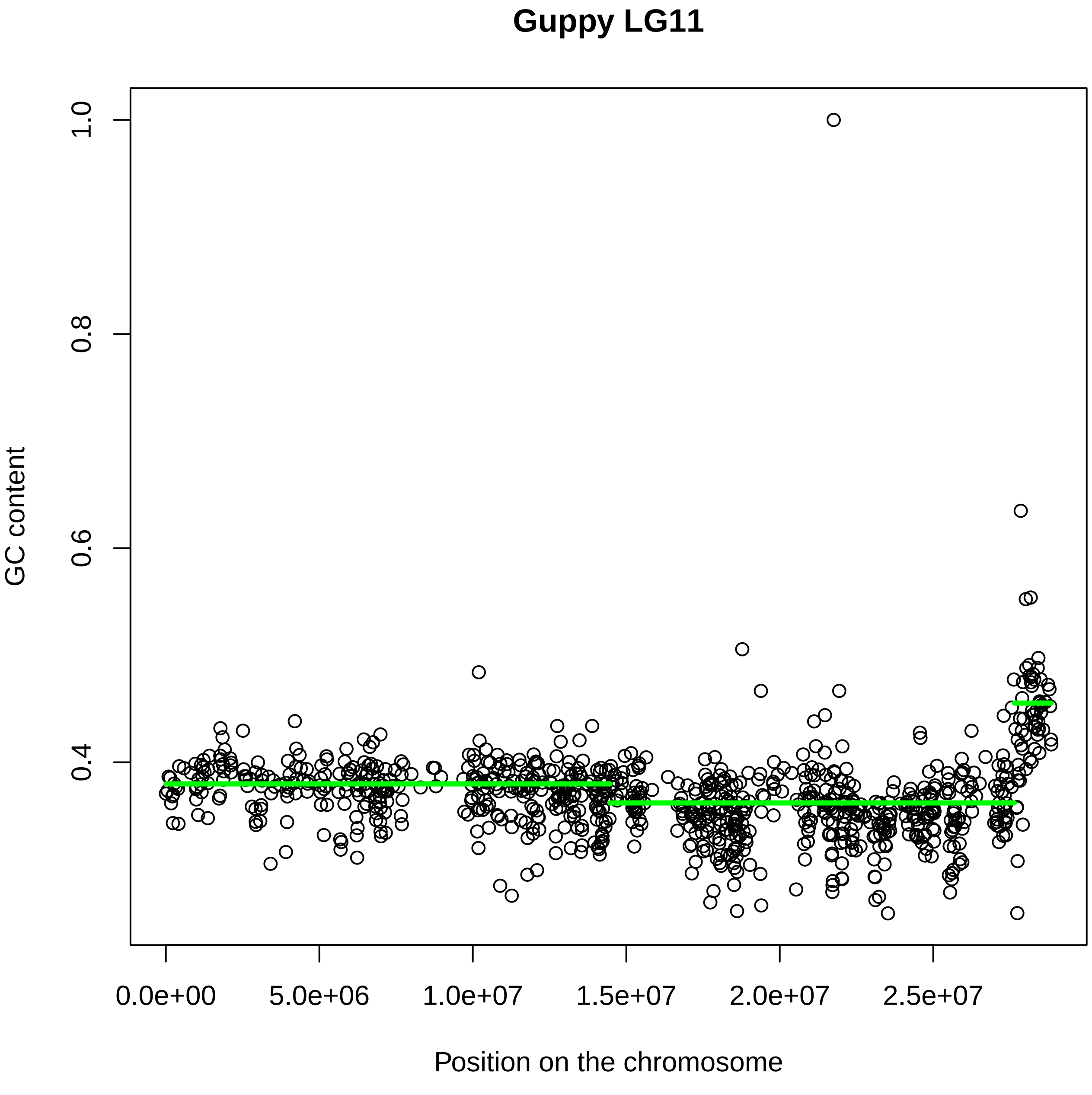
Guppy LG11
GC content
1.0
0.8
0.6
0.4
0.0e+00
5.0e+06
1.0e+07
1.5e+07
2.0e+07
2.5e+07
Position on the chromosome

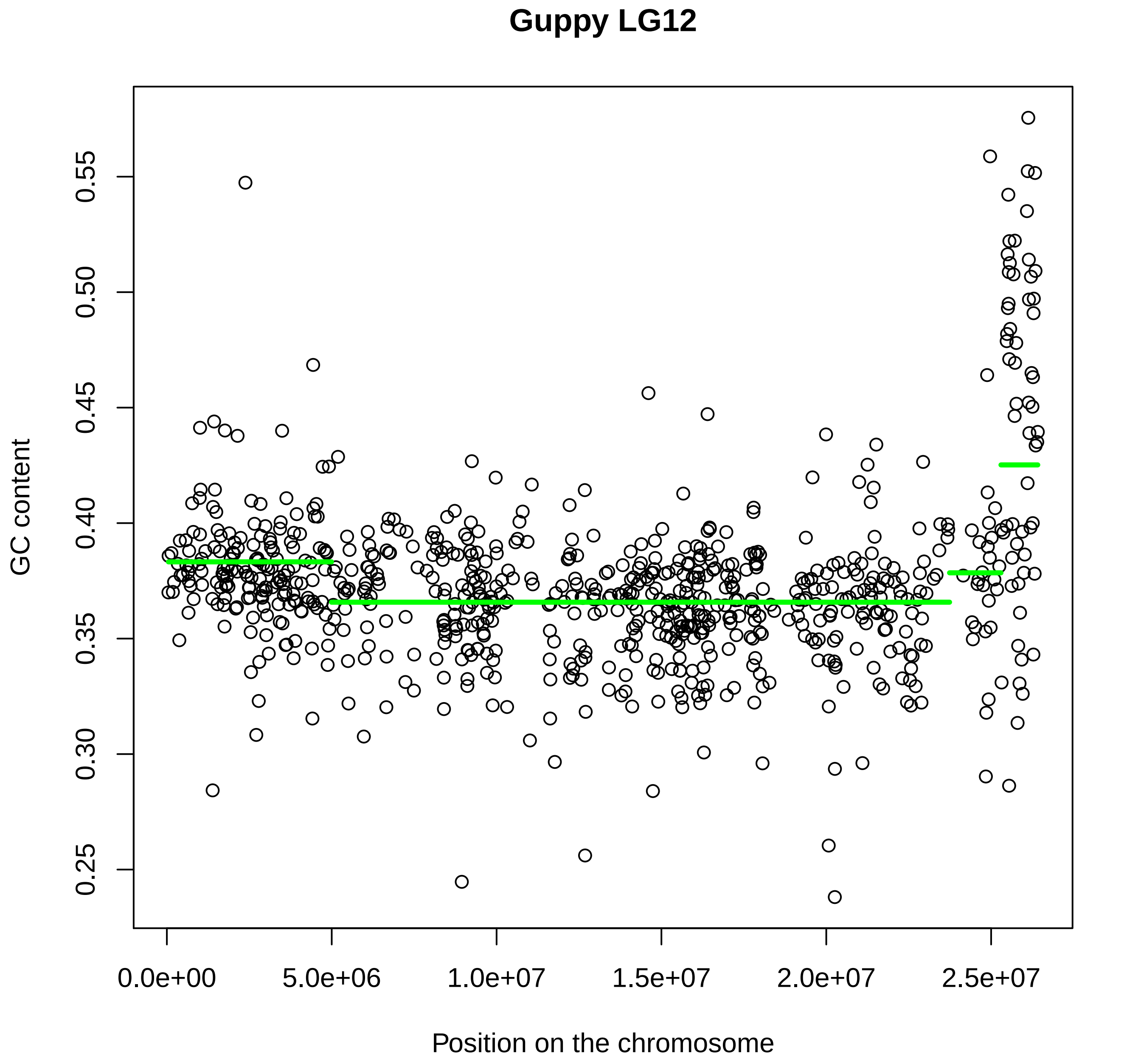

Guppy LG12
GC content
0.25
0.30
0.35
0.40
0.45
0.50
0.55
0.0e+00
5.0e+06
1.0e+07
1.5e+07
2.0e+07
2.5e+07
Position on the chromosome

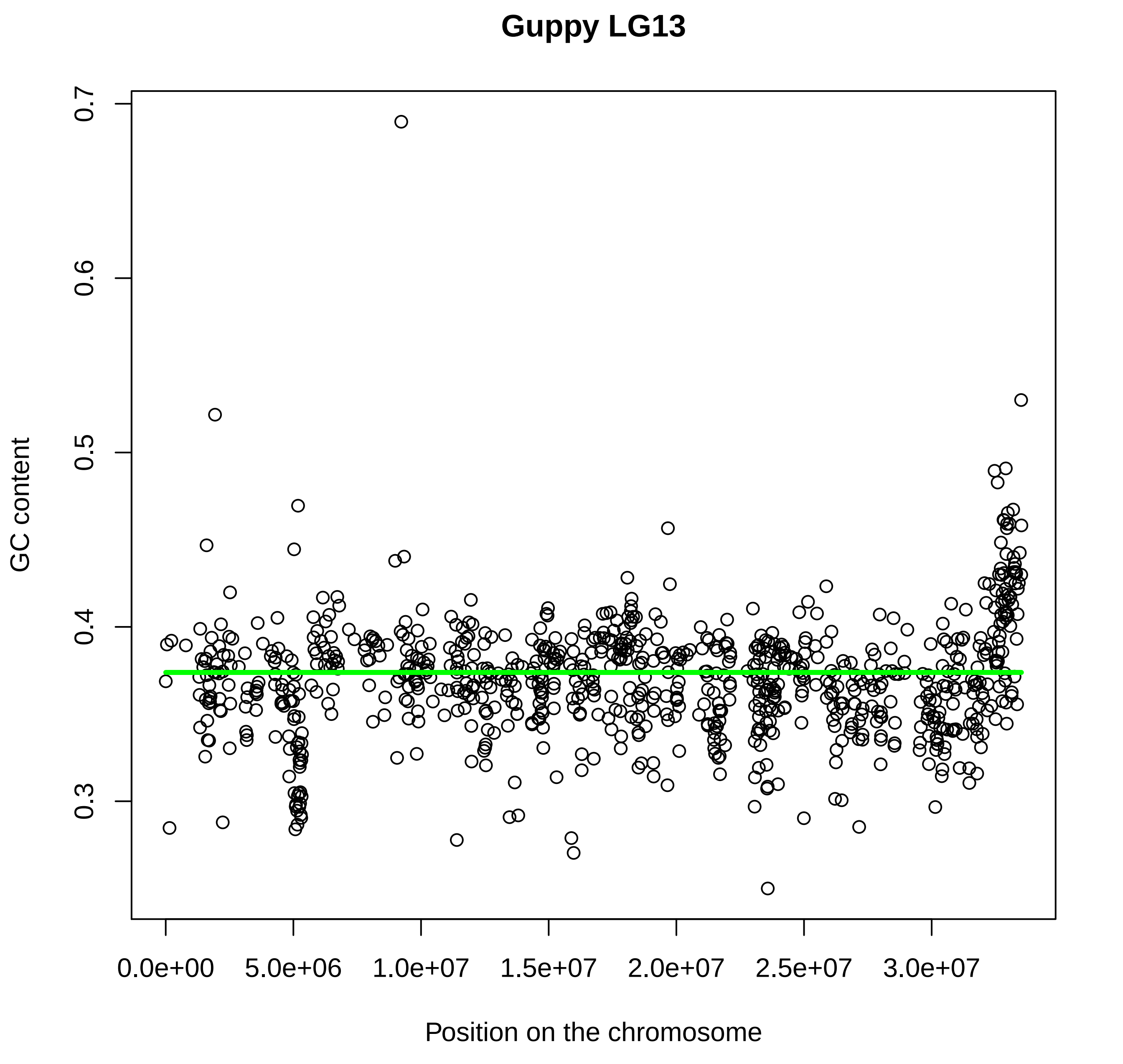
Guppy LG13
GC content
0.3
0.4
0.5
0.6
0.7
0.0e+00
5.0e+06
1.0e+07
1.5e+07
2.0e+07
2.5e+07
3.0e+07
Position on the chromosome

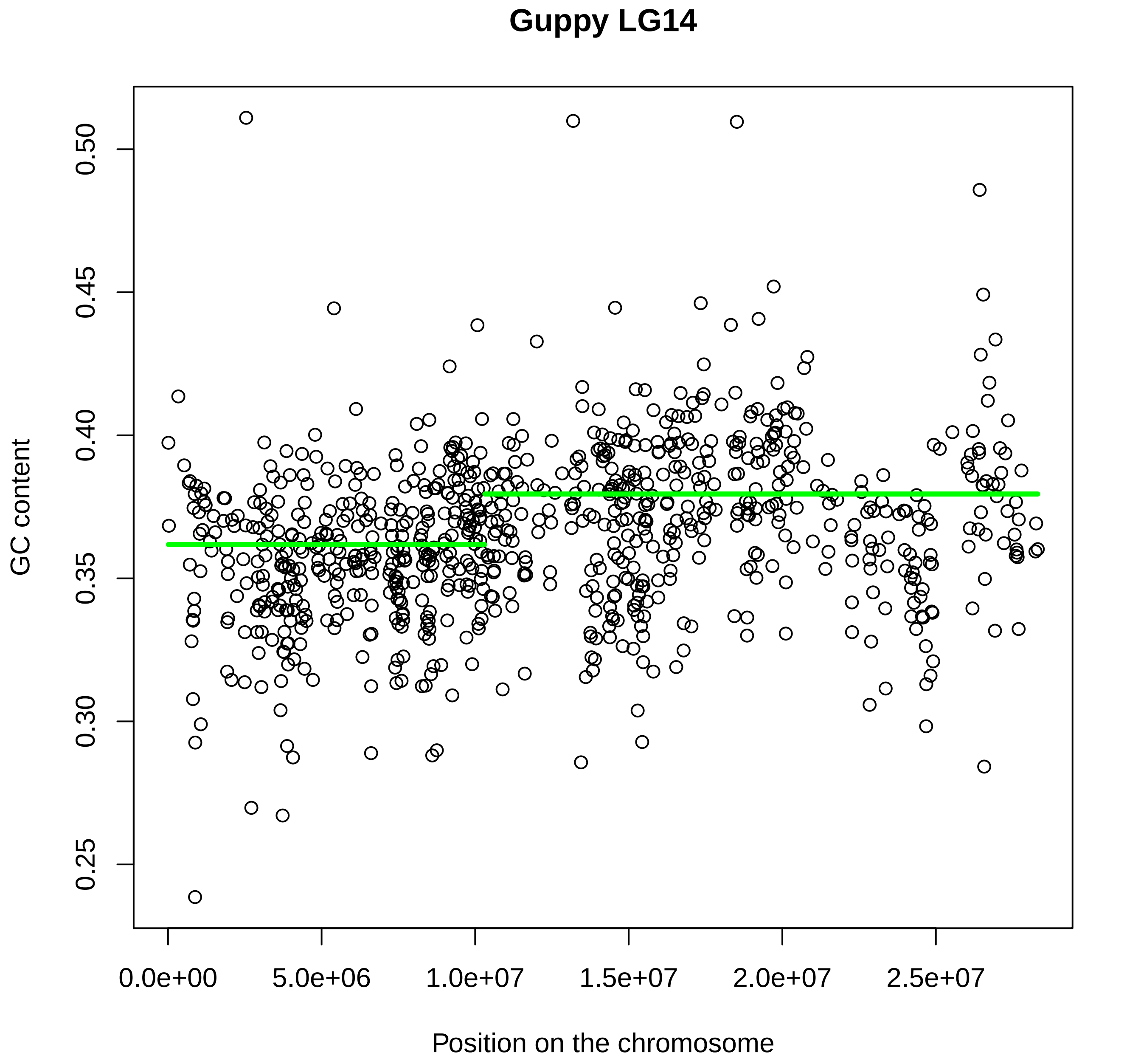

Guppy LG14
GC content
0.25
0.30
0.35
0.40
0.45
0.50
0.0e+00
5.0e+06
1.0e+07
1.5e+07
2.0e+07
2.5e+07
Position on the chromosome

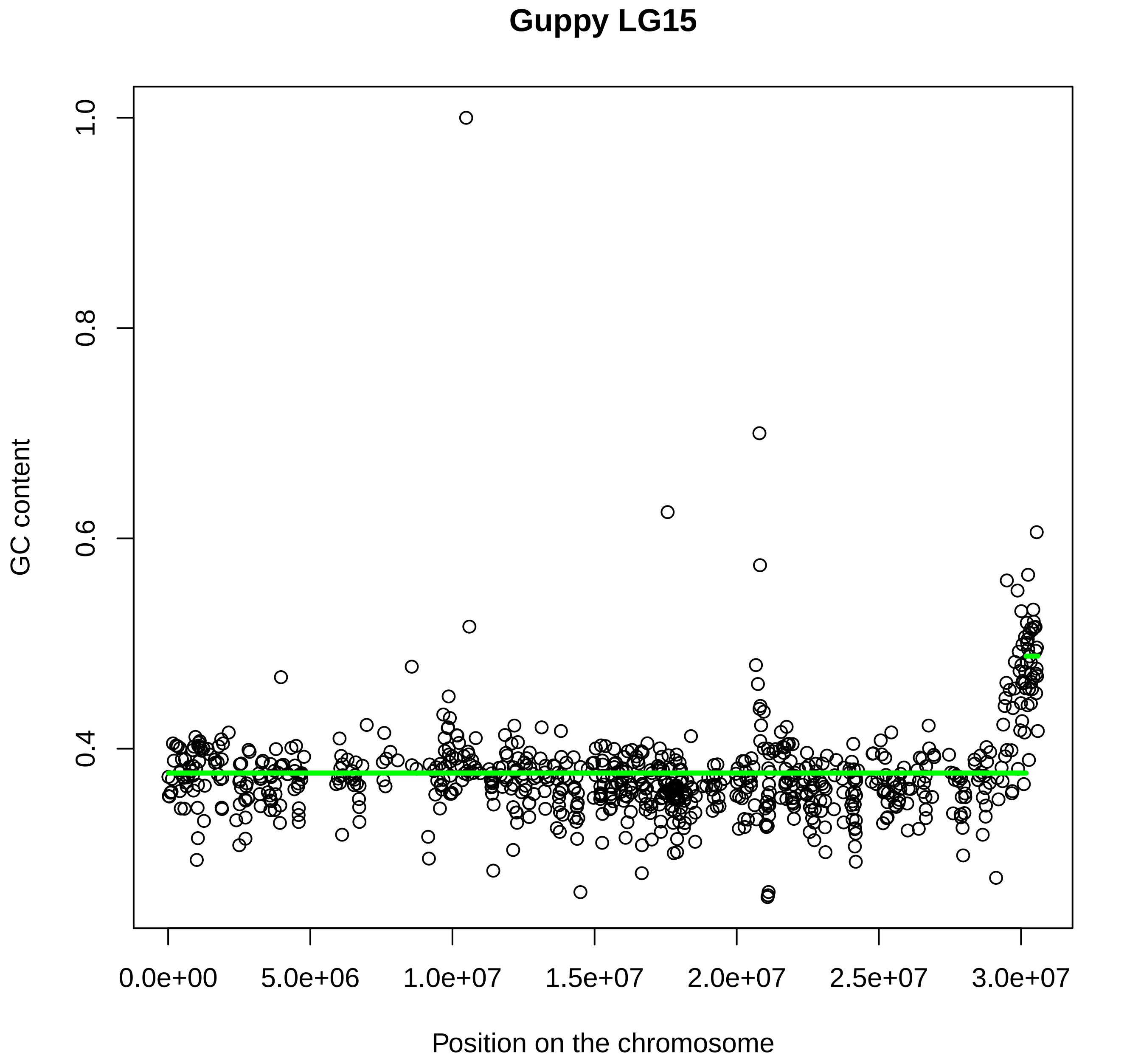

Guppy LG15
GC content
1.0
0.8
0.6
0.4
0.0e+00
5.0e+06
1.0e+07
1.5e+07
2.0e+07
2.5e+07
3.0e+07
Position on the chromosome

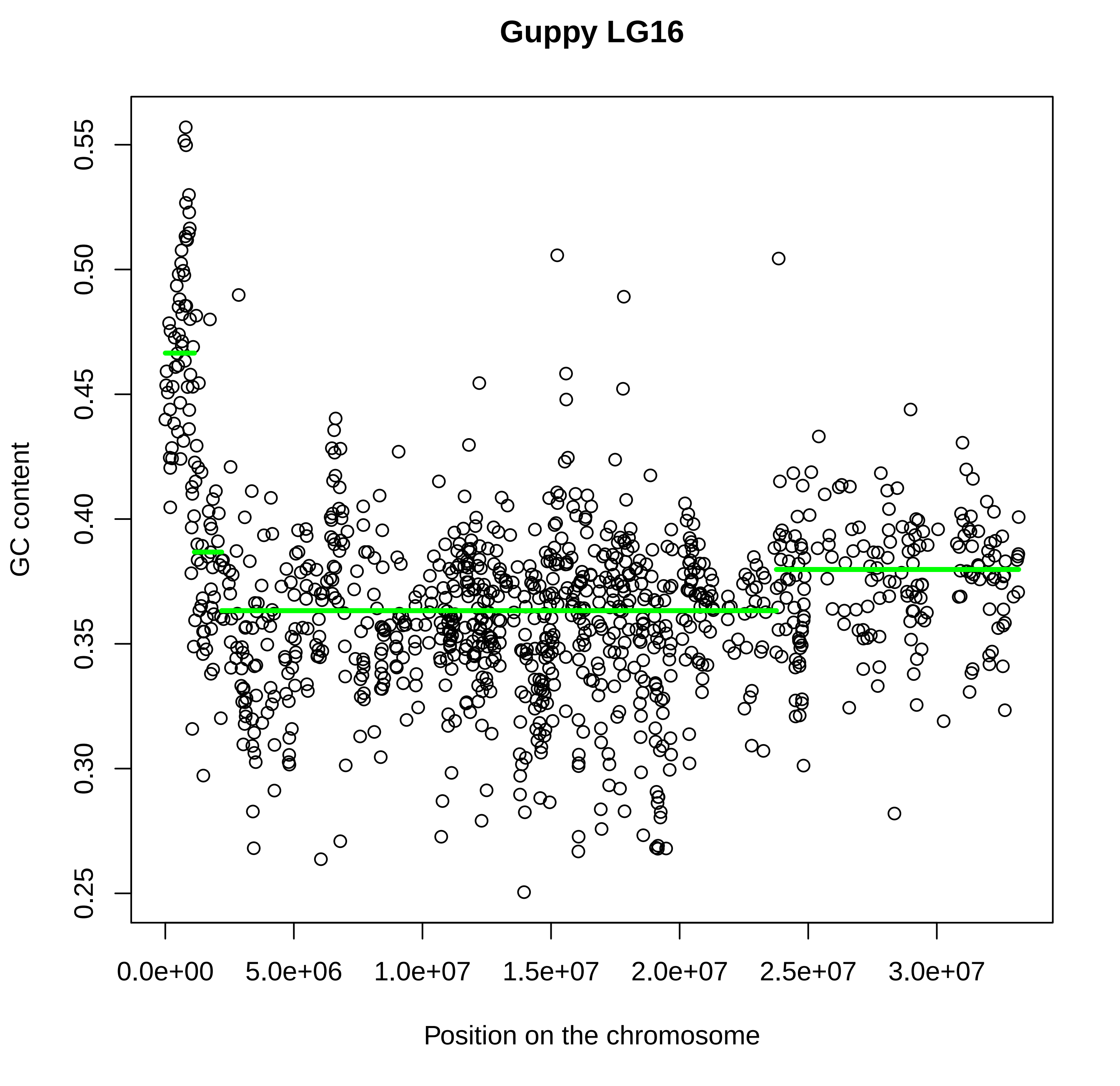

Guppy LG16
GC content
0.25
0.30
0.35
0.40
0.45
0.50
0.55
Position on the chromosome
0.0e+00
5.0e+06
1.0e+07
1.5e+07
2.0e+07
2.5e+07
3.0e+07

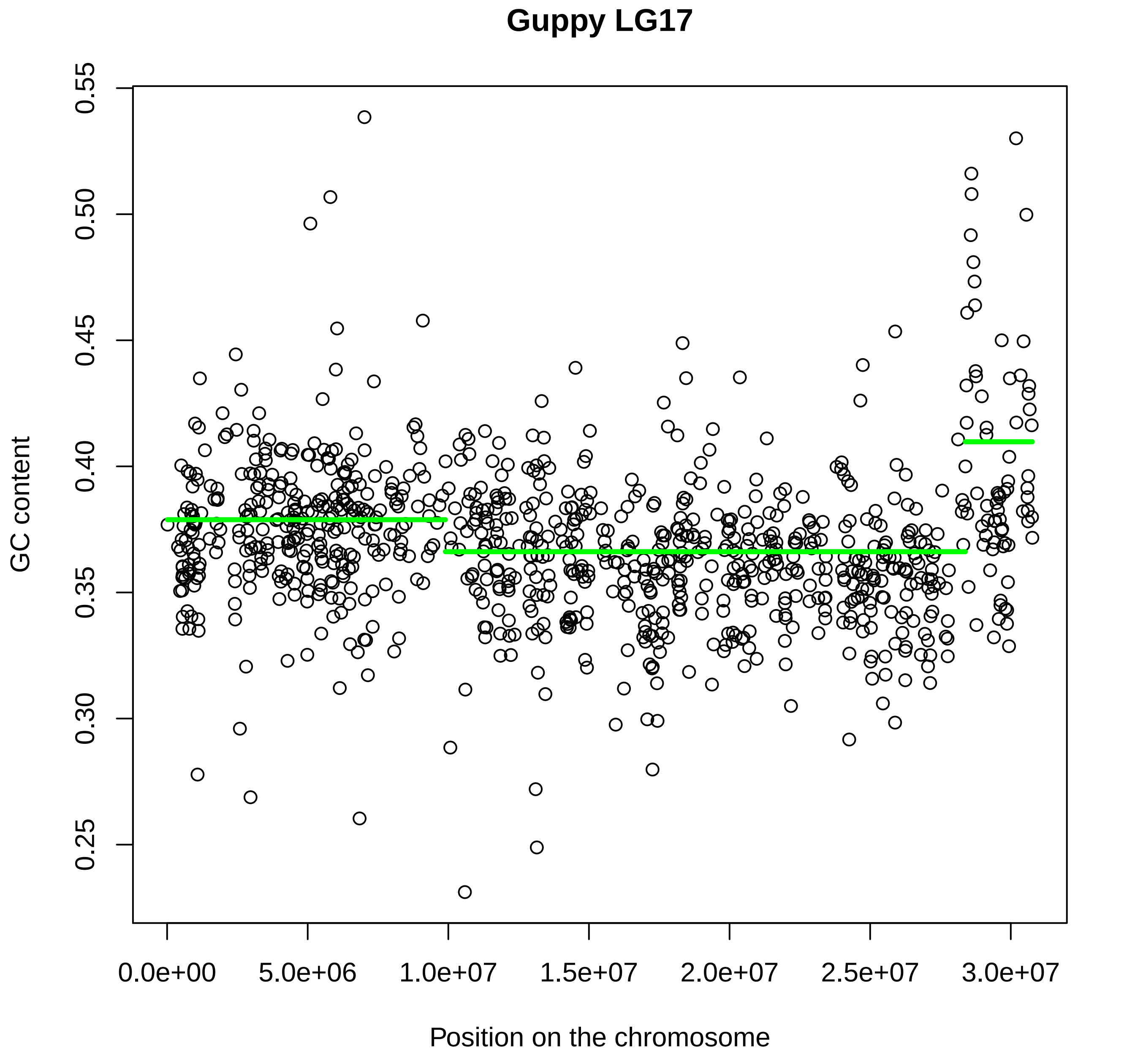

Guppy LG17
GC content
0.25
0.30
0.35
0.40
0.45
0.50
0.55
Position on the chromosome
0.0e+00
5.0e+06
1.0e+07
1.5e+07
2.0e+07
2.5e+07
3.0e+07

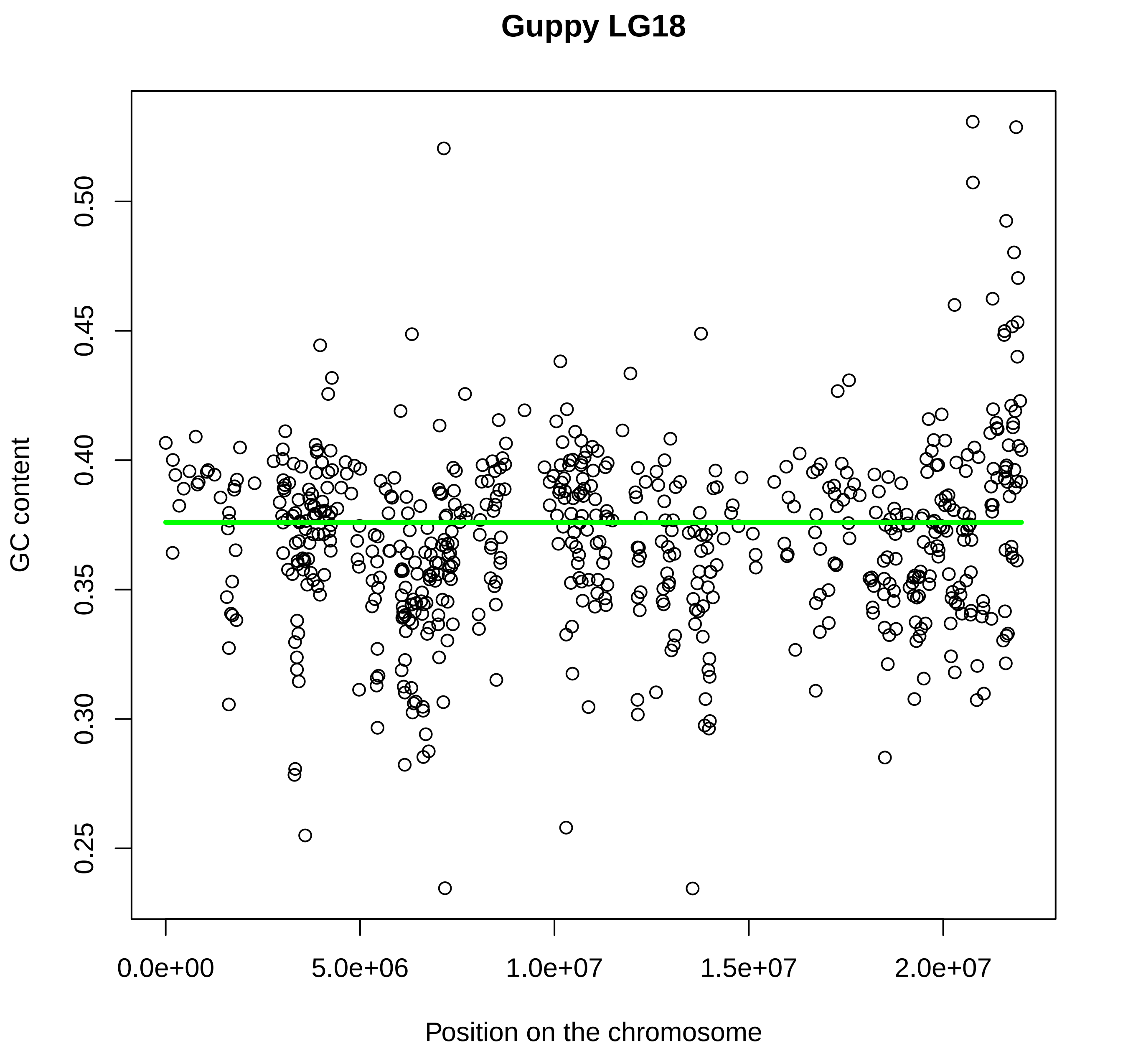
Guppy LG18
GC content
0.25
0.30
0.35
0.40
0.45
0.50
Position on the chromosome
0.0e+00
5.0e+06
1.0e+07
1.5e+07
2.0e+07

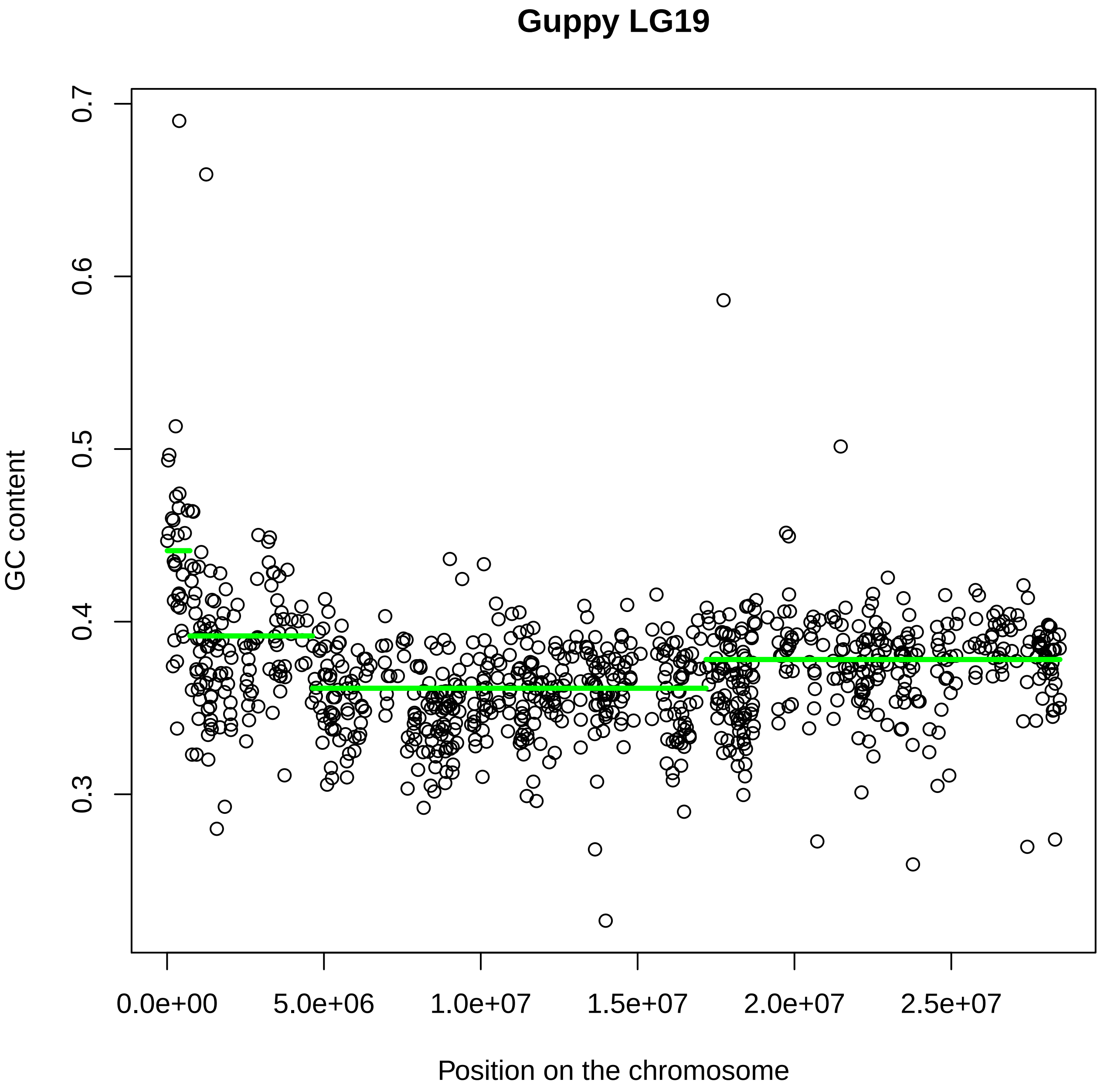

Guppy LG19
GC content
0.3
0.4
0.5
0.6
0.7
0.0e+00
5.0e+06
1.0e+07
1.5e+07
2.0e+07
2.5e+07
Position on the chromosome

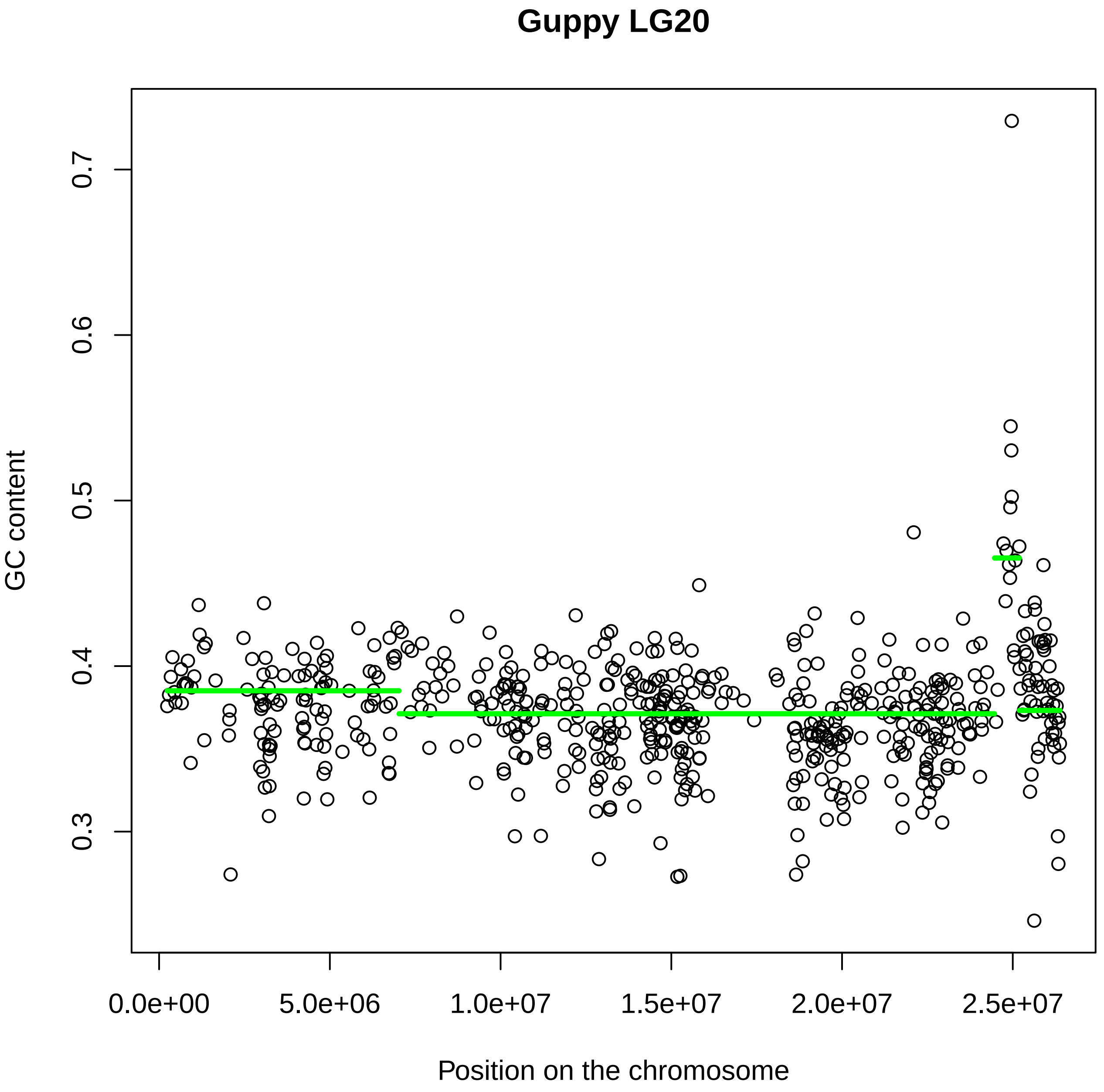
Guppy LG20
GC content
0.3
0.4
0.5
0.6
0.7
0.0e+00
5.0e+06
1.0e+07
1.5e+07
2.0e+07
2.5e+07
Position on the chromosome

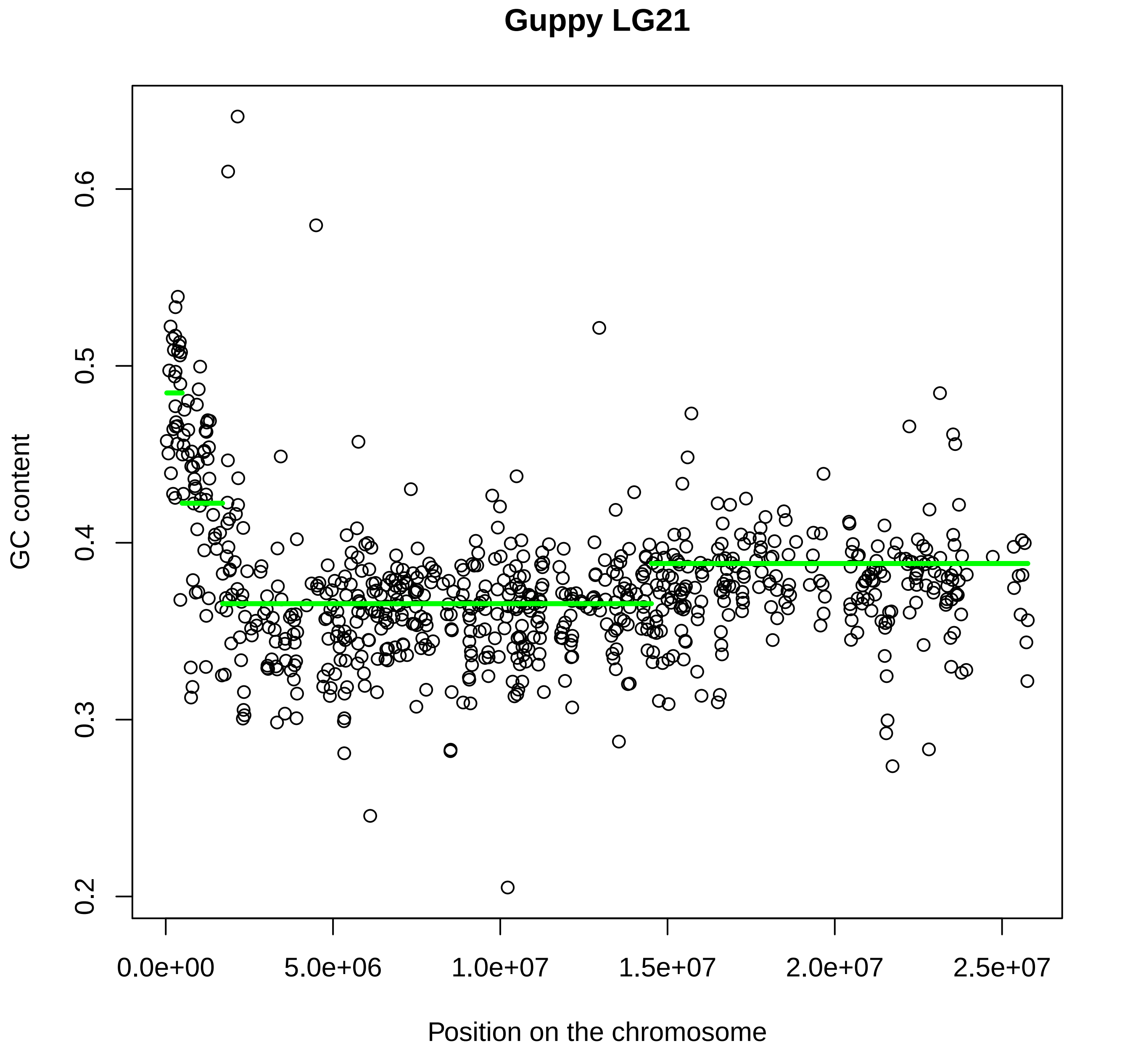
Guppy LG21
GC content
Position on the chromosome
0.2
0.3
0.4
0.5
0.6
0.0e+00
5.0e+06
1.0e+07
1.5e+07
2.0e+07
2.5e+07

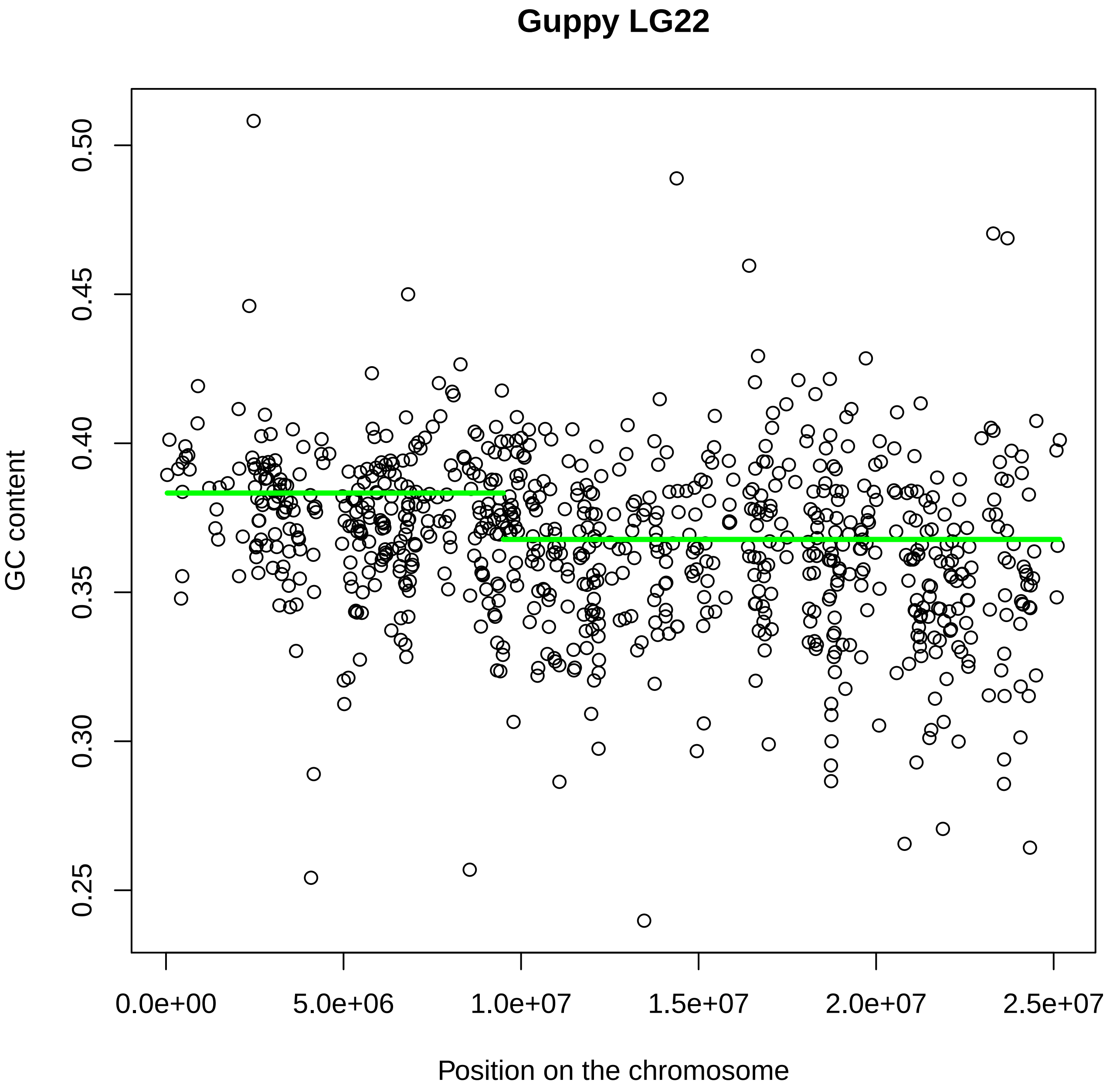

Guppy LG22
GC content
0.25
0.30
0.35
0.40
0.45
0.50
0.0e+00
5.0e+06
1.0e+07
1.5e+07
2.0e+07
2.5e+07
Position on the chromosome

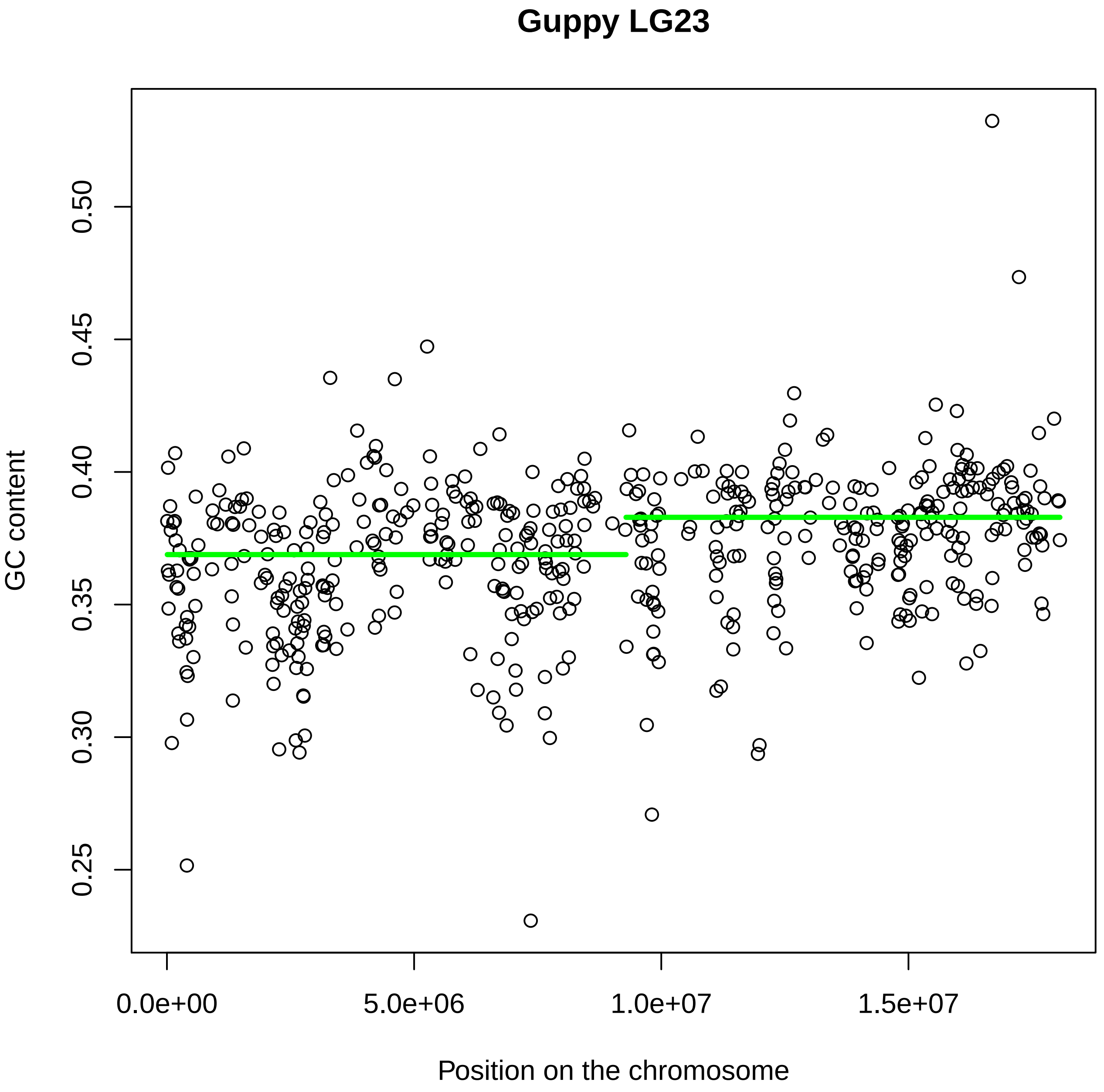

Guppy LG23
GC content
0.25
0.30
0.35
0.40
0.45
0.50
0.0e+00
5.0e+06
1.0e+07
1.5e+07
Position on the chromosome
