## Supplementary material for "Using GC content to compare recombination patterns on the sex chromosomes and autosomes of the guppy, *Poecilia reticulata*, and its close outgroup species": Figure S3 new.pdf

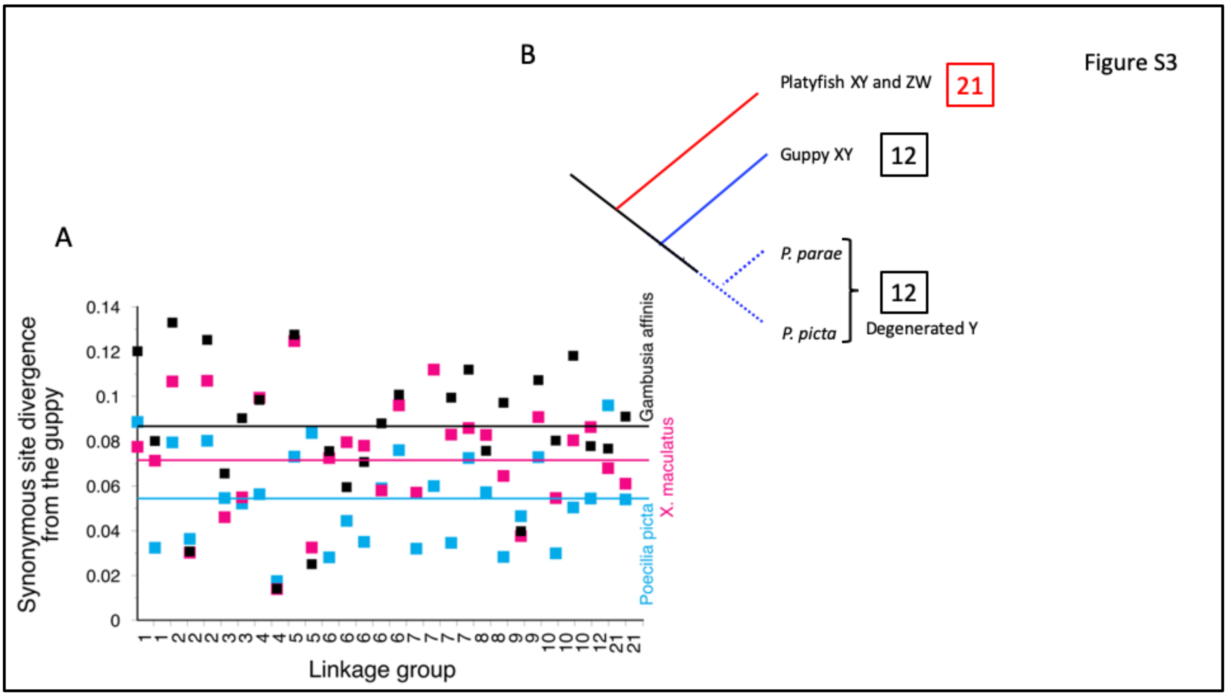


**Supplementary Figure S3.** Relationships between the species used in GC intron analyses. A. Synonymous site divergence between the guppy versus the platyfish, *Xiphophorus maculatus*, and *P. picta*. B. Schematic diagram of the relationships based on relative synonymous site divergence values, to show that *P. picta* is a closer outgroup than the platyfish.
