## Supplementary material for "Using GC content to compare recombination patterns on the sex chromosomes and autosomes of the guppy, *Poecilia reticulata*, and its close outgroup species": Figure S4.pdf

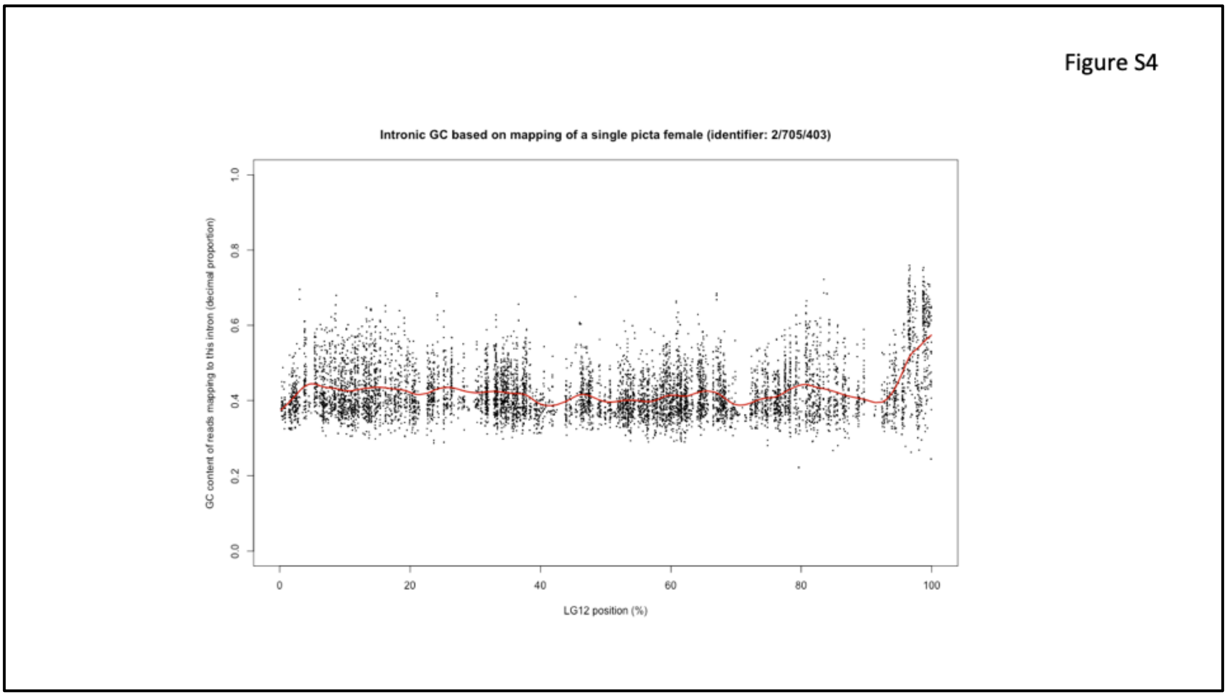


**Supplementary Figure S4.** GC content of introns in *P. picta* genes with homologs on the guppy LG12. Because the *P. picta* assembly is not contiguous, each dot represents an individual intron. The pattern is therefore less clear than for the guppy, where we pooled introns for each gene (see Figure S2). The red line shows smoothed GC content values (based on a smooth spline with 1/250 the maximum degrees of freedom for each chromosome).
