## Supplementary material for "Using GC content to compare recombination patterns on the sex chromosomes and autosomes of the guppy, *Poecilia reticulata*, and its close outgroup species": Figure S5 (GYks_CPA)_NEW.pdf

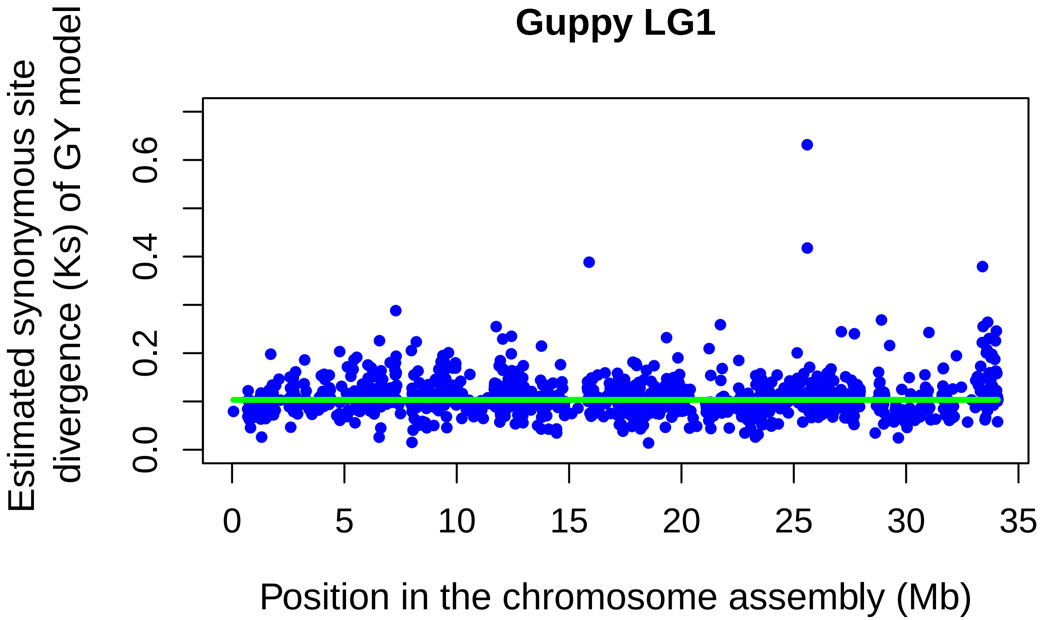
Guppy LG1
Estimated synonymous site divergence (Ks) of GY model
0.0
0.2
0.4
0.6
0
5
10
15
20
25
30
35
Position in the chromosome assembly (Mb)

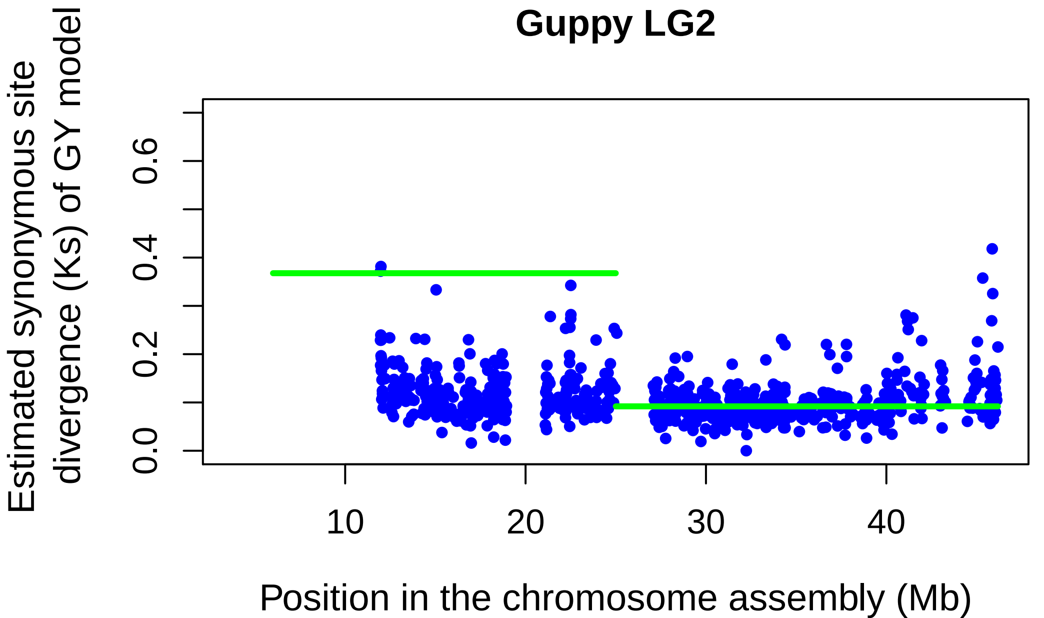
Guppy LG2
Estimated synonymous site divergence (Ks) of GY model
0.0
0.2
0.4
0.6
10
20
30
40
Position in the chromosome assembly (Mb)

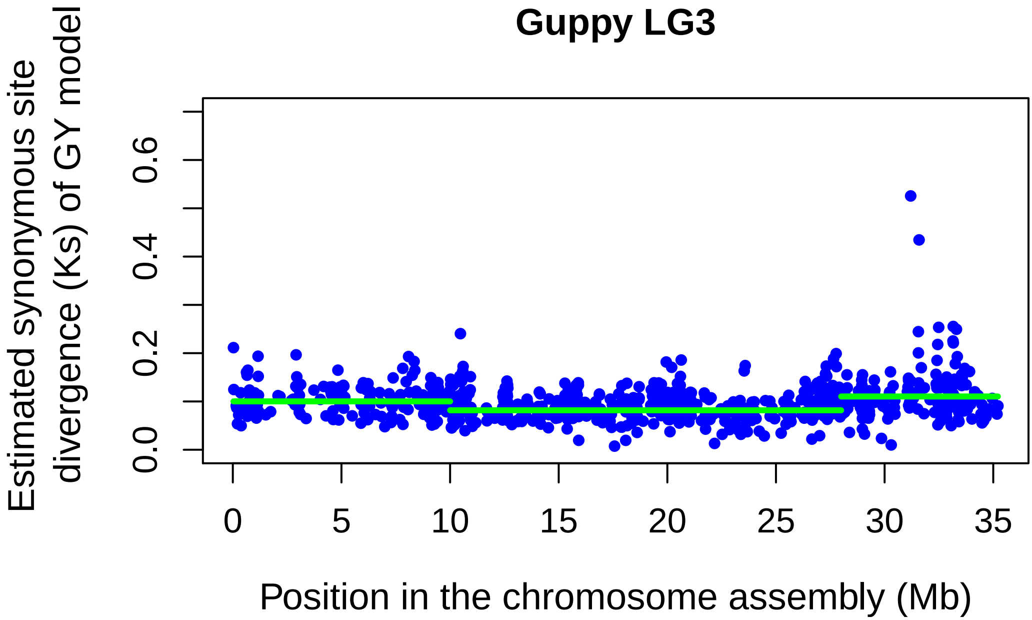

Guppy LG3
Estimated synonymous site divergence (Ks) of GY model
0.0
0.2
0.4
0.6
0
5
10
15
20
25
30
35
Position in the chromosome assembly (Mb)


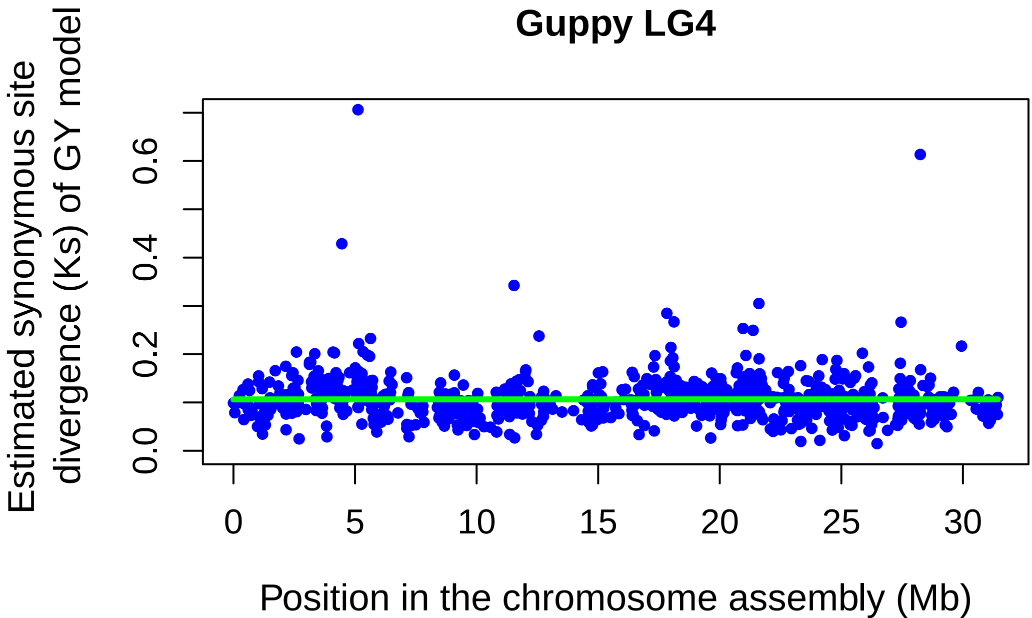

Guppy LG4
Estimated synonymous site divergence (Ks) of GY model
0.0
0.2
0.4
0.6
0
5
10
15
20
25
30
Position in the chromosome assembly (Mb)

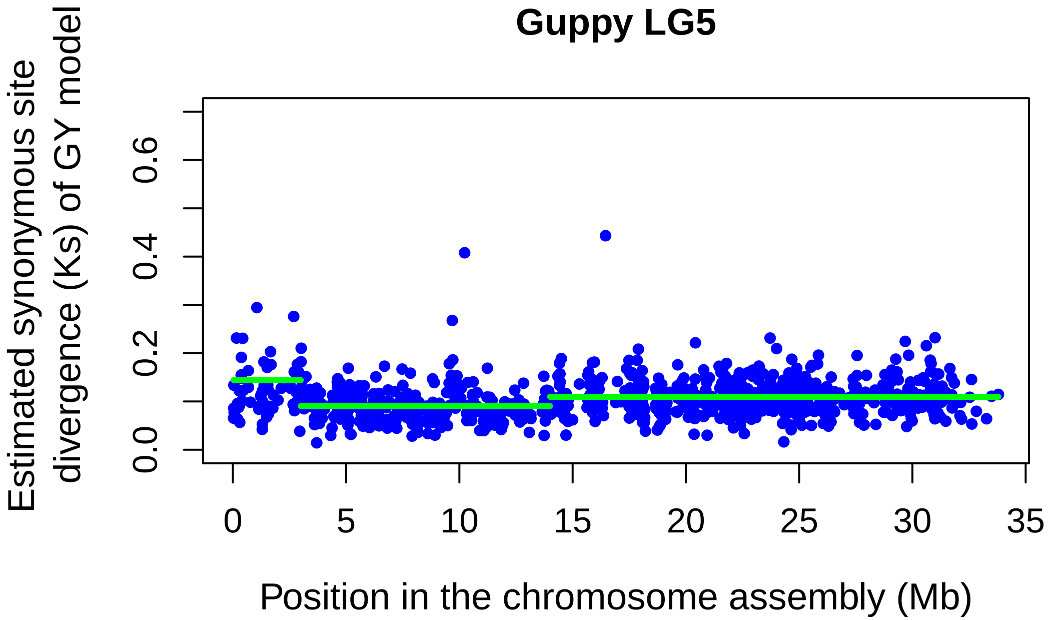
Guppy LG5
Estimated synonymous site divergence (Ks) of GY model
0.0
0.2
0.4
0.6
0
5
10
15
20
25
30
35
Position in the chromosome assembly (Mb)

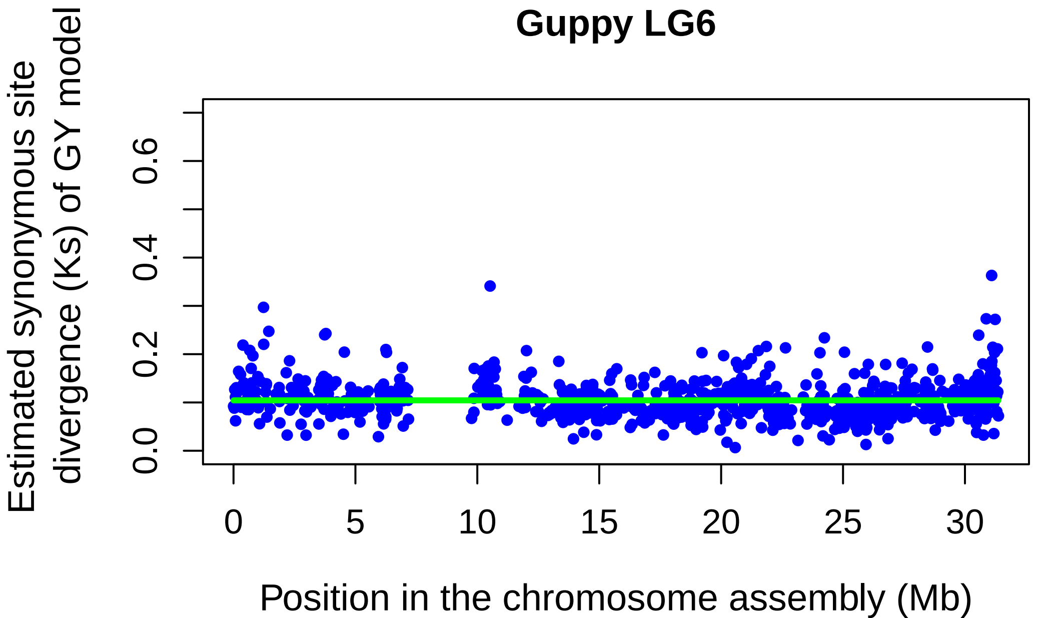
Guppy LG6
Estimated synonymous site divergence (Ks) of GY model
0.0
0.2
0.4
0.6
0
5
10
15
20
25
30
Position in the chromosome assembly (Mb)

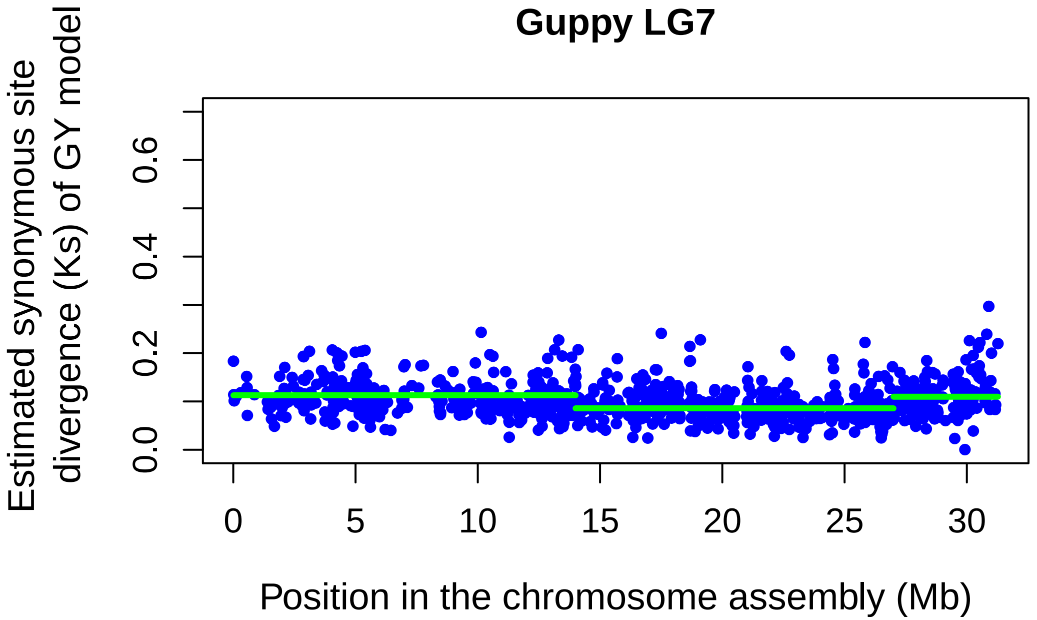
Guppy LG7
Estimated synonymous site divergence (Ks) of GY model
0.0
0.2
0.4
0.6
0
5
10
15
20
25
30
Position in the chromosome assembly (Mb)

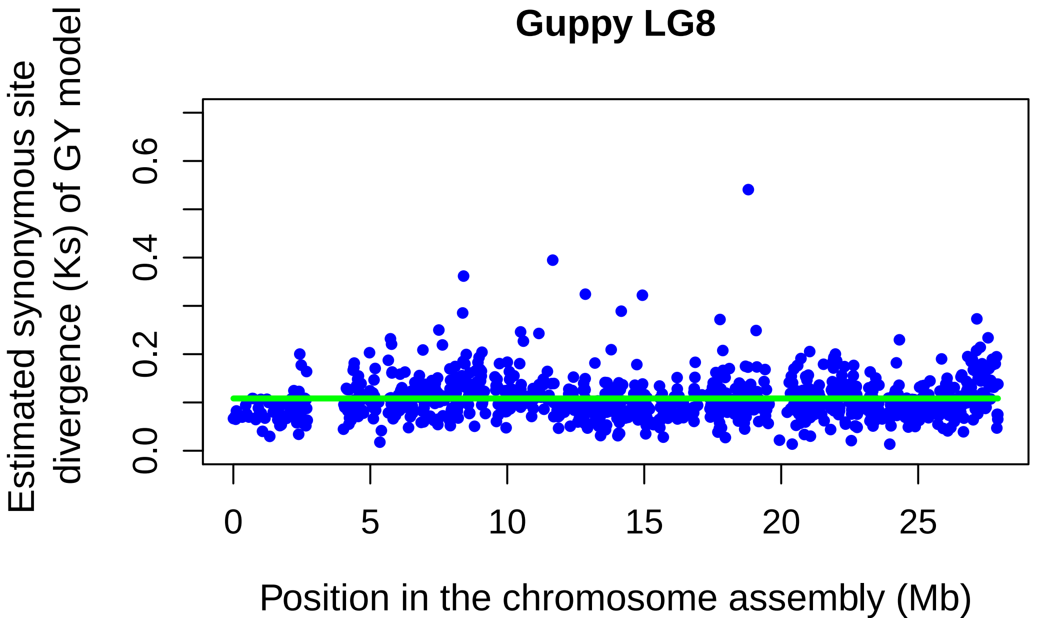
Guppy LG8
Estimated synonymous site divergence (Ks) of GY model
0.0
0.2
0.4
0.6
0
5
10
15
20
25
Position in the chromosome assembly (Mb)

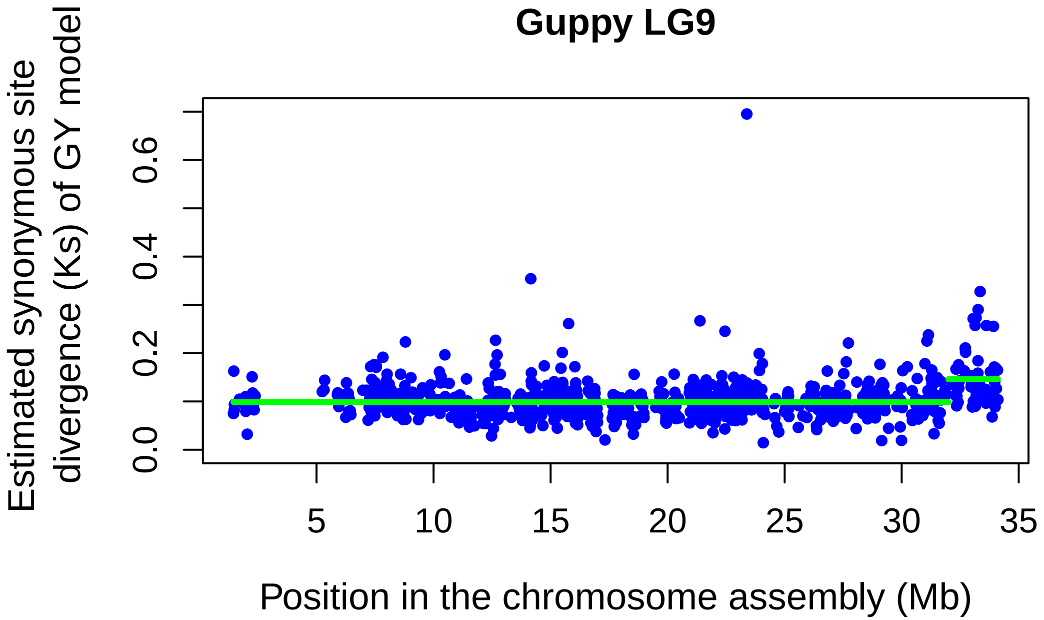
Guppy LG9
Estimated synonymous site divergence (Ks) of GY model
0.0
0.2
0.4
0.6
5
10
15
20
25
30
35
Position in the chromosome assembly (Mb)

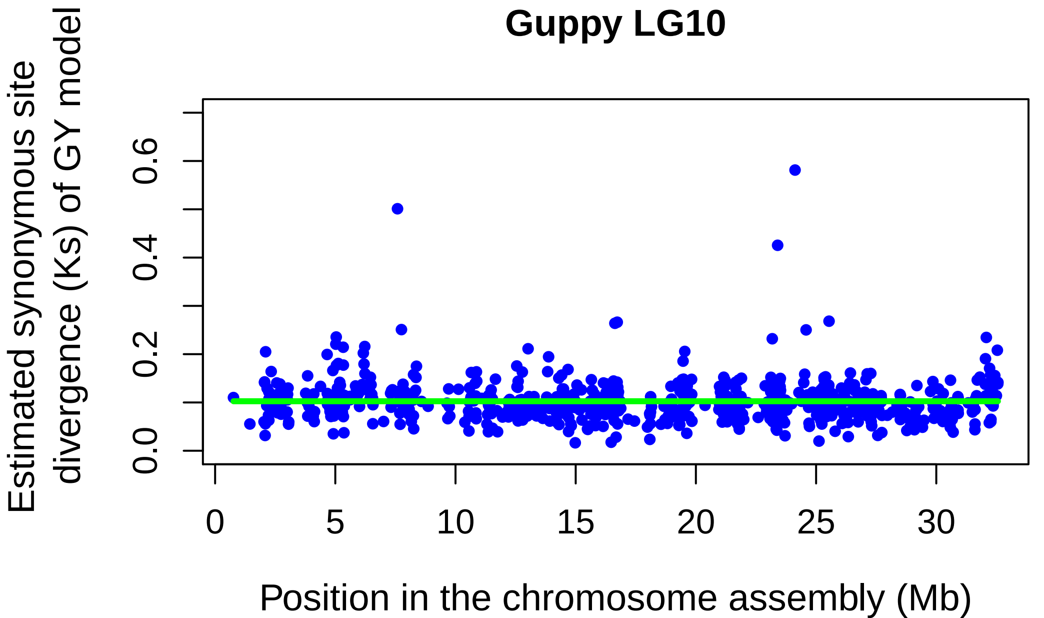
Guppy LG10
Estimated synonymous site divergence (Ks) of GY model
0.0
0.2
0.4
0.6
0
5
10
15
20
25
30
Position in the chromosome assembly (Mb)

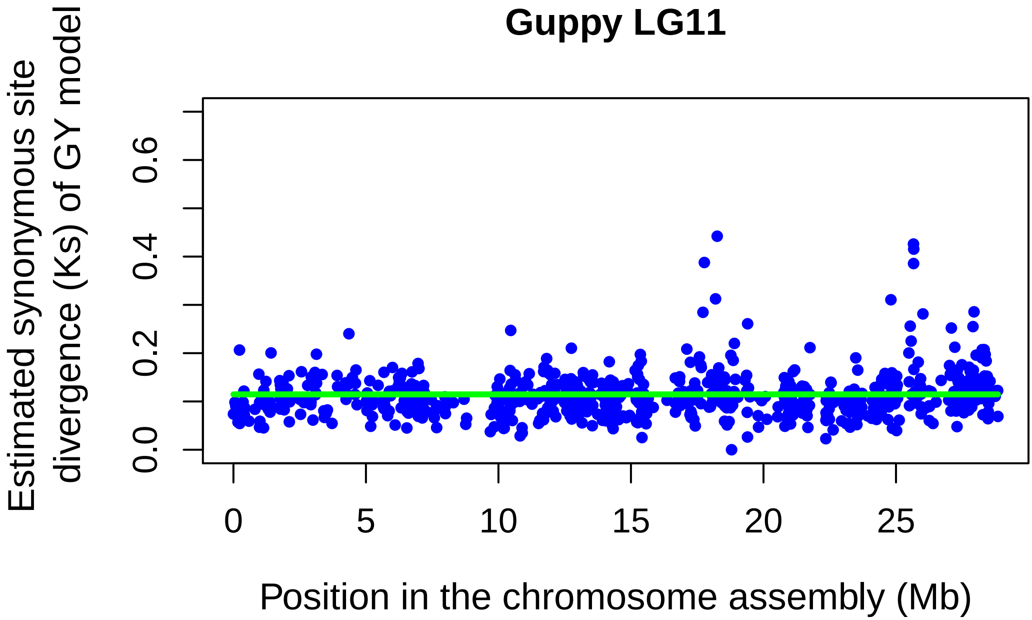
Guppy LG11
Estimated synonymous site divergence (Ks) of GY model
0.0
0.2
0.4
0.6
0
5
10
15
20
25
Position in the chromosome assembly (Mb)

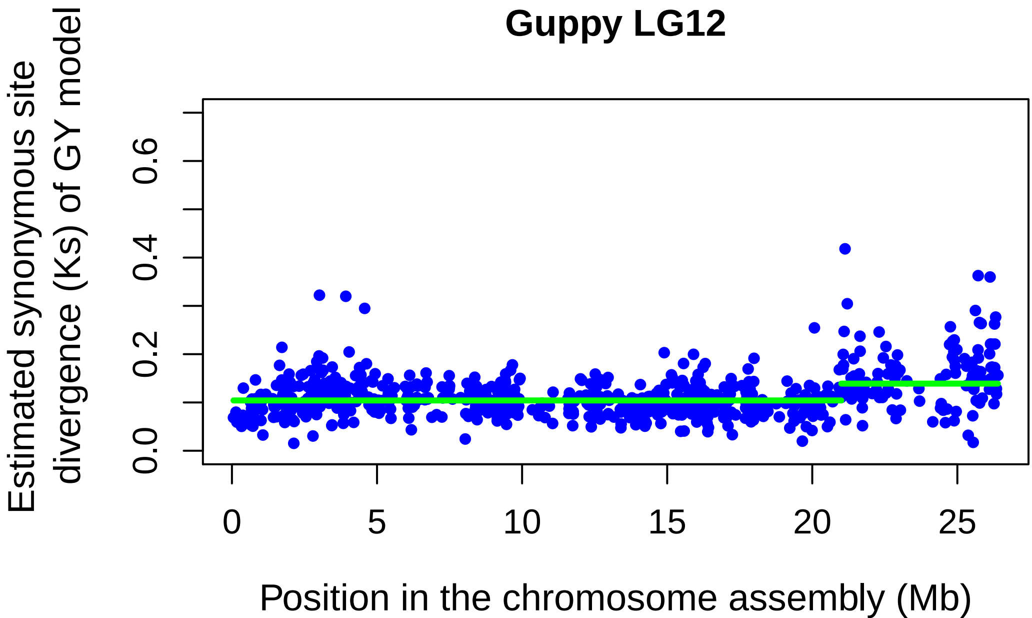
Guppy LG12
Estimated synonymous site divergence (Ks) of GY model
0.0
0.2
0.4
0.6
0
5
10
15
20
25
Position in the chromosome assembly (Mb)

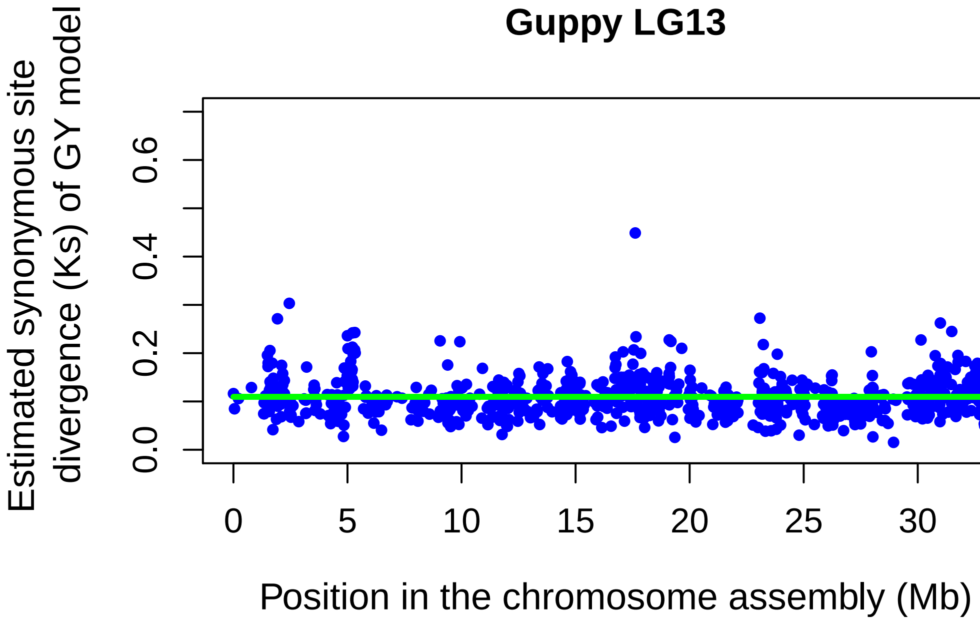
Guppy LG13
Estimated synonymous site
divergence (Ks) of GY model
0.0
0.2
0.4
0.6
0
5
10
15
20
25
30
Position in the chromosome assembly (Mb)

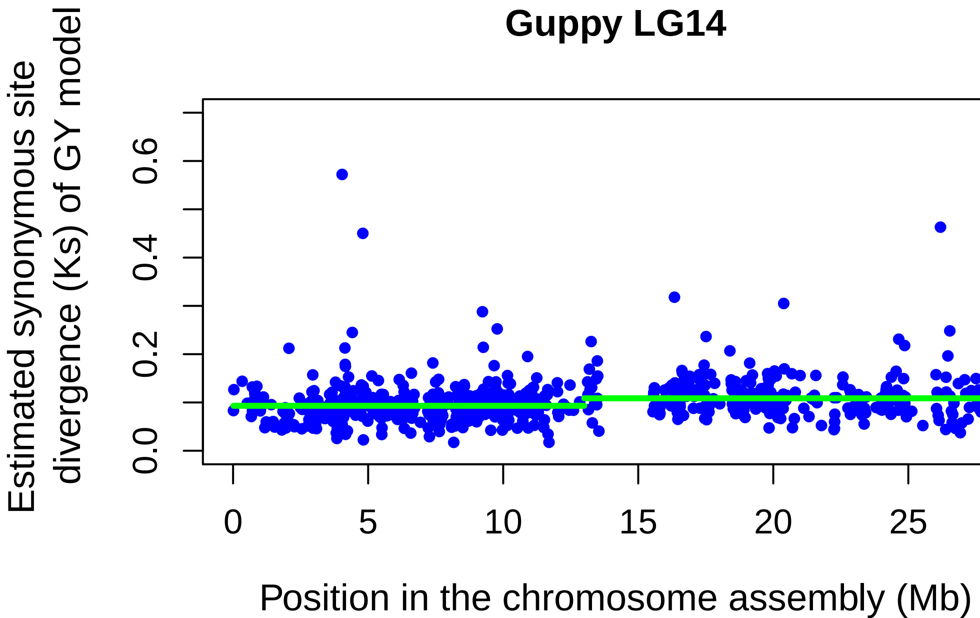
Guppy LG14
Estimated synonymous site
divergence (Ks) of GY model
0.0
0.2
0.4
0.6
0
5
10
15
20
25
Position in the chromosome assembly (Mb)

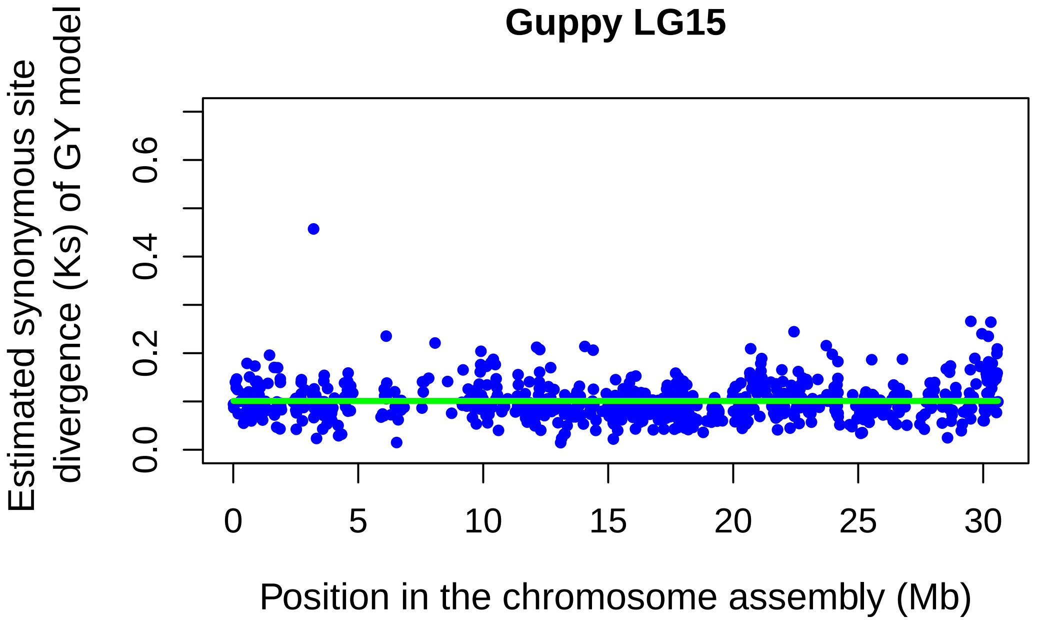
Guppy LG15
Estimated synonymous site divergence (Ks) of GY model
0.0
0.2
0.4
0.6
0
5
10
15
20
25
30
Position in the chromosome assembly (Mb)

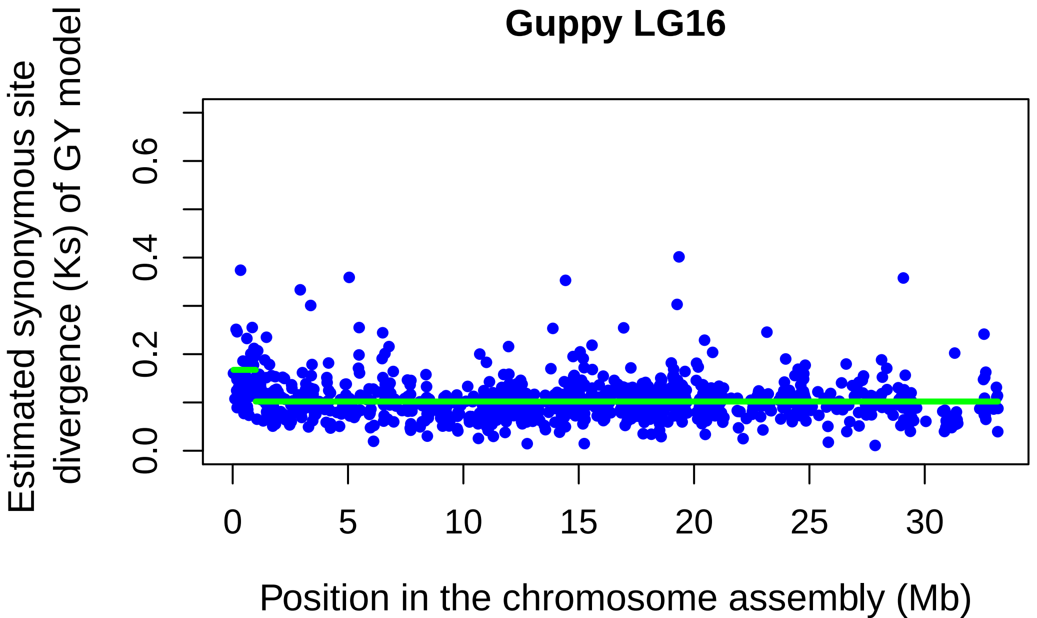
Guppy LG16
Estimated synonymous site divergence (Ks) of GY model
0.0
0.2
0.4
0.6
0
5
10
15
20
25
30
Position in the chromosome assembly (Mb)

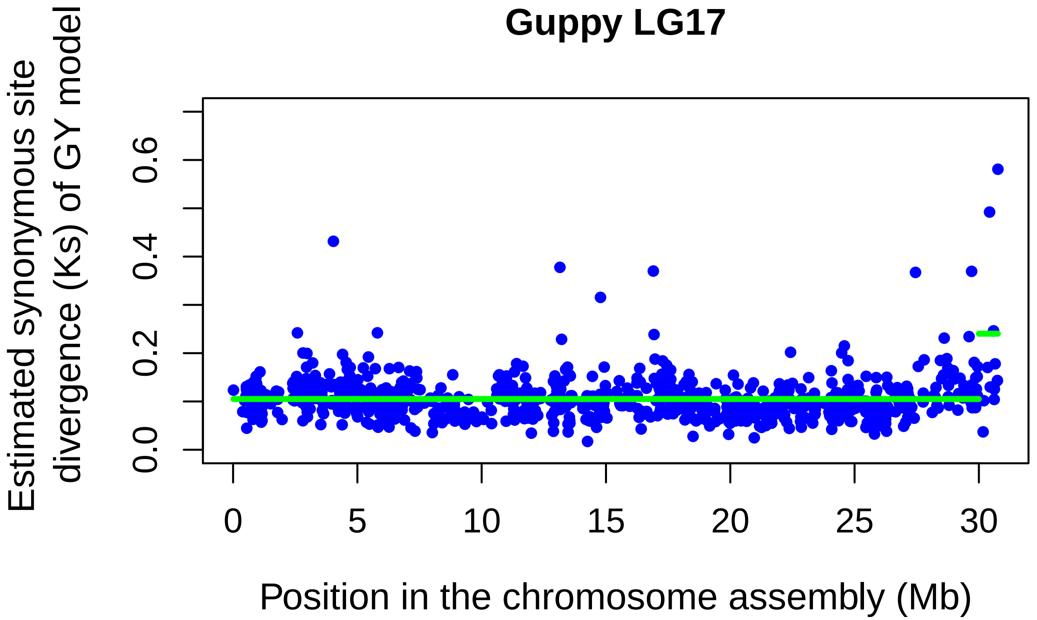
Guppy LG17
Estimated synonymous site divergence (Ks) of GY model
0.0
0.2
0.4
0.6
0
5
10
15
20
25
30
Position in the chromosome assembly (Mb)

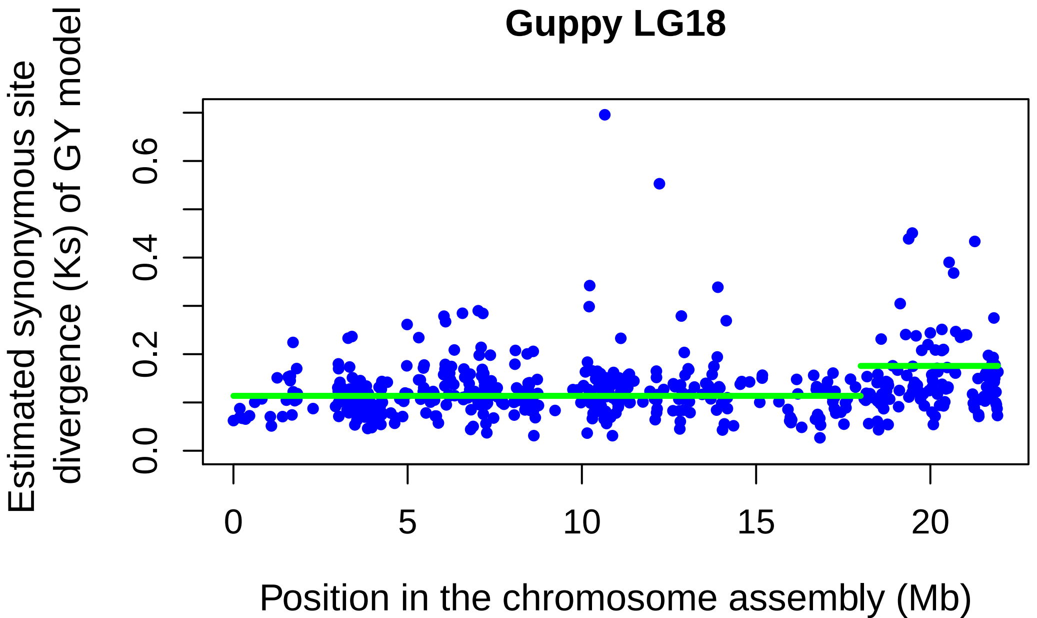
Guppy LG18
Estimated synonymous site divergence (Ks) of GY model
0.0
0.2
0.4
0.6
0
5
10
15
20
Position in the chromosome assembly (Mb)

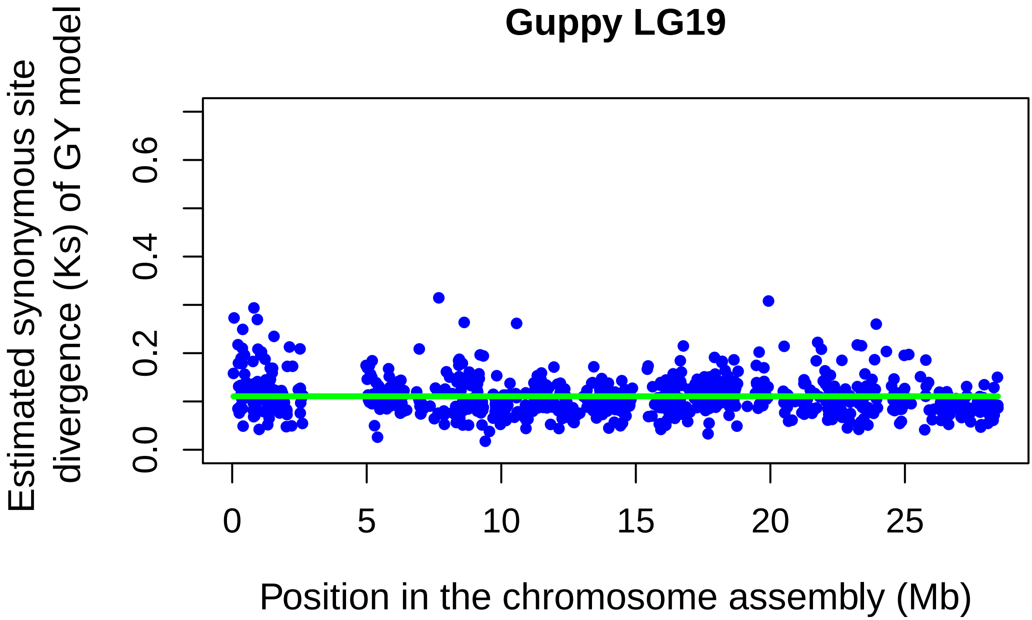
Guppy LG19
Estimated synonymous site divergence (Ks) of GY model
0.0
0.2
0.4
0.6
0
5
10
15
20
25
Position in the chromosome assembly (Mb)

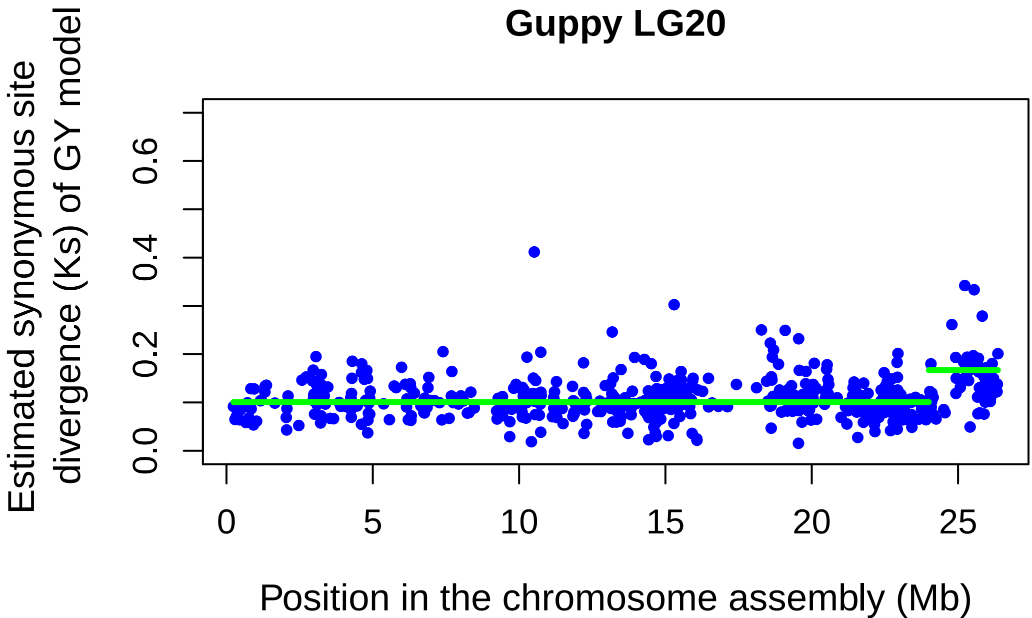
Guppy LG20
Estimated synonymous site divergence (Ks) of GY model
0.0
0.2
0.4
0.6
0
5
10
15
20
25
Position in the chromosome assembly (Mb)

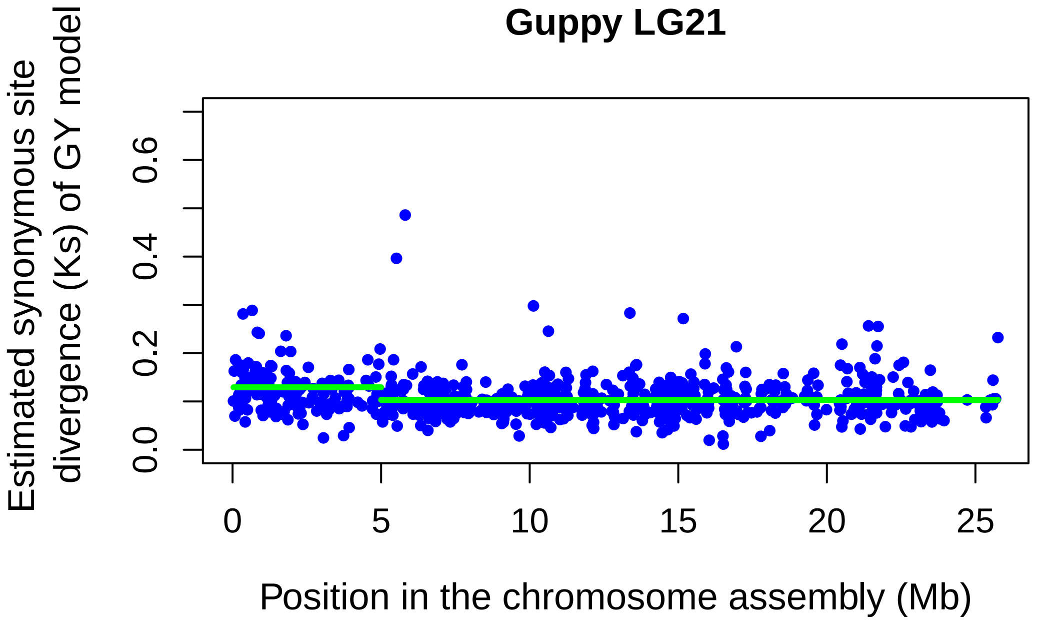

Guppy LG21
Estimated synonymous site divergence (Ks) of GY model
0.0
0.2
0.4
0.6
0
5
10
15
20
25
Position in the chromosome assembly (Mb)


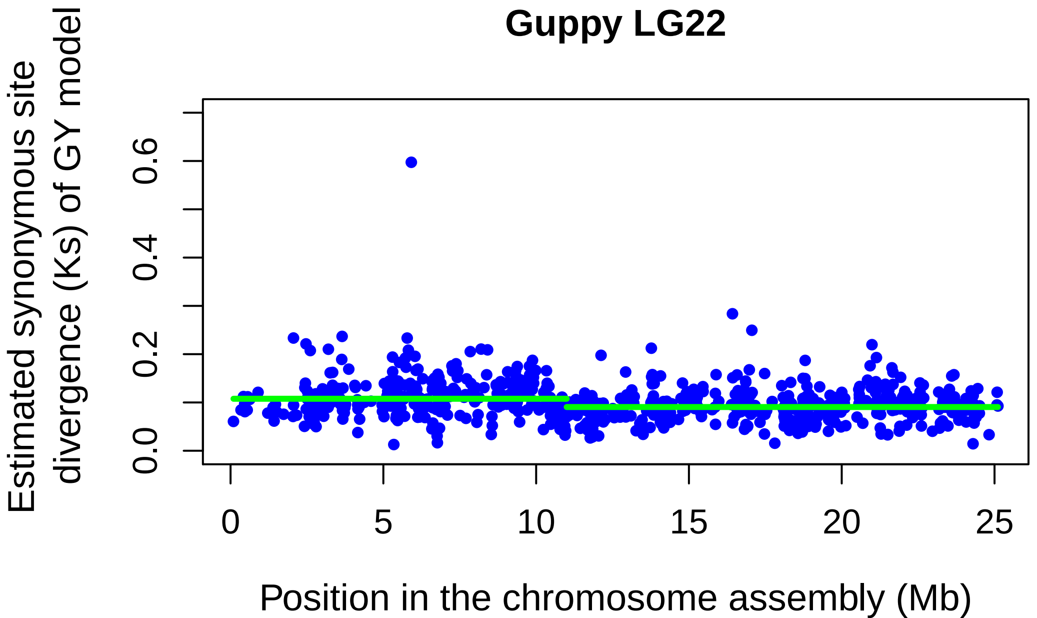

Guppy LG22
Estimated synonymous site divergence (Ks) of GY model
0.0
0.2
0.4
0.6
0
5
10
15
20
25
Position in the chromosome assembly (Mb)

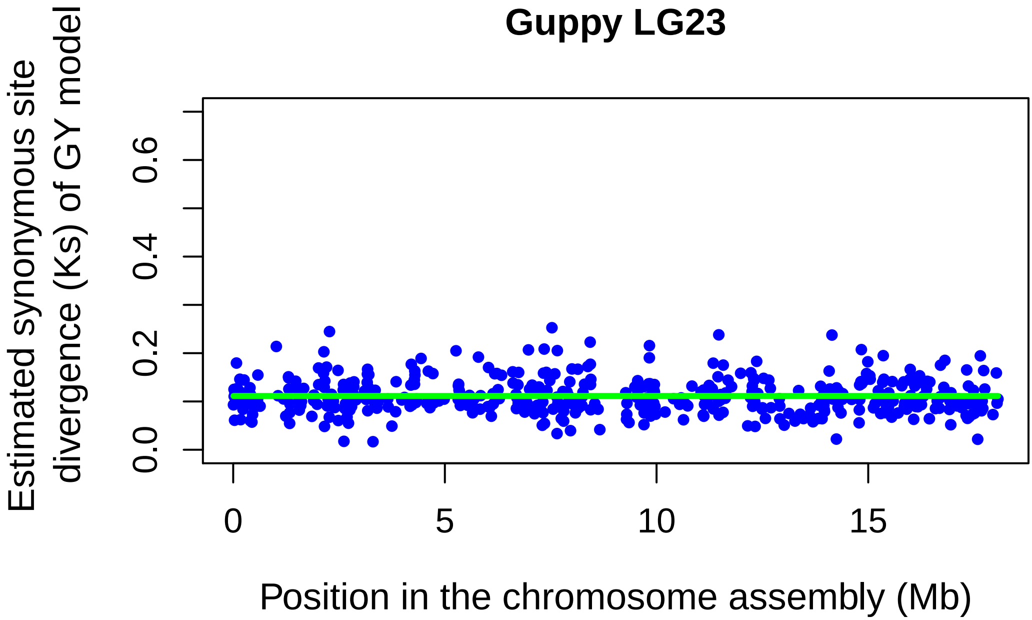
Guppy LG23
Estimated synonymous site divergence (Ks) of GY model
0.0
0.2
0.4
0.6
0
5
10
15
Position in the chromosome assembly (Mb)

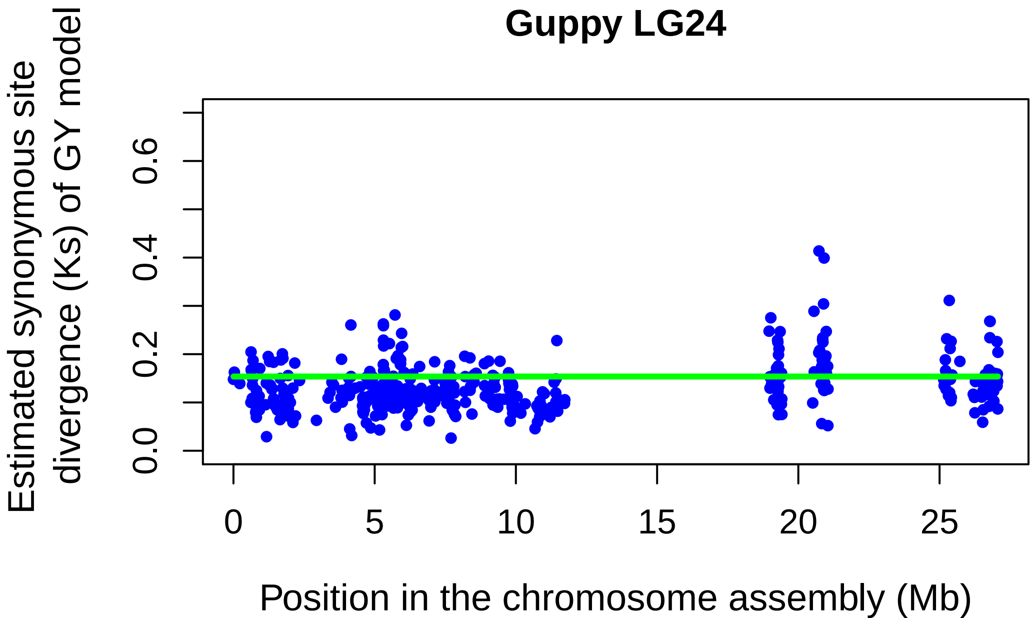
Guppy LG24
Estimated synonymous site divergence (Ks) of GY model
0.0
0.2
0.4
0.6
0
5
10
15
20
25
Position in the chromosome assembly (Mb)
