## Supplementary material for "Using GC content to compare recombination patterns on the sex chromosomes and autosomes of the guppy, *Poecilia reticulata*, and its close outgroup species": Figure S6 new.pdf

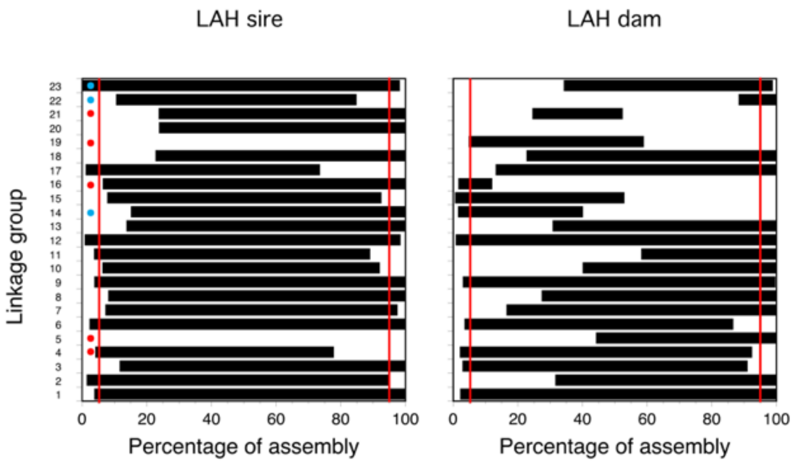


**Supplementary Figure S6.** Regions of each guppy chromosome with high-throughput mapping data informative in male meiosis (left) and meiosis of the dam (right). In the diagram for the sire meiosis, the red dots indicate the five guppy chromosomes whose left-hand assembly end has the GC spike, when a spike was detected, and blue dots indicate chromosomes for which no spike was detected; all other chromosomes had a spike at the right-hand end of the assembly, except that a middle region spike was also detected for the fusion chromosome, LG2.
