## Supplementary material for "Using GC content to compare recombination patterns on the sex chromosomes and autosomes of the guppy, *Poecilia reticulata*, and its close outgroup species": Figure S8.pdf

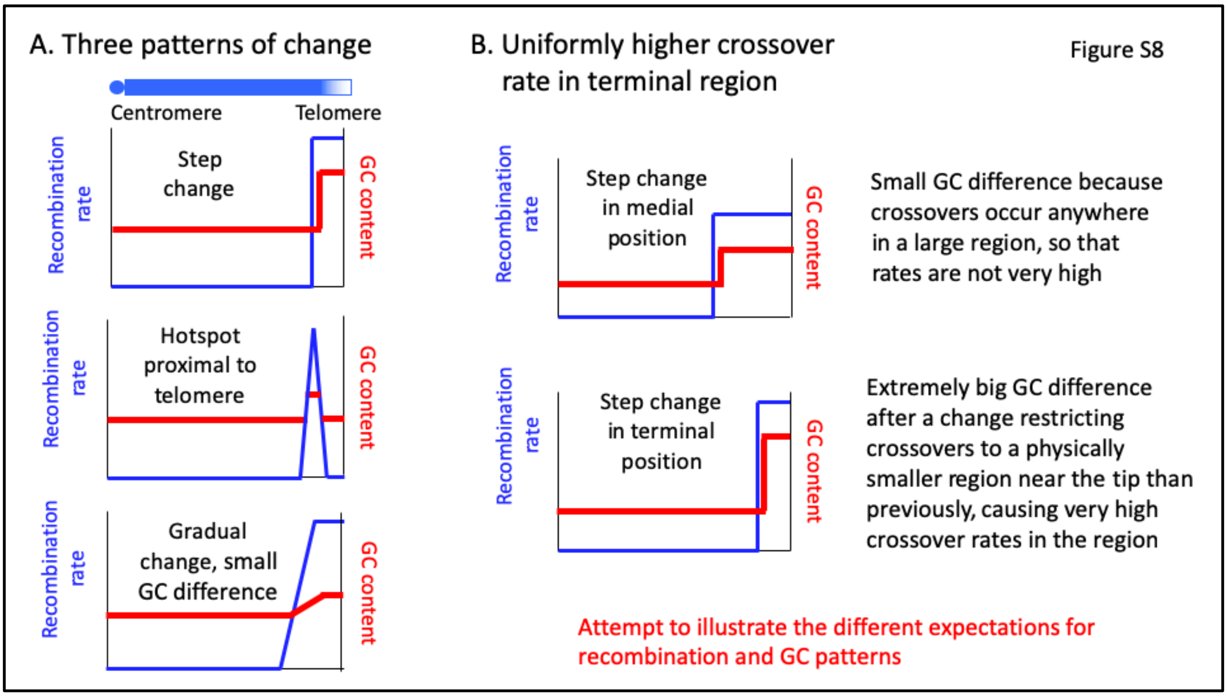


**Supplementary Figure S8.** Possible patterns of crossovers, and the expected GC patterns. A. Crossovers could occur uniformly across the region where they occur, or be localised. B. If crossovers occur uniformly across a large recombining region, the GC content will be only slightly higher in the region. If, however, they change to occur exclusively in a physically small region, the recombination rate will be very high, and a large increase in GC is expected.
