## Supplementary material for "Using GC content to compare recombination patterns on the sex chromosomes and autosomes of the guppy, *Poecilia reticulata*, and its close outgroup species": Figure S9.pdf

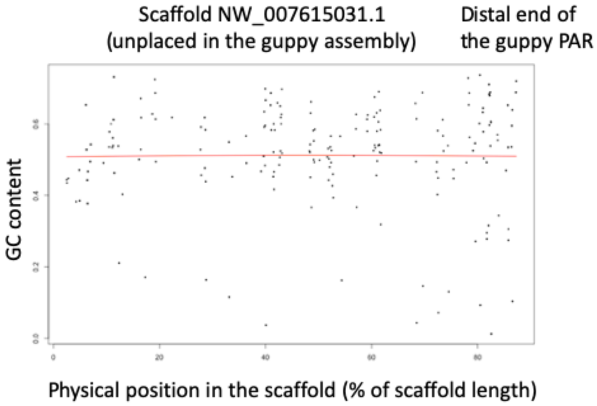


**Supplementary Figure S9.** High GC content in a scaffold that is unplaced in the guppy assembly, NW_007615031.1, but that we found to be located at the terminus of the guppy sex chromosome pair (unpublished result in Charlesworth et al. 2020).
