## Supplementary figures and images for "Using GC content to compare recombination patterns on the sex chromosomes and autosomes of the guppy, *Poecilia reticulata*, and its close outgroup species"

### Figure S1.pdf

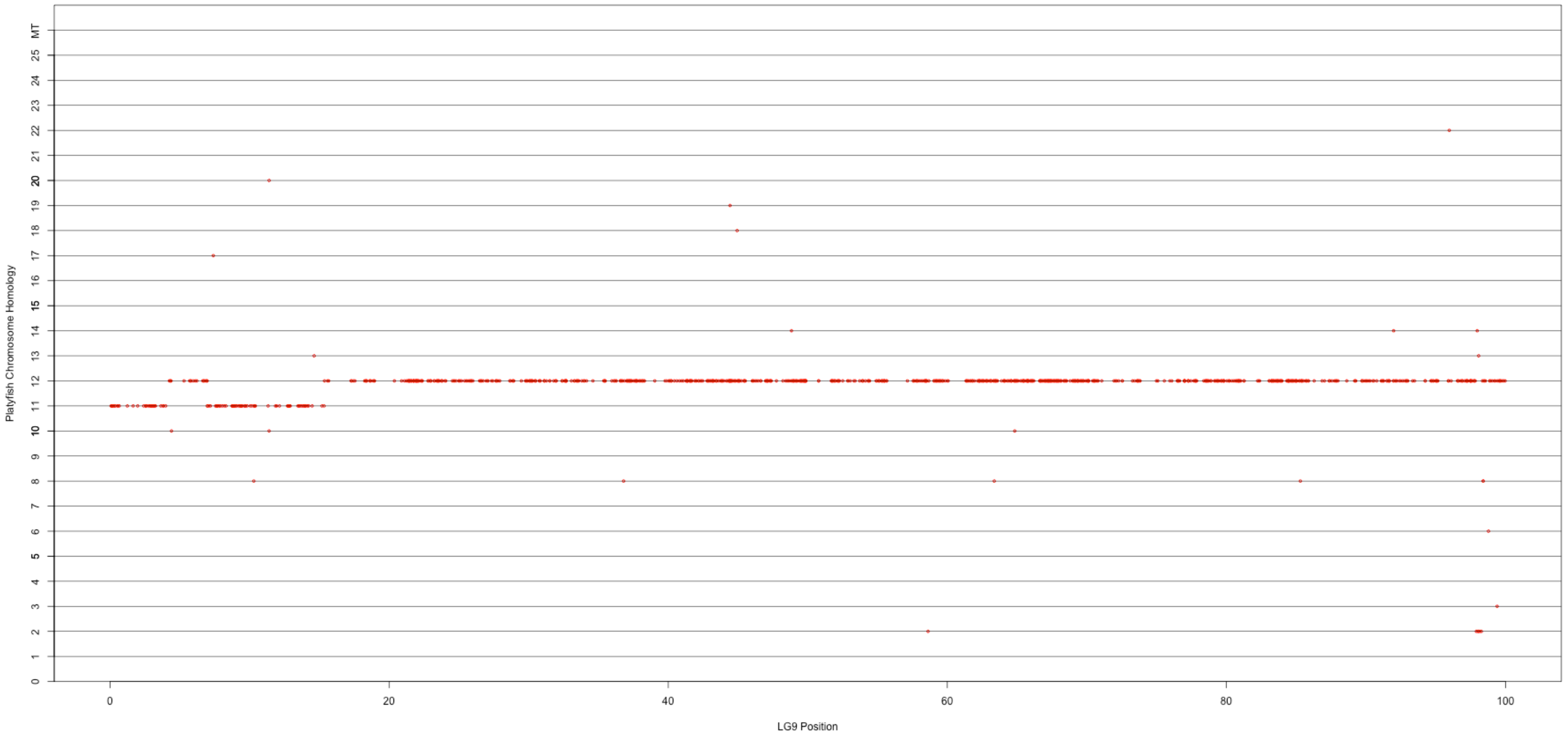

### Figure S7 (LGs 1 to 23 (7 F reversed).pdf

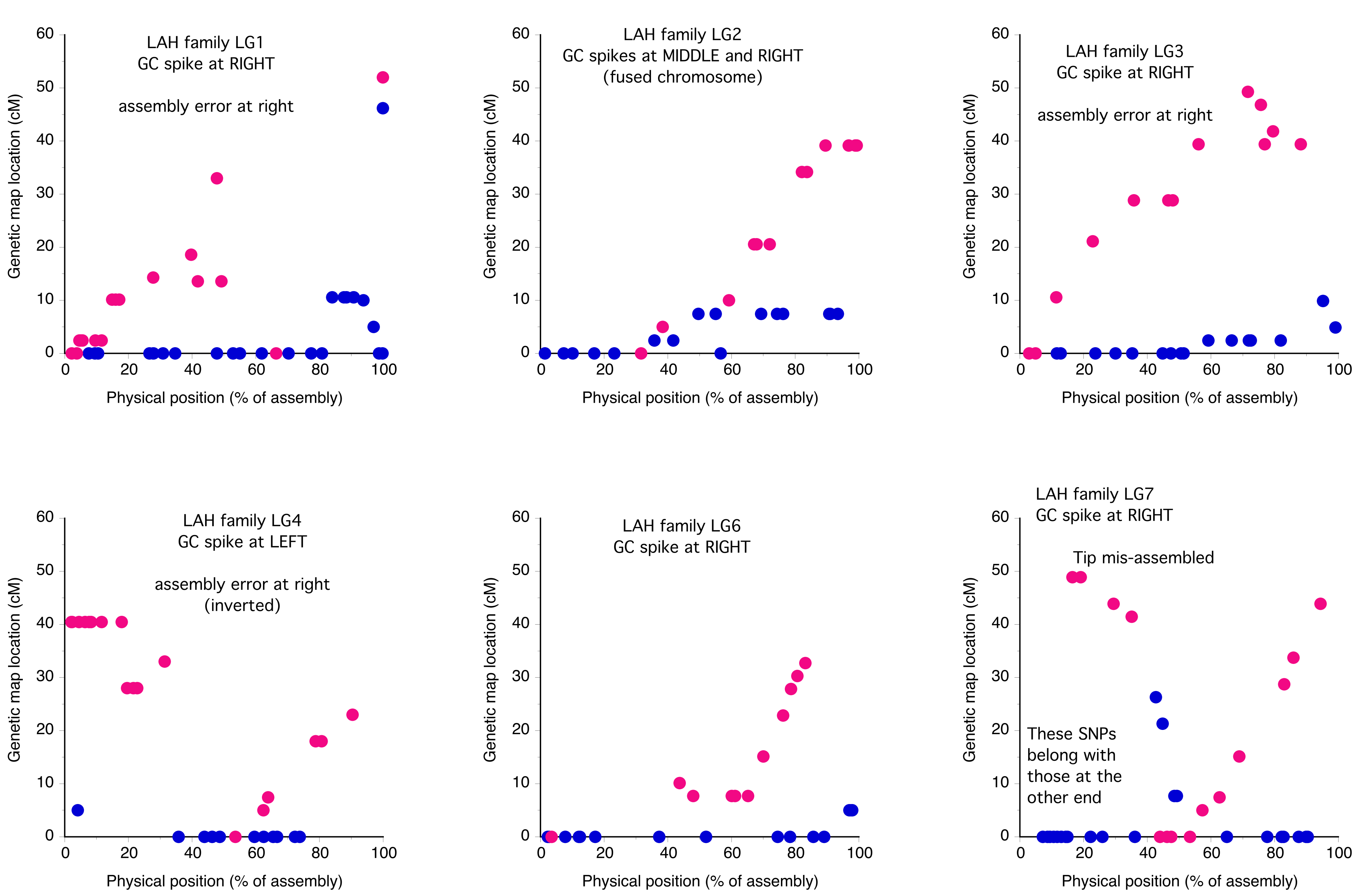

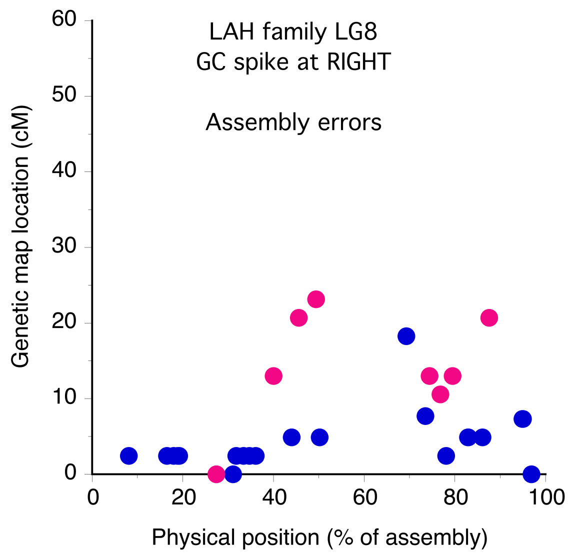

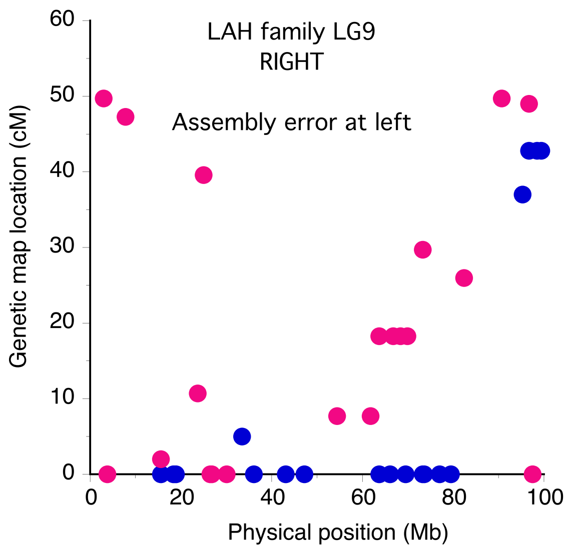

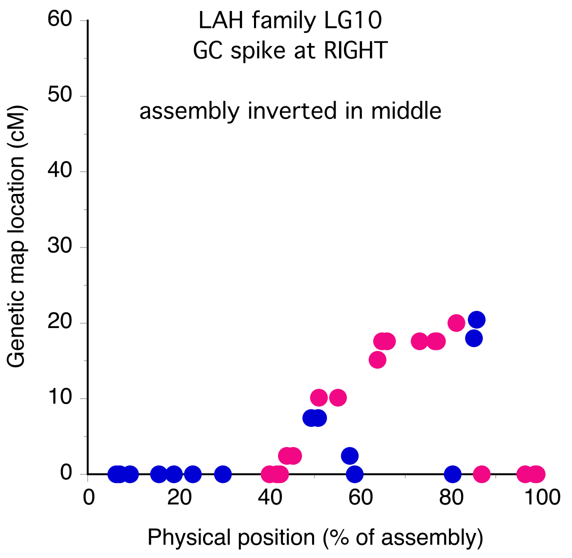

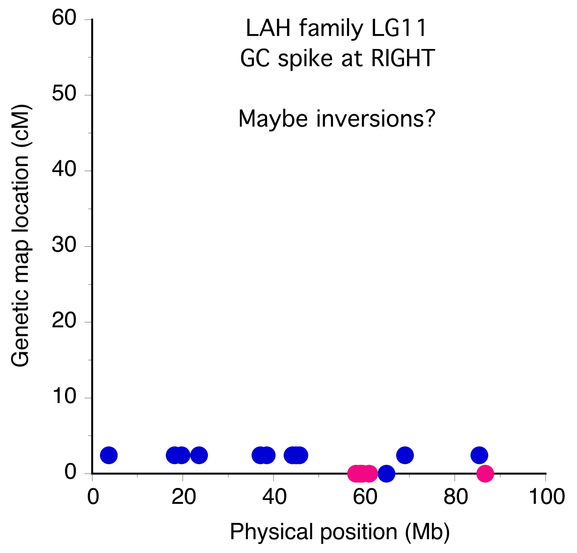

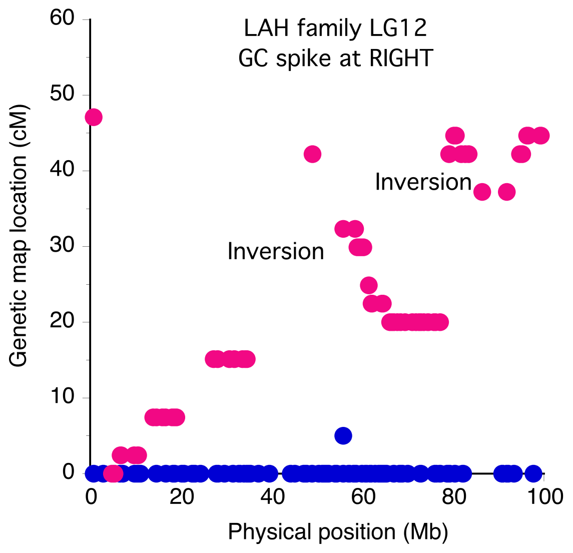

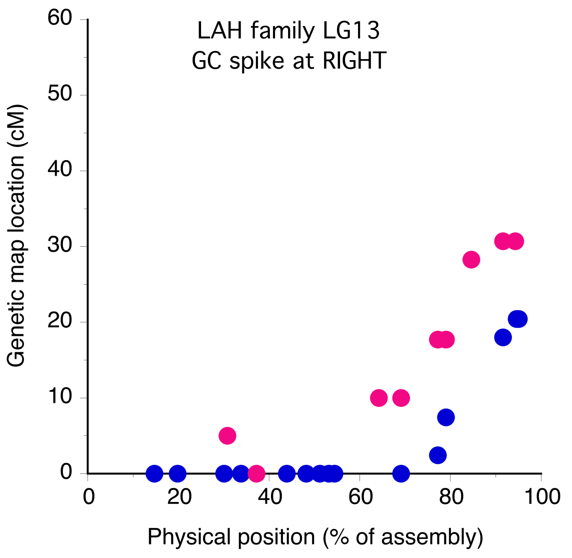

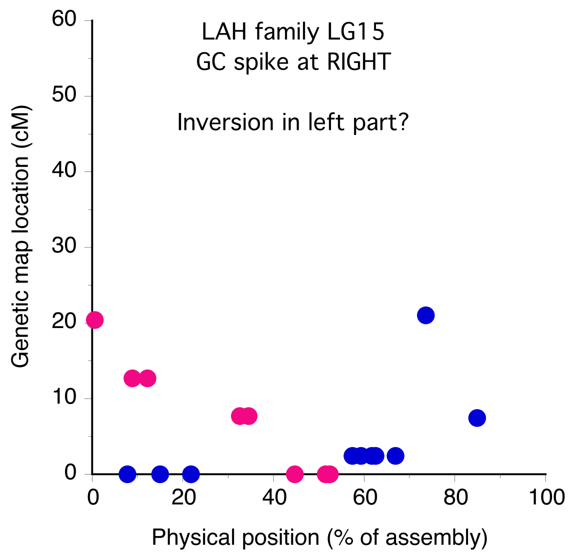

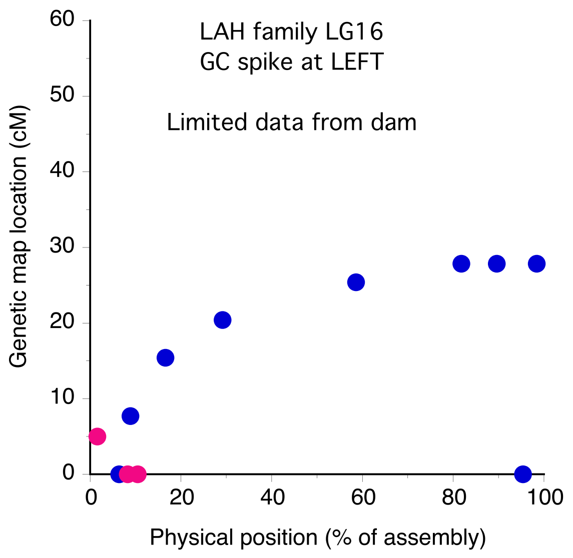

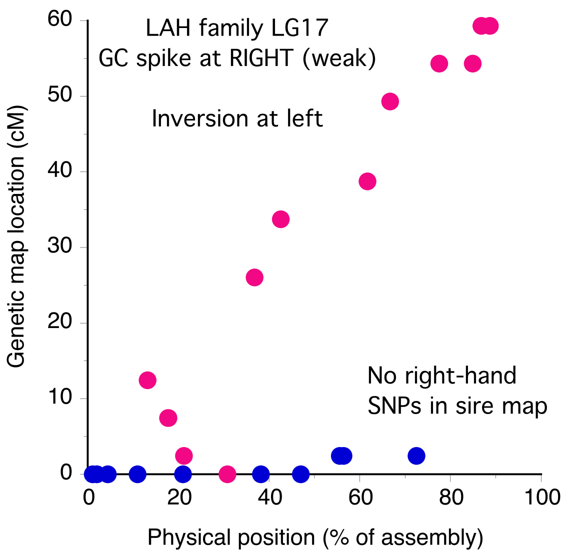

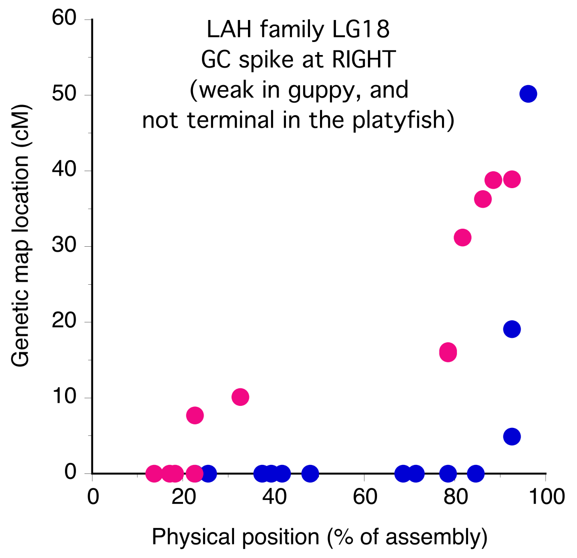

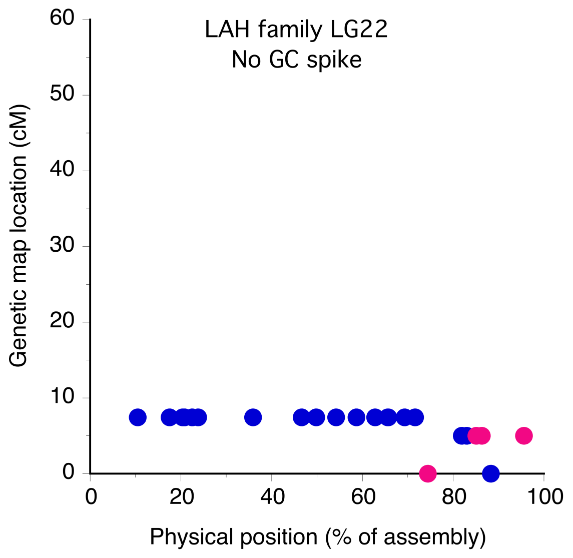

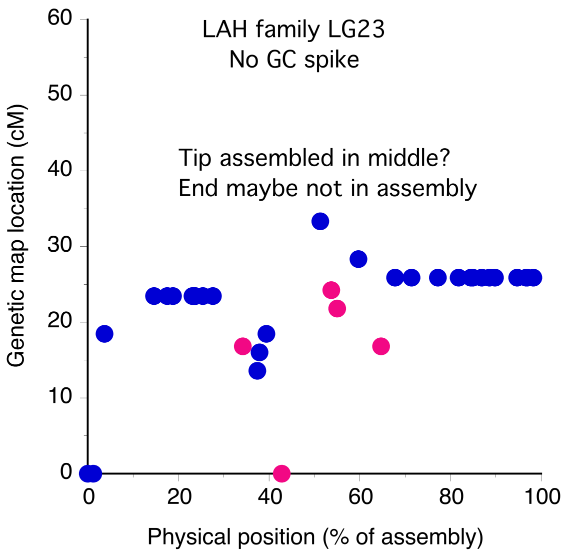
